# Rapid generation of homozygous fluorescent knock-in human cells using CRISPR/Cas9 genome editing and validation by automated imaging and digital PCR screening

**DOI:** 10.1101/2021.06.23.449557

**Authors:** Moritz Kueblbeck, Andrea Callegari, Beatriz Serrano-Solano, Jan Ellenberg

**Affiliations:** Harvard Medical School

## Abstract

We have previously described a protocol for genome engineering of mammalian cultured cancer cells with CRISPR/Cas9 to generate homozygous knock-ins of fluorescent tags into endogenous genes ^1^. Here, we are updating this protocol to reflect major improvements in the workflow regarding efficiency and throughput. In brief, we have improved our method by combining high efficiency electroporation of optimized CRISPR/Cas9 reagents, screening of single cell derived clones by automated bright field and fluorescence imaging, rapidly assessing the number of tagged alleles and potential off-targets using digital PCR (dPCR) and automated data analysis. Compared to the original protocol ^1^, our current procedure (i) significantly increases the efficiency of tag integration, (ii) automates the identification of clones derived from single cells with correct subcellular localization of the tagged protein and (iii) provides a quantitative and high throughput assay to measure the number of on- and off-target integrations with dPCR. The increased efficiency of the new procedure reduces the number of clones that need to be analysed in-depth by more than ten-fold, and yields up to 20% of homozygous clones in polyploid cancer cell lines in a single genome engineering round. Overall, we were able to dramatically reduce the hands-on time from 30 days to 10 days during the overall ∼10 weeks procedure, allowing a single person to process up to 5 genes in parallel, assuming that validated reagents – e.g. PCR-primers, dPCR-assays, Western Blot antibodies – are available.

## INTRODUCTION

### Development of the protocol

Fluorescence live-cell imaging techniques enable capturing abundance, stoichiometry and dynamics of endogenously expressed proteins or shedding light on fundamental biological processes such as cell division, ^2^ mitotic chromosome formation ^3^ or 3D genome architecture establishment ^4 5^. However, a reliable quantitative assessment requires that all alleles of the target protein of interest (POI) are tagged with the fluorescent marker. This protocol therefore focuses on the generation and the screening of homozygously tagged POI with large fluorescent or functional tags in human cell lines.

Despite the ease of designing CRISPR/Cas9 genome editing reagents to generate knock-ins at the locus of interest in cell lines and different model organisms ^6 7 8 9^, it is often necessary to screen through several hundreds of clones to identify those bearing the desired genetic modification without off-target changes. This implies culture and maintenance of many unique clones, entailing highly laborious, time-consuming and error-prone procedures that consume large stocks of disposable materials and expensive reagents. This problem becomes particularly challenging when attempting to generate homozygous knock-ins in polyploid cells, such as human cancer derived cell lines, e.g. HeLa or U-2 OS.

As tagging more than one allele happens with lower efficiency, most current approaches ^6 7 8 9 10 11 12^ generate heterozygous clones which are used for subsequent rounds of iterative CRISPR tagging until homozygosity is achieved, making the whole procedure time-consuming and prone to off-target modifications. Multiple rounds of engineering can easily lead up to one year of work before achieving homozygosity for ≥3 alleles in cancer cells.

These approaches describe very involved procedures to select knock-ins using antibiotics and removal of the antibiotic-resistance cassette ^11^ or generate knock-ins either via single-stranded oligo DNA nucleotides (ssODN) ^10^ or long, single-stranded DNAs (ssDNA) donors ^12^. However, ssODNs are more suited for single-base substitutions rather than for fluorescent proteins (FPs) or self-labelling tags (e.g. Halo-tag), and knock-in generation and ssDNAs synthesis is an expensive and laborious procedure.

This protocol introduces several improvements to achieve a high percentage of homozygous clones in a single round of engineering. In particular, we optimized (i) generation, (ii) screening and (iii) in-depth validation of knock-in cell lines.

In brief, we implemented the efficient delivery of improved CRISPR reagents and the Fluorescence Activated Cell Sorting (FACS) strategy of targeted cells. On top, we introduced automated identification of colonies derived from single clones, followed by automated imaging to determine correct sub-cellular localization of the fluorescently tagged protein. We then proceeded with the establishment of a quantitative measurement of the copy number of tag integrations by dPCR and automated computational scoring of on- and off-target integrations.

First, we optimized the delivery of CRISPR reagents into cells by electroporation, using the Neon™ electroporation system (Thermo Fisher Scientific). We migrated from lipofection to electroporation because of improved cell viability (up to 95%), which allows in turn to screen the edited clones already at days after electroporation compared to 7-10 days as previously reported ^1^. This minimizes the generation of multiple daughter cells from a unique genome-edited parental cell, providing higher genetic diversity during the screening step.

Second, we compared two different ways of delivering CRISPR reagents, which are:

i. The triple-plasmid method (PPP), where one plasmid carries the knock-in donor, a second plasmid encodes for the Cas9(D10A) and a guide-RNA (gRNA) complementary to the target sense strand and a third plasmid encoding the nickase and the second gRNA complementary to the target anti- sense strand.
ii. The donor plasmid (P) combined with ribonucleoparticles (RNP) method (PRNP), where we used in vitro pre-assembled RNPs formed by purified recombinant Cas9 protein and synthetic gRNA (sgRNA).

The PPP method has been compared to PRNP since the use of RNPs has been shown to generate a lower rate of off-target integrations ^13 14^. We targeted the Nup93 locus in HeLa Kyoto cells with mEGFP and scored the number of hetero- and homozygous clones and tested for off-target integrations using dPCR. The PRNP method generated a higher fraction of homozygous clones with ∼28% compared to PPP with ∼14% (Fig. 5D). PRNP also produced fewer off-target integrations with 50% compared to PPP with 73% (Fig. 5D).

Based on the encouraging results obtained with PRNP, we extended this approach further into the plasmid-free PCRNP method where a linear donor molecule generated by PCR is combined with the Cas9/sgRNA RNPs. The PCRNP method combines the higher efficiency and lower off-target frequency of RNP delivery with more rapid downstream selection steps provided by the use of a donor PCR product. Beside the simplicity of using a PCR product instead of a plasmid, one major advantage of introducing a donor PCR in the PCRNP method is to further decrease the waiting time between electroporation and selection from 5 to 2 days compared to PPP and PRNP due to no (<0.05%) transient fluorescent expression of the tag (Fig. 2C left panel; donor-ctrl). Furthermore, a single round of fluorescence-activated cell sorting (FACS; Fig. 2C) could be used instead of two consecutive sorting cycles with a few days interval, moving clone selection as a critical step earlier in the protocol, which avoids additional stress and artefacts by long cell culture prior to selection.

**Figure 1.**
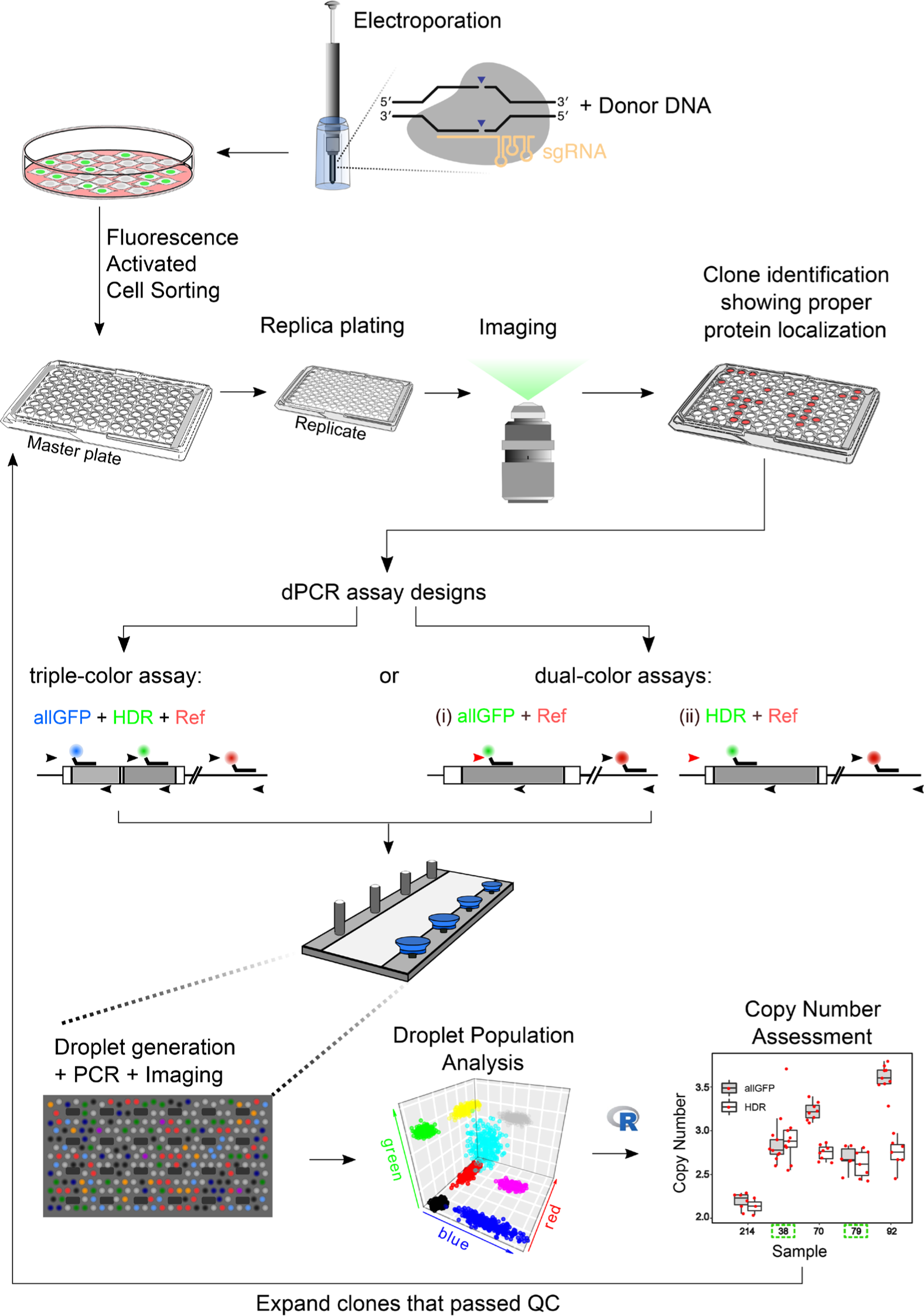
Scheme representing the main steps of the improved pipeline. After delivery of CRISPR components by electroporation, fluorescent edited cells are sorted by Fluorescence Activated Cell Sorting (FACS). Plates are taken for automated, high-throughput BF and fluorescence image acquisition. Automatic composition and rendering of acquired images quickly enables a visual identification of positive clones that are analysed by dPCR using a dual- or triple-colour assay. Using a home-made R analysis software, dPCR results are used for quantitative assessment of tagged copy number and off-target. Clones which passed the QC are highlighted with a green dashed square; namely clones #38 and #79 in this example.

**Figure 2.**
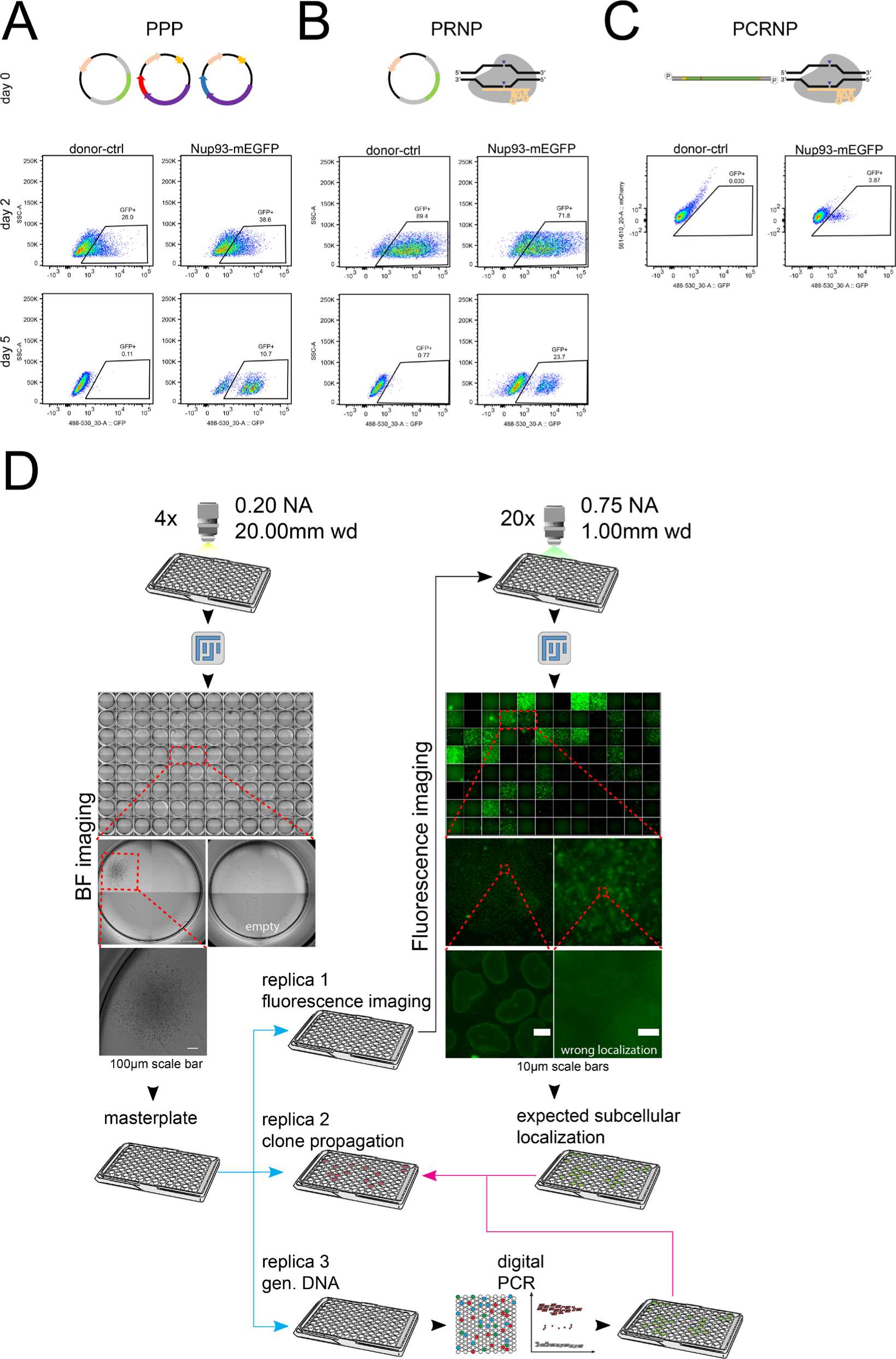
Delivery of CRISPR reagents by electroporation, followed by dual FACS sorting. (A-C, upper panels) Population enrichment is performed 48h post-electroporation. mEGFP+ cells were collected and sorted together in standard 6-well culture plates. Gating of fluorescent channels was empirically determined based on wild-type, unmodified cells. ∼38-72% of electroporated cells were sorted (black frame). Remarkably, control experiments run in parallel by electroporating the donor plasmid alone (donor-ctrl) resulted in a significant proportion of mEGFP expressing cells (28-89%, depending on the CRISPR strategy). (A-B, lower panels) Single-cell sorting of enriched cell populations. After five additional days, single cell sorting of mEGFP^+^ cells (black frame; ∼39%-72%) was performed in 96-well plates. (C) FACS analysis of cells electroporated with PCR-donor and preassembled RNPs (PCRNP strategy). (left panel) delivery of PCR-donor alone produced <0.05% of mEGFP^+^ cells. (right panel) The combination of the PCR-donor with RNPs resulted in ∼4% of mEGFP-expressing cells that were directly sorted in a 96-well plates. (D) Automated imaging of FACS-sorted clones. (left panel) Bright-field (BF) and (centre panel) fluorescent wide-field (WF) imaging of 96-well plates containing individual growing clones. (4x) objective was used to image each well for BF, with 0.20 Numerical Aperture (NA) and 20.00mm working distance (wd) and (20x) objective was used to image each well for WF, with 0.75 Numerical Aperture (NA) and 1.00mm working distance (wd). Image reconstruction was performed with a Fiji plugin developed for this specific purpose. (bottom panels) Representative wells containing single colonies (left panel) or fluorescent cells (centre panel). Empty wells or clones displaying mis-localized fluorescent protein were discontinued.

After single cell FACS sorting into 96-well plates, we have also implemented an automated bright-field (BF) imaging step to verify colonies truly derived from single cells directly imaging the original plastic plates. This is a basic but important step to avoid propagation of non-monoclonal populations resulting from errors in single-cell FACS sorting. Microscope imaging was optimized to allow quick interactive inspection of growing colonies on the computer screen using a dedicated plugin for the open source image analysis software Fiji (Fig. 2D, left panel). The same automated microscope was used to collect wide-field (WF) fluorescence images to identify and propagate only clones showing the expected subcellular localization of the tagged protein (Fig. 2D, centre panel). Overall, this procedure enables fast and reproducible identification of single-cell derived colonies with the correct subcellular localization of the tagged protein.

In our improved protocol, positive clones verified by BF and fluorescence microscopy were next analysed by digital PCR to measure the number of tagged alleles as well as off-target tag integration elsewhere in the genome. To this end we have developed a triple-colour dPCR assay that allowed performing both target assays simultaneously in a single dPCR run, which simplifies and accelerates the screening of clones. Using a computational routine implemented in the open source statistical analysis software R, the collected dPCR datasets were automatically processed to rapidly quantify tagged allele copy numbers and assess off-target integrations. This information enabled the rapid and reliable identification of hetero- and homozygous clones, allowing us to weed out clones with off-target events at an early step of the pipeline and propagate only properly engineered clones, thereby saving time and material consumption.

Finally, we proceeded with an analysis of the expression of the tagged proteins using capillary electrophoresis, instead of the previously used traditional Western blot. With this simplified assay, protein sample preparation and capillary electrophoresis can be performed within 4 hours, automatically generating quantitative protein expression data. This is a technique of choice due to the simplicity of usage, the speed, the quantitative results and because it also outperforms conventional 2D gel electrophoresis and Western blotting when working with large proteins and small kDa tag size shifts.

An overall summary scheme of the improved screening protocol is detailed in Fig. 1A scheme of the entire pipeline, including time required for design, delivery, screening and in-depth-analysis, is shown in Sup. Fig. 1.

### Comparison with our previous CRISPR editing protocol

In our previous protocol ^1^, we established a CRISPR pipeline which had ∼0.5-1.5% overall editing efficiency, considering heterozygous and homozygous clones together. The low efficiency was mainly due to (i) limited efficiency of the transfection procedure and comparatively high associated cell death, (ii) long waiting time between transfection of CRISPR reagents and single cell sorting by FACS, diminishing genetic diversity due to clonal cell proliferation, (iii) high frequency and late detection of off-target integration that led to late dropping out of many of the propagated clones and (iv) limited throughput and non-quantitative screening to identify the correctly edited clones.

Compared to our previous protocol ^1^, we opted to deliver endotoxin-free plasmids using electroporation. This strategy enhanced delivery efficiency up to ∼35%, maintaining 80-95% of cell viability when tested in HeLa Kyoto cells with 3 fluorescent expressing plasmids simultaneously (PPP method, data not shown).

To improve CRISPR specificity and avoid extra integrations due to prolonged expression of Cas9 ^15 13^ (see also Fig. 3 and Fig. 5), we used RNPs with purified recombinant Cas9 in combination with a donor plasmid to limit Cas9 activity within ∼12h post transfection^16^. In comparison with our former approach, the RNPs method gave a higher rate of targeted alleles and lower rate of off-target events (Fig. 5D). We then developed this approach further by replacing the donor plasmid by a linear PCR-amplified DNA donor. This is a major improvement compared to our previous pipeline since this (i) simplifies the production of the donor reagent, since it is much shorter and uses a relatively low cost gBlock as template (ii) allows to perform a single FACS round instead of two and (iii) allows to sort cells 2 days after transfection instead of 5. Moreover, we have taken advantage of the simple modification of the donor reagent in the PCRNP approach to test the effect of reducing the size of the homology arms. The use of shorter homology arms without reducing gene targeting efficiency then allowed us to develop a single step triple-colour dPCR assay for on-target copy number measurement and off-target detection. On top the usage of a PCR product as a donor allows modifying the ends, which has been reported to generate higher Homology-Directed Repair (HDR) rates ^17^.

**Figure 3.**
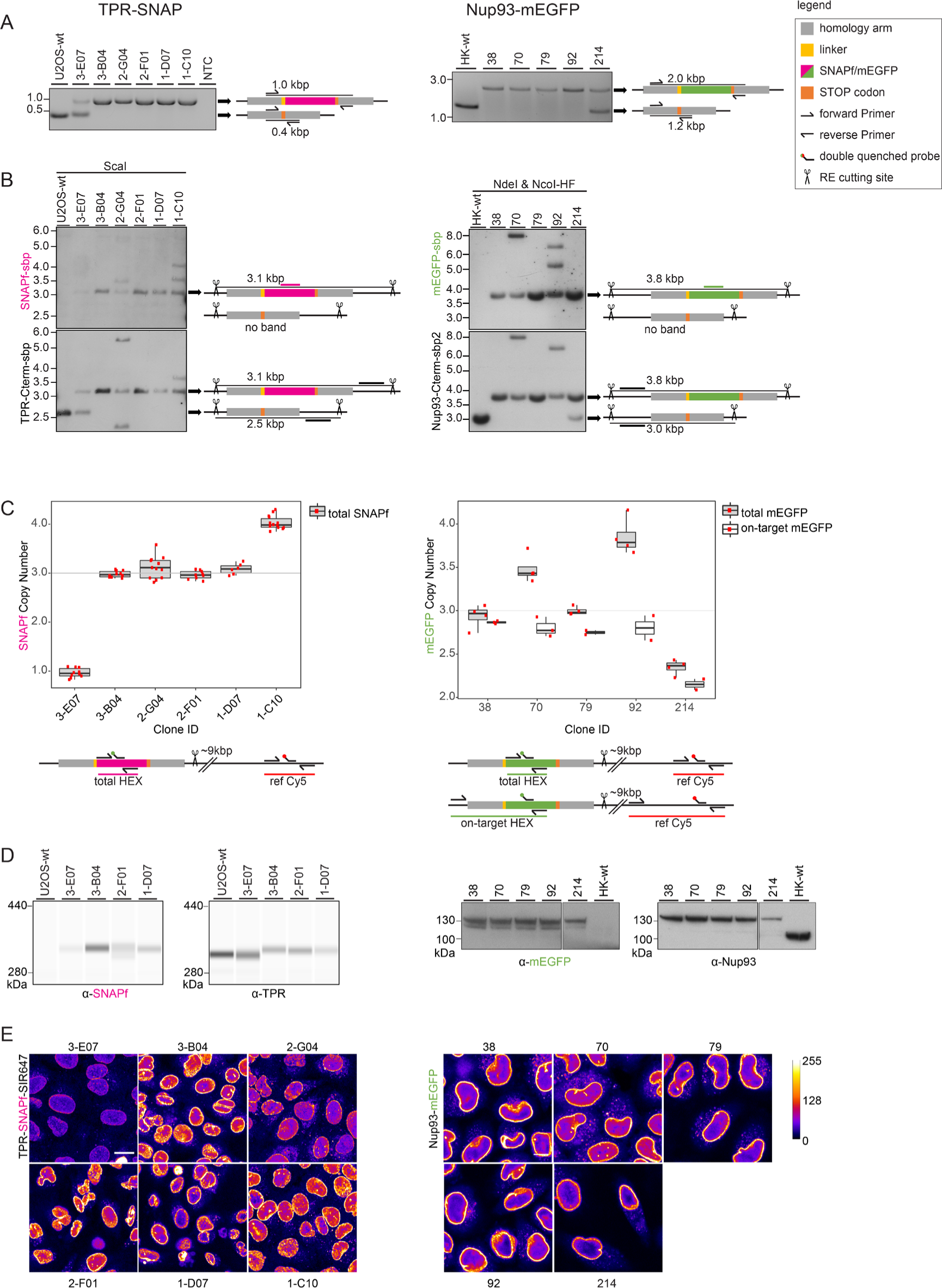
In-depth validation of genome edited U2OS-TPR-SNAP and HK-Nup93-mEGFP clones. (A) Standard-PCR performed at the target integration site. Representative clones are shown with their corresponding bands. The upper band indicates a successful integration of the tag (SNAP or mEGFP). (B) Southern Blot performed genome-wide. Representative clones are shown with their corresponding bands. A band at 3.1 kbp for TPR-SNAP or a 3.8 kbp band for Nup93-mEGFP indicates a correctly tagged allele detected with their corresponding tag specific as well as endogenous probes. A band at 2.5 kbp for TPR- SNAP or a 3.0 kbp band for Nup93-mEGFP indicates a wild-type allele detected with their corresponding endogenous probes. Any other bands detected with both probes indicate extra integrations or major rearrangements of the genome. (C) dPCR performed genome-wide. Representative clones are shown with their corresponding box-plots. Every red dot represents an individual measurement. Technical replicates for U2OS-TPR-SNAP and biological replicates for HK-Nup93-mEGFP. Total number of tag integration for both cell lines and on-target tag integrations for HK-Nup93-mEGFP cell line. (D) Capillary electrophoresis for U2OS-TPR-SNAP, the « Lane View» setting has been chosen to display the results; Western Blot for HK- Nup93-mEGFP. (E) Confocal fluorescence imaging.

Differently from our previous protocol ^1^, a single round of cell sorting at 48h post-transfection (Fig. 2C) enabled us to minimize the contribution of clonal cell expansion deriving from the same unique parental cell genome editing event. Importantly, this novel approach maximizes genetic diversity among the sorted clones. Also, transient expression of the tag that can result from the donor plasmid alone has been experimentally documented ^18^ and reproduced in our laboratory (Fig. 2A and 2B). As compared to our published protocol ^1^, we optimized the time between electroporation and cell sorting to reduce residual donor expression levels in the recipient cell line, which resulted in waiting ∼5 days post-electroporation (Fig. 2A and 2B). At this point, we then added a second FACS step to enrich again single fluorescent cells for the following selection steps. This strategy can still be used, but we now highly recommend the PCRNP approach to avoid clonal expansion prior to cell sorting, shorten selection time, and to reduce stress on cells due to successive round of expansion culture and FACS.

The screening of growing clones has been now updated using an imaging-based, automated screening step that relies on an in-house developed Fiji plugin ^19^ complemented with high-throughput, automatic fluorescence microscopy to exclude clones with mis-localized tagged proteins.

In the previous version of the protocol ^1^, we discriminated between homo- and heterozygous clones using conventional PCR, which failed to detect off-target integrations since it only interrogated the locus of interest. This approach required a subsequent Southern Blot as a gold-standard method to detect off-target integrations of the donor. However, Southern Blot is a laborious and low-throughput technique that requires comparatively large amounts of genomic DNA and therefore cell material, delaying the overall procedure. In the improved protocol presented here, we introduced dPCR to quantitatively characterize on- and off-target genome-editing events. We show that dPCR can assess total and on-target tag integrations and can be performed with small amounts of cell material. Additionally, we have improved the dPCR screening throughput by implementing a one-step triple-colour assay. In this assay, we quantitatively performed copy-number assessment of tagged alleles in a robust and reproducible way and simultaneously probed for off-target events in a single dPCR run. Since the triple-colour assay requires only small number of cells, it can be performed at an early step of the overall genome editing pipeline and because dPCR requires only ∼3-4 hours, the culture of incorrectly edited clones can be discontinued on the same day.

We have also furthermore used capillary electrophoresis (Jess, Bio-Techne®) to quantitatively assess the expression of the tagged protein. Protein sample preparation and electrophoresis are completed within 4 hours and automatically generate quantitative data, allowing computational scoring of clones. Capillary electrophoresis can be done with a large variety of protein sizes and small shifts in mobility due to tag integration can be detected even for large proteins. In our case, we could for example reliably detect the 20 kDa shift produced by the SNAP tag on the 267 kDa TPR protein (Fig. 3D, left panel, anti-TPR), which is very difficult with conventional Western blotting ^1^.

With our improved protocol, once all reagents are available, generation, screening and in-depth analysis of selected clones takes ∼10 weeks to be completed (Supplementary Fig. 1). Compared to the previous protocol, this represents a significant improvement in start-to-finish time as well as hands-on-time during the procedure, allowing to produce fully validated cell lines faster and handling more target genes in parallel. We have moved several diagnosis steps earlier in the pipeline to reduce the number of clones to handle and dedicate human and material resources to promising clones only. The previous version of the protocol typically required several editing rounds to tag all alleles successfully. Consequently, this could lead to a year of work before obtaining a homozygous clone. Here, we describe a method which generates high percentages of homozygous clones in a single editing round after 2-3 months.

Taken together, the improved CRISPR reagents and their delivery together with the combination of automated imaging and dPCR-based screening allowed us to perform tag copy-number and off-target assessment in a robust and reproducible way. With this protocol, we achieved homozygous-tagging efficiencies of around 20% in a triploid locus of HeLa Kyoto and in a single editing round, resulting in fully validated cell lines after only two months.

## EXPERIMENTAL DESIGN

### gRNA / donor design

gRNAs sequences were designed either using the CRISPR design tool implemented in Benchling (https://www.benchling.com/crispr/) or the Integrated DNA Technologies (IDT) design tool (https://eu.idtdna.com/site/order/designtool/index/CRISPR_CUSTOM).

These CRISPR gRNA design tools allow users to visualize, optimize, and annotate multiple gRNA sequences at a time. They also predict and score on- and off-target mutation rates and optimize guides for higher activity and lower off-target effects. Even if Benchling/IDT software provides on- and off-target specificity of a selected sgRNA, the algorithms are not perfect predictors and we recommend selecting multiple sgRNAs for one engineering experiment to maximize the chances of achieving genome editing. Additionally, the efficiency of DNA cutting varies substantially and depends on the type of cells used, the transfection method, the target sequence, etc. To account for these factors, the selected sgRNAs should be tested before use, for example with a T7 Endonuclease-I assay to score for their relative cutting efficiencies. We describe here the design of sgRNAs for the C-terminal tagging of Nup93 with mEGFP in HeLa Kyoto cells and of TPR with SNAP in U-2 OS cells. The same criteria can be used for N-terminal tagging. The input sequence used to feed the designing-tools for sgRNA design spans ∼50 bp up- and downstream of the target STOP codon. The «paired guides» option is chosen when using Cas9 (D10A) «nickase» mutant; otherwise, the «single guide» option for wild-type Cas9 is selected. Importantly, the location of genome editing must be carefully considered. First, sgRNAs should be chosen to produce nicks as close as possible to the insertion site ^20^. Ideally, the cutting site should be less than 10 bp away from the knock-in site, because double-strand breaks (DSBs) or nicks that are closer to the mutation site typically result in higher levels of HDR. Moreover, whenever possible, it is strongly advisable to pick one sgRNAs producing nicks within the START or the STOP codon. In fact, such sgRNA will produce a cut exclusively on the target genome sequence without affecting the donor DNA that should stay intact. If cutting does not occur within START or STOP codons, we recommend favouring sgRNAs targeting a non-coding region of the selected gene to avoid altering the protein-coding sequence. Besides sgRNAs design, we introduced a standardized cloning procedure using Golden Gate to minimize design and cloning steps required to insert sgRNAs sequences in the pX335 receiving plasmid. Specifically, a sgRNA-expression vector can be obtained by cloning the 20-bp target sgRNA sequence into pX335, which encodes a human U6 promoter-driven sgRNA expression cassette and a CBh-driven Cas9-D10A nickase (pX335-U6- Chimeric_BB-CBh-hSpCas9n(D10A); Addgene plasmid #42335). The insert containing the sgRNA sequence is obtained by purchasing two complementary oligonucleotides from IDT or from Sigma-Aldrich as Reverse Phase (RP)- or HPLC-purified DNA oligonucleotides. To standardize cloning, all sgRNA sequences are flanked by the same short stretch of nucleotides and when annealed, the resulting product carries BpiI- compatible overhangs which serve to clone into pX335 vector as previously described ^21^. Note that the Protospacer Associated Motif (PAM) sequence required for target recognition by the Cas-nuclease is never present as part of the sgRNA itself. Unlike in our previous version of this protocol, cloning of sgRNAs was carried out by using the Golden Gate protocol ^22^ (see also: https://international.neb.com/golden-gate/golden-gate). This systematic procedure abolished the need to design cloning strategies and insures 100% cloning efficiency. After amplification, it is highly recommended to purify the guide-expression plasmid with an endotoxin-free plasmid maxiprep kit (e.g. Qiagen) before its use for electroporation.

Additionally, we tested a combination of an HDR donor plasmid and RNPs generated in vitro by complexing chemically modified sgRNAs with purified recombinant SpCas9 to target the Nup93 locus in HeLa Kyoto cells. Besides the advantage of avoiding cloning procedures, the RNP-based CRISPR format limits the Cas9-dependent cutting activity to up to ∼48 hours ^23^, minimizing the likelihood to generate off-target integrations of the tag. Synthetic chemically modified sgRNAs were purchased from Thermo Fisher Scientific (TrueGuide Synthetic gRNA). They are synthesized as 100-mer sgRNAs that incorporate 2’-O-methyl analogues and 3’-phosphorothioate internucleotide linkages in the terminal three nucleotides on 5’ and 3’ ends of the sgRNA. These modifications enhance the editing efficiency by increasing binding to the target site and inhibiting nuclease degradation, respectively ^24 25^. For the PCRNP method we ordered modified sgRNAs (Alt-R® CRISPR-Cas9 sgRNA) from IDT.

Donor plasmids include the nucleotide sequence of the tag. In this study, we either used the fluorescent reporter mEGFP or the self-catalytic SNAP to tag human Nup93 in HeLa Kyoto or TPR in U-2 OS, respectively. To minimize the risk of sterically hindering the protein function, we recommend the addition of a flexible linker to connect the fluorescent tag to the N- or C-terminus of the target protein. Whenever possible, we encourage the reader to use linker sequences that have been already documented for the protein of interest. In the current protocol, we used HDR donor plasmids designed according to the criteria illustrated in our previous report ^1^. Briefly, we included ∼800 bp homology arms on both sides of the tag sequence. We chose homology regions using the specific cell line genomes, obtained from our internal databases (http://bluegecko/cgi-bin/HeLa_genome.pl) or (http://bluegecko/cgi-bin/U2OS_genome.pl). If these specific genomes are not retrievable, one can use homology regions from ENSEMBL (https://www.ensembl.org) using the human genomic database as a direct source. Another option is a locus Sanger Sequencing to obtain the specific cell line sequences. Importantly, we initially assumed that HDR donor plasmids including homology regions of >500 bp in length could maximize the rate of HDR, as previously reported by us ^1^ and others ^26^. If the sgRNA target region is exclusively within the homology arm, and doesn’t span over the START or STOP codon, silent mutations of >1 bp in the donor are recommended, to avoid donor recognition and cutting ^27^.

Motivated by the ease and low-cost of propagating and retrieving plasmid DNAs, we first explored the possibility to enhance the efficiency of our CRISPR pipeline using the PPP method. Our optimized procedure generated ∼13.5% of homozygous clones (Fig. 5D). It is noteworthy that this efficiency rate cannot be directly compared with those published in our previous report. In fact, in our current protocol, we shifted FACS sorting at an earlier step, that is, 48 hours post-electroporation and under these new conditions, we observed a significant proportion of cells expressing either free mEGFP or SNAP from the donor plasmids. To overcome this problem, we established the minimum waiting time to obtain a negligible fraction of transiently-expressing cells. We performed a time-course of transient expression of the donor plasmid alone, assessing the number of fluorescent cells from 24h up to 6 days after electroporation (data not shown). We concluded that 5 days were enough to neglect the contribution of donors transient expression, enabling us to shorten the overall cell-line generation time compared to our previous protocol.

Importantly, recent findings ^28^ suggest that homology regions <100 bp can be successfully used to generate large knock-ins in human cells and other model organisms with HDR efficiencies comparable to those achieved with long homology regions. This is a relevant aspect that should be considered when designing a strategy of CRISPR screening based on dPCR. In fact, dPCR performs best with amplicons < 125 bp. The dPCR assessment of the on-target integrations necessarily needs a primer annealing outside of the homology region and a different primer pairing within the tag sequence (Fig. 1). Homology arms of ∼800 bp would lead to amplicons well beyond the optimal length suggested for the optimal performance of dPCR assays, even though guidelines for long homology regions have been described ^29^.

The transient expression of the tag leaking from the donor plasmids combined with the need of implementing a quantitative CRISPR screening strategy based on dPCR triggered us to test a novel CRISPR design. We generated a linear donor DNA tailored to include two 40 bp-long homology regions (Fig. 4A). Moreover, in order to generate two independent amplicons for the dPCR triple-colour assay, a silent point mutation within the mEGFP ORF was introduced to create a novel MseI-restriction site. This donor DNA was successfully tested in a proof-of-principle experiment where we assessed on-target and total tag copy-number using the same reference assay in a unique triple-colour dPCR assay (Fig. 1). This novel strategy fully solves several issues at once. First, the leaky expression of tags by donor plasmids is no longer an issue since the linear donor does not show significant tag expression (Fig. 2C, <0.05%). Second, the use of 40 bp homology arms is optimal for dPCR performance because of efficient product amplification of small templates (Fig. 4C) ^30^. Third, a single FACS round can be performed 48h after electroporation, enabling an early selection of fluorescent clones. Fourth, the triple-colour assay increases the throughput of the CRISPR screening, and shortens the overall cell-line generation time.

**Figure 4.**
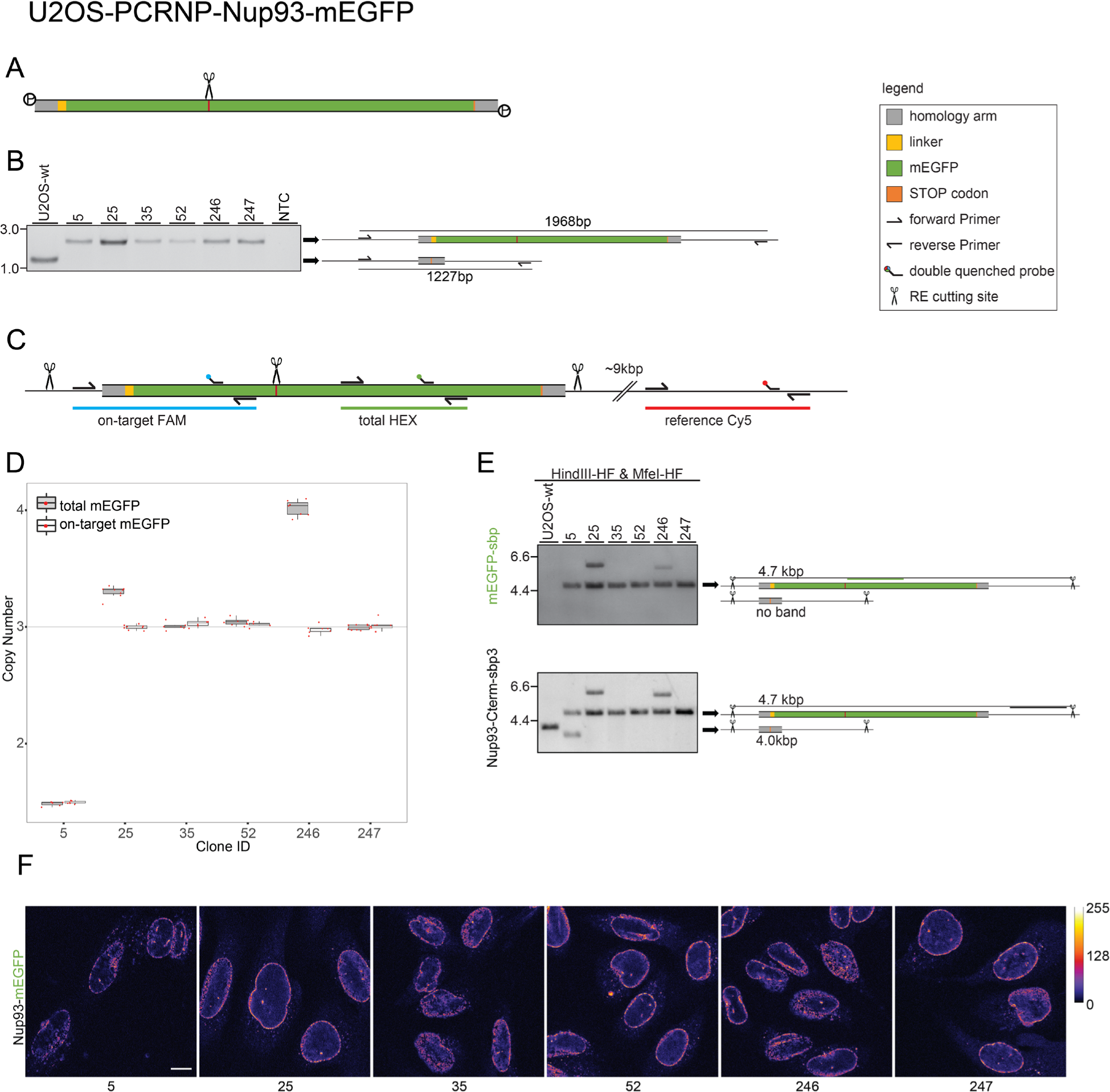
In-depth validation of genome edited U2OS-Nup93-mEGFP clones. (A) cartoon of the donor. (B) Standard-PCR performed at the target integration site. Representative clones are shown with their corresponding bands. The upper band indicates a successful integration of the tag (mEGFP). (C) dPCR triple assay design. (D) dPCR performed genome-wide. Representative clones are shown with their corresponding box-plots. Every red dot represents an individual measurement. Technical replicates for U2OS-Nup93-mEGFP. Total number of tag integrations and on-target tag integrations in the U2OS-Nup93- mEGFP cell line. (E) Southern Blot performed genome-wide. Representative clones are shown with their corresponding bands. A band at 4.7 kbp for Nup93-mEGFP indicates a correctly tagged allele detected with their corresponding tag specific as well as endogenous probes. A band at 4.0 kbp band for Nup93- mEGFP indicates a wild-type allele detected with their corresponding endogenous probes. Any other bands detected with both probes indicate extra integrations or major rearrangements of the genome. (F) Confocal fluorescence imaging.

**Figure 5.**
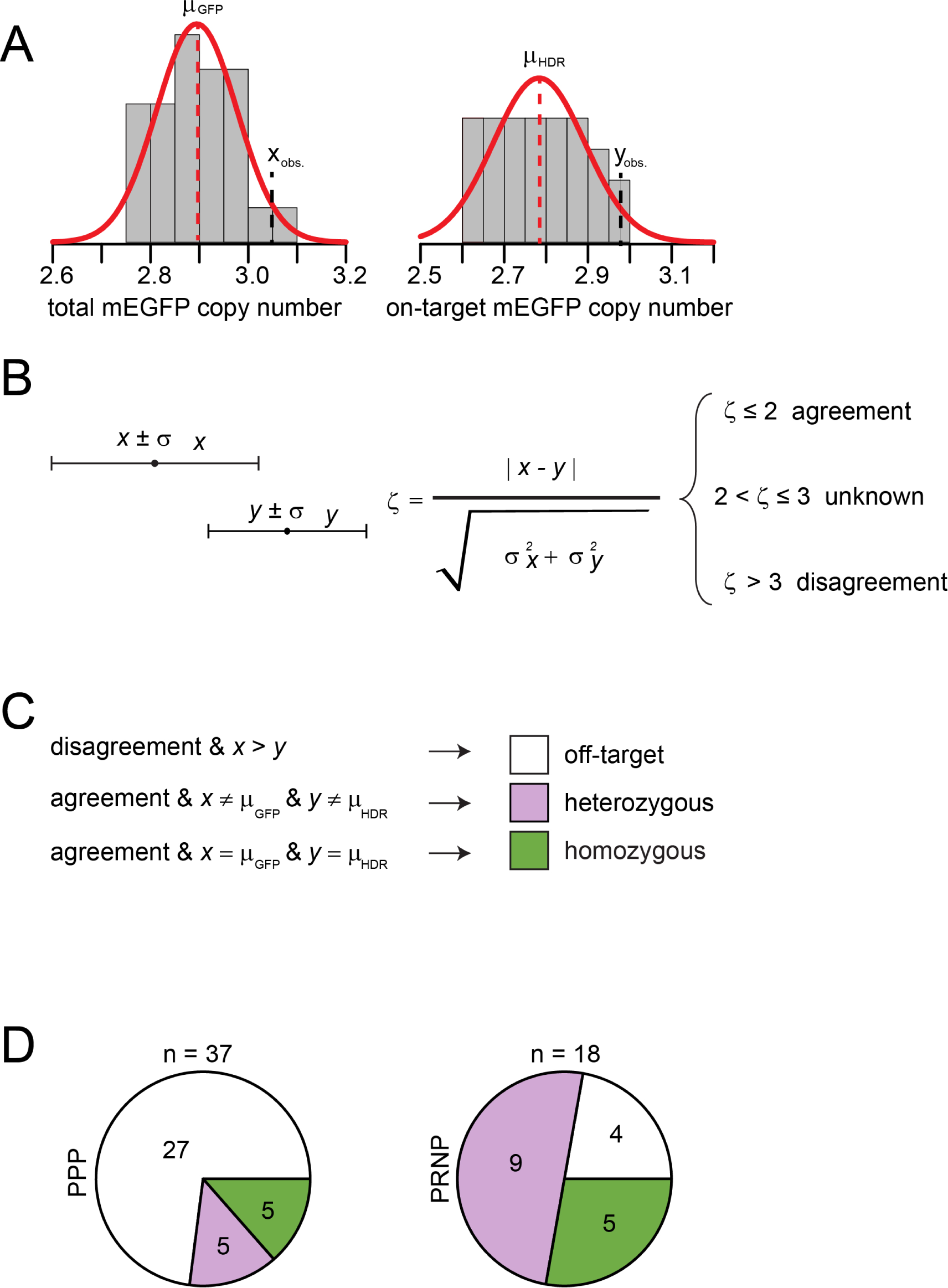
Statistical classification of genome edited HK-Nup93-mEGFP clones as a screening tool. (A) Measured distributions of a pre-validated reference clone (#38) for allGFP and HDR dPCR-assays. (B) Application of a double-sided t-test. (C) Classification criteria to determine homozygous and heterozygous clones. (D) Pie-chart of classified, edited clones generated by two different CRISPR strategies «PPP» vs.

### Electroporation

The delivery strategy of the CRISPR reagents is a critical step that affects the overall editing efficiency. Special care must be taken when designing the electroporation experiment to minimize cell stress and maximize cell survival and delivery rates. In our experience, a preliminary optimization of the delivery parameters represented the best practice. For example, since HeLa batches may vary considerably across different laboratories, we opted to establish the best electroporation conditions for our specific HeLa Kyoto cells. To this purpose, we run electroporation through 24 different conditions and measured the corresponding survival rate and delivery rate 48h after electroporation. The optimization run was repeated twice, according to the specific type of CRISPR reagents used (i.e. either plasmids or RNPs in combination with plasmids). As expected, we noticed that the best electroporation parameters empirically established significantly differed from those reported for the generic HeLa cells ^31^. Moreover, as previously documented ^32^, CRISPR reagents type affected delivery conditions (see protocol details).

### dPCR assay design

dPCR enables quantification of the amount of a given DNA sequence in a reaction mixture. In this protocol, we implemented dPCR to quantitatively assess the number of tag copies integrated by CRISPR in edited genomes. First, using existing cell lines which had been validated by conventional methods, we performed dPCR in order to assess the validity of the technique to predict genotypes. For this, we combined the information retrieved from two dPCR assays; the «on-target assay» that scores specific integrations at the locus of interest and the «total-tag assay» that measures the overall number of integrations in the whole genome.

In the «on-target assay», one primer anneals within the fluorescent tag sequence and the second on a position located outside of the homology region of the target gene. In the «total-tag assay», both primers bind within the tag sequence (Fig. 1; dPCR assay designs).

As previously documented ^33^, both assays were normalized to a similar-length reference situated about 9- 10kb away from the CRISPR target site on the same chromosome. Assays have been carried out independently (dual-colour assay; Fig. 1) or in combination (triple-colour assay; Fig. 1). Following the validation of our approach, we applied the dPCR assays to screen many CRISPR-engineered polyploid clones, generated either by PPP or PRNP, classified their genotypes as either homo- or heterozygous for the tagged alleles and assessed the number of off-target integrations.

An essential detail in the design of dPCR assay concerns the size of amplicons. Even if the optimal size is ∼125 bp, products of 75–200 bp in length can be used. Importantly, short PCR products are typically amplified with higher efficiency than longer ones, but a PCR product should be at least 75 bp long to allow discrimination from any primer-dimers that may form in the assay. Genomic regions suspected to produce secondary structures should be avoided as much as possible, as well as those presenting long (> 4) repeats of single bases. Moreover, as a general recommendation, regions with a GC content of 50–60% should be chosen.

We designed dPCR assays using the Primer3Plus tool freely available at http://www.bioinformatics.nl/cgi-bin/primer3plus/primer3plus.cgi. Briefly, after retrieving the input sequence, the user can opt to design de novo forward (fw), reverse (rv) and internal probe (int) oligonucleotides or to recycle well-established primers or probes to retrieve a new dPCR assay. For dPCR on-target assays spanning through one of the two homology arms, it is essential to exclude these regions as templates for the primer design. Primers should ideally be ∼20 bp in length with a melting temperature of ∼60°C and a GC-content of 20-60%. Importantly, internal probe oligonucleotides must have a melting temperature 3-10°C higher than primers while sharing their same optimal length and GC-content. General settings in Primer3Plus require also to specify the dPCR reaction conditions, such as concentrations of mono-/divalent cations, deoxynucleotide triphosphates (dNTPs) and primers. Moreover, the computation of thermodynamic parameters and salt correction should be performed by specifying Santa Lucia’s parameters and formulas algorithm ^34^. Amongst the several dPCR assays proposed, we picked the first one in the ranked output list.

For the triple-colour assay, we recommend using the IDT dPCR design tool (https://eu.idtdna.com/Primerquest/Home/Index).

Afterwards, we tested the optimal annealing temperature that maximized the amplification yield by running a standard PCR using a gradient of different annealing temperatures. We show a representative result in Supplementary Fig. 2A.

More importantly, we determined the optimal annealing temperature that allowed for best separation of fluorescent and non-fluorescent droplets by running dPCR reactions over a gradient of annealing temperatures and detecting the resulting separation between the two droplet populations (Supplementary Fig. 2C). This optimization step is critical to set the performance of a dPCR assay and retrieve a reliable measurement of the number of tag integrations. In fact, dPCR readouts strongly rely on how clearly fluorescent and non-fluorescent droplets can be distinguished. For example, we observed that the results of one dPCR assay substantially differed using annealing temperatures of 61.2°C and 64.4°C (Supplementary Fig. 2C), meaning that a couple of degrees of difference may significantly affect the overall assay performance.

CRISPR efficiency depends, amongst other factors, on Cas9/gRNAs cutting efficiency and specificity, recipient cell type and CRISPR reagents transfection method. Importantly, the method presented here may be adopted to accurately and efficiently assess newly published reagents for their editing efficiency. To this purpose, we show in the current protocol how dPCR successfully enabled us to compare two different sets of CRISPR reagents (PPP and PRNP) by measuring their editing efficiencies. Our strategy can easily be extended to any new experimental design and/or reagent whose efficiency – either in terms of HDR, NHEJ events or both – needs to be quantitatively assessed.

A preliminary test of the designed dPCR specificity can be carried out by performing a quantitative PCR (qPCR), using the pre-optimized annealing temperature and identical components of the dPCR experiment. Such a test allows to estimate the threshold cycle (Ct) of the designed dPCR assay. We assumed the assay to be specific when Ct ≤ 30 using ∼25 ng of genomic template DNA. A non-template control (NTC) and a wild-type (i.e. non-edited) genome were included as negative controls (Supplementary Fig. 2B).

## LIMITATIONS

Advances in the design and delivery of CRISPR reagents together with improved strategies to sort and screen edited clones led us to obtain efficiencies of homozygous clones in polyploid cells up to ∼20% (Fig. 5). However, even our improved pipeline can still require to troubleshoot some technical limitations linked to each step of the newly outlined CRISPR pipeline, which we summarize here.

First, electroporation represents a more labour intensive technique when compared to lipofection and electroporation conditions should be carefully adjusted for each recipient cell line and mix of CRISPR reagent used, implying a preliminary optimization before implementation to create novel cell lines. Nevertheless, because it maximizes delivery rates up to 35% with three expression plasmids simultaneously and survival up to ∼95% in our hands, we would now recommend electroporation as the method of choice for CRISPR genome editing for homozygous knock-ins. When aiming for high-throughput approaches, suitable electroporation platforms and experimental designs are commercially available and can also be adapted to the cell type and CRISPR reagent combination used.

Concerning FACS selection of genome-edited cells, it is good to be aware that the use of multi-functional self-labeling tags such as SNAP or Halo requires the subsequent coupling of organic fluorophore ligands that are compatible with the instrument excitation and emission filter settings of the cell sorter. Also, live cell permeable dyes are generally desired, unless the POI is expressed at the cell surface. In our experience, a major limitation in the use of organic fluorophores with such self-labelling tags is their background signal in cells not expressing the tag. We have established a staining procedure for both SNAP and Halo tags that enabled us to discriminate edited cells from non-specifically stained cells. Nevertheless, further optimization may be needed when attempting to extend the same labelling strategy for tagged proteins with low abundance. Very low expression levels of the POI may result in a fluorescence signal that is undistinguishable from background labelling by FACS, so that cells cannot be reliably sorted. Another fundamental aspect of FACS followed by single cell deposition relies on the possibility to grow genetically pure clonal lines from an individual parental cell. The user should be aware that not all cell types may survive single cell deposition or expand efficiently from single cells. To face this potential limitation, we encourage the reader to carry out a pilot optimization of cell growth conditions after single cell deposition to identify the best combination of media supplements and growth conditions for a particular cell line. For example, we have successfully expanded U2 O-S cells by using pre-conditioned medium (see procedures, 5| FACS).

Automatic (fluorescence) imaging of several clones at a time represents another important improvement of our new CRISPR pipeline. However, we should emphasize that standard tissue-culture plates may severely impair imaging performances because of the plastic thickness and/or transparency. We overcame this limitation by performing bright-field (BF) imaging of the parental plate using a low-magnitude, dry objective (4x) with 20 mm working distance. However, since wide-field fluorescence imaging of expressing cells was incompatible with standard plastic dishes, we performed high-throughput fluorescence detection with cells expanded on thin glass-bottom 96-well plates. In this way, we could obtain two image-based datasets for interactive computer-aided selection of potential good clones at an early step of the CRISPR pipeline.

We emphasized the importance of a careful design of dPCR assays to obtain a reliable assessment of the number of on- and off-target tag integrations. In parallel, we outline here critical aspects and limitations that may affect the accuracy of dPCR results and the following screening outcomes. First, the separation between fluorescent and non-fluorescent droplets is critical to perform an accurate quantification of the DNA of interest. The possibility to distinguish between fluorescent and non-fluorescent droplets is dependent on several parameters, including biochemical and physical factors ^35^. Amongst those, we already mentioned that amplicon lengths >200 bp may impair dPCR amplification efficiency, limiting the possibility to accurately assess the copy number of the target or reference sequence ^36^. Since one of our aims was to quantify the number of on-target mEGFP integrations in HeLa Kyoto host cells, we had to perform dPCR with a large, suboptimal amplicon length of ∼1300 bp. To overcome this challenge, we combined the use of a double-quenched probe with an optimized, three-step dPCR cycling program, as previously described ^29^.

The number of detectable targets in dPCR is limited by the number of fluorescent detectors available in the reader device and a reference assay also needs to be included. At the time when we performed the tests, three fluorescent dyes could be detected in parallel (Naica™ system, STILLA), which means that two different target assays and a single reference assay could be combined in one dPCR reaction. In the current protocol, limitations adjusting PCR conditions to the very different amplicon sizes were overcome by performing a single triple-colour assay that exploited a re-designed CRISPR donor (Fig. 1). Alternatively, two separate dual-colour assays may be carried out (Fig. 1).

One potential drawback of dPCR data-analysis is represented by fluorescence spillover ^37^. This phenomenon must be thoroughly addressed to accurately identify dPCR droplets sub-populations. In practice, we corrected for fluorescence spillover by evaluating the contribution of each fluorophore to the resulting fluorescent signal in each detection channel, as previously described ^37^. Briefly, we prepared single-colour control samples where only one detector can read positive droplets. From these single-colour controls, the amount of interfering fluorescent signal in the other detectors was computed to generate the so-called «compensation matrix». Once calculated, this matrix is used to correct the measured fluorescence values, yielding the actual fluorophore signal. As a result, various droplet populations may be unequivocally identified. We carried out compensation corrections by using the algorithm implemented in the Crystal Miner™ software (STILLA).

Overall, the dPCR assay described in this protocol allows performing a copy-number assessment of mEGFP or SNAP tags as full-length insertions at the locus of interest (i.e. on-target integration) or at any other random insertion locus in the genome (i.e. off-target integration). Therefore, this assay does not account for any micro-insertions or deletions and/or single-point mutations that might have occurred as side-effects of CRISPR genome editing. Detection of such genomic modifications requires dedicated assays such as for example partial or full genome sequencing designed according to the specific mutations to be detected. As a final step in our validation routine, we amplify and perform Sanger sequencing of the genomic region spanning the tagged target in the homozygous knock-in clones obtained.

## PROCEDURES

### Cell culture

HeLa Kyoto cells were grown in high-glucose DMEM medium supplemented with 10% vol/vol heat-inactivated FBS, 100 U/ml penicillin-streptomycin solution, 1.0 mM sodium-pyruvate and 2.0 mM L-glutamine. U-2 OS cells were grown in McCoy5A modified medium supplemented with 10% vol/vol heat-inactivated FBS.

### Design of guide RNAs (gRNAs) ● TIMING 1d

**1|** *sgRNA design*. Two different formats of guide RNAs (gRNAs) were used in this report: the synthetic, chemically modified gRNAs (sgRNAs) to be complexed with the wild-type recombinant Cas9 (wt-Cas9) to form ribonucleoparticles (RNPs), and the paired gRNAs cloned in pX335 plasmid for transient expression with the mutant S.p.Cas9-D10A (Cas9-D10A). Both formats were designed using the CRISPR tool implemented in Benchling (https://www.benchling.com/crispr/), or by IDT (https://eu.idtdna.com/site/order/designtool/index/CRISPR_CUSTOM) as follows:

> (A) Around 100 bp of genomic region spanning the stop codon from the C-terminal region of the target gene were selected with Bluegecko (http://bluegecko/cgi-bin/HeLa_genome.pl) or alternatively with ENSEMBL (https://www.ensembl.org/index.html)

▴ NOTEIf the genome of interest is not available, ENSEMBL can be used as alternative. Otherwise a locus PCR can be performed and sequenced by Sanger Sequencing to know the editing site.

> (A) (B) The retrieved sequence was saved as a GenBank or FASTA file using the SnapGene® editor v5.0.1(www.snapgene.com) and uploaded in the Benchling or IDT CRISPR tool.

gRNAs were designed as either «single guide» or «paired guide», depending on whether wild-type or Cas9D10A (i.e. nickase) was used. As for conventional gene targeting, the location of genome editing must be carefully selected. First, sgRNAs should support the generation of nicks as close as possible to the insertion site ^20^. Moreover, whenever possible, it is strongly advisable to pick gRNAs producing nicks within the stop codon. In fact, such gRNA will produce a cut exclusively on the target genome sequence without affecting the donor DNA. Moreover, the selection of sgRNAs should minimize the likelihood of generating off-target insertions. Here, paired gRNA forward (FW) and reverse (RV) oligos were designed to generate BbsI overhangs after annealing, to allow cloning in the pX335 plasmid using the Golden Gate strategy (see protocol details). Paired gRNAs and chemically modified sgRNAs nucleotide sequences used in this work are listed in Table 1. The sgRNAs have been purchased either at Thermo Fisher Scientific or IDT.

**Table 1:** PCR setup

| Component | stock concentration | Volume [μL] | Final concentration |
| --- | --- | --- | --- |
| HotStar HiFidelity PCR Buffer | 5x | 10 | 1x |
| Q-Solution | 5x | 10 | 1x |
| Primer fwd (See Supplementary Table 2) | 10 μM | 5 | 1 μM |
| Primer rev (See Supplementary Table 2) | 10 μM | 5 | 1 μM |
| HotStar HiFidelity DNA Polymerase | 2.5 units/μL | 1 | 2.5 units/reaction |
| human genomic DNA |  | variable | Optimal~1 ng – 200 ng/reaction |
| nuclease-free H<sub>2</sub>O | | up to 50μL | |

### Cloning of paired gRNAs in pX335 ● TIMING 7d

**2|** *gRNA cloning*. Cloning of paired gRNAs into pX335 plasmid implies the design of complementary forward and reverse oligos for each gRNA flanked by short sequences to generate a BbsI overhangs on each side (Table 1). This was achieved as follows:

A. Purify the receiving plasmid pX335-U6-Chimeric_BB-CBh-hSpCas9n(D10A) using the EndoFree®Plasmid Maxi kit.
B. Resuspend forward and reverse gRNA oligos in dH2O at 100 µM final concentration.
C. Mix 1 μL of each oligo with 1 μL T4 ligation buffer 10x, 0.5 μL T4 PNK and 6.5 μL dH2O to reach a 10 μL final reaction volume.
D. Incubate the reaction at 37°C for 30 minutes, 95 °C for 5 minutes and ramp down to 25 °C at 5 °C/minute to anneal oligos.
E. Dilute phosphorylated, annealed oligos to 100µL with dH2O.
F. Assemble a Golden Gate digestion/ligation by mixing 1µL of diluted oligos with 25 ng pX335-U6- Chimeric_BB-CBh-hSpCas9n(D10A), 12.5µL Rapid Ligation Buffer 2x, 1µL BbsI restriction enzyme, 2.5µL BSA 10x, 0.125µL T7 Ligase and 7µL H2O. In parallel, prepare a negative control by replacing oligos by dH2O.
G. Incubate reactions in a thermocycler for 6 cycles of 5 minutes at 37°C and 5 minutes at 20°C.
H. Transform 100µL of chemically competent bacteria with 2µL of each Golden Gate reaction and negative control. Incubate cells on ice for 5 minutes and heat-shock at 42°C for 45 seconds.
I. Transfer heat-shocked cells into ice for 1 minute and supplement with 300µL of Terrific Broth at room temperature.
J. Incubate at 37°C for 30 minutes with shaking (∼800 rpm).
K. Plate ∼50-100µL of transformed cells onto ampicillin-agar Petri dishes under sterile conditions and incubate overnight (ON) at 37°C to allow colonies to grow.
L. The next day, pick 3 individual colonies and inoculate them separately in ∼3-5 mL of LB medium + ampicillin to amplify DNA.
M. Purify plasmid DNA using the QIagen MiniPrep Kit and perform Sanger sequencing to check for the correct insertion of gRNAs at the expected BbsI restriction site in the pX335 plasmid.
N. Identify the correct pX335 construct and grow in ∼200 mL of LB medium + ampicillin in order to prepare a larger amount of DNA.
O. Purify the plasmid DNA using the endotoxin-free Maxi Prep kit from QIagen and determine the concentration using the Nanodrop device (Thermo Fisher Scientific).

### Donor design ● TIMING 1d

**3|** *Donor design.* Two different donor plasmids were used to target the C-terminal regions of Nup93 and TPR genes with the monomeric, enhanced GFP (mEGFP) and the SNAP-tag, respectively, in either HeLa Kyoto or U-2 OS human cell lines.

▴ NOTE Forthe PCRNP approach a 5’-phosphorylated PCR donor has been used.

A. Retrieve from ENSEMBL the C-terminal genomic sequences of each target locus (entries ENSG00000102900 for Nup93 and ENSG00000047410 for TPR, in this protocol) and generate a new sequence file using e.g. SnapGene® editor.
B. Annotate the position of the stop codon within the genomic sequence; the linker and the tag nucleotide sequences were inserted in frame at the 3’-end, making sure to remove the stop codon from the genomic sequence and to avoid adding any start codon via the linker or the tag sequence.
C. Add left and right homology regions based on the retrieved genomic sequence by extending ∼800 bp upstream and downstream of the stop codon. To check whether the donor sequence is in frame, the amino acid sequence of the last exon obtained from ENSEMBL (transcripts IDs: ENST00000308159.10 for Nup93 and ENST00000367478.9 for TPR) should match exactly the translated sequence of the donor. In parallel, the translation frame of the linker and the tag can be checked.

▲NOTE for the PCRNP approach the homology arms have been reduced to 40bp each.

(D) Both donor vectors presented in this work – pMA-mEGFP and pMA-SNAP – have been obtained via gene synthesis (Thermo Fisher Scientific) in the pMA backbone (available from the GeneArt Cloning Vector collection) and sequence-checked (see Supplementary Material for the full sequences).

▲NOTE For PCRNP approach, a synthetic gBlock has been purchased at IDT and reconstituted in TE buffer pH8.0 to a concentration of 10ng/µL. The generation of 5’- phosphorylated PCR product from reconstituted gBlock template (15ng) was obtained with 5’-phosphorylated primers. Previous comparisons of mEGFP expression 48 hours post electroporation, between unmodified and 5’-phosphorylated donors as HDR templates have shown that the ratio of mEGFP expressing cells was significantly higher in the 5’-phosphorylated donor cell population (Supplementary Fig. 5).

### Electroporation ● TIMING 1d

**4|** *Electroporation*. Electroporation was used to deliver CRISPR reagents either as plasmids, PCR-products or RNPs. In this work, we used the Neon® Transfection platform (ThermoFisher Scientific) to perform bulk electroporation of HeLa Kyoto or U-2 OS cells using the 10µL-tip format. HeLa Kyoto cells were electroporated using either RNPs and pMA-mEGFP donor plasmid (PRNP method) or with three plasmids, (named triple-plasmid or PPP method) two of them expressing the gRNAs and the Cas9-D10A with the BFP reporter gene and the third one – that is, pMA-mEGFP – used as donor vector. Importantly, both CRISPR formats targeted the same C-terminal genomic region of the Nup93 locus. U-2 OS were electroporated using the PPP approach. Specifically, the pMA-SNAP donor vector was co-transfected together with the two plasmids expressing the gRNAs and the Cas9-D10A fused to either a mEGFP or a BFP reporter gene to target the C-terminal region of the human TPR locus. In a third CRISPR experiment (PCR-product in combination with RNPs or PCRNP method), U-2 OS cells were electroporated with RNPs from IDT (sgRNA and HIFI Cas9 V3) combined with a phosphorylated PCR product as HDR donor.

A. Electroporation of HeLa Kyoto and U-2 OS cells with three plasmids (triple-plasmid approach, PPP).

i. Harvest HeLa Kyoto or U-2 OS cells from a 10 cm dish (∼70-80% confluent) and resuspend in 10 mL growing medium.
ii. To count cells, mix 18µL of cell suspension with 2µL of acridine orange/propidium iodide solution and add 12µL into a fluorescent photon slide. Count twice using Luna™ Dual Fluorescent Cell Counter (Logos).
iii. Wash cells twice with 10 mL DPBS (without Ca^2+^ and Mg^2+^) and pellet at 200×g, 5 minutes at room temperature (RT). Discard the supernatant and resuspend the cell pellet in ∼2 mL of DPBS.
iv. Pellet 1.0 x 10^5^ HeLa or 1.5 x 10^5^ U-2 OS cells/electroporation reaction in 1.5 mL- Eppendorf tubes using slow speed (200×g, 10 minutes, RT).
v. Discard DPBS supernatant and resuspend the cell pellet in 100µL of Buffer R supplemented with 1.75 µg of each gRNA-encoding plasmid and 2.5 µg of pMA-mEGFP or pMA-SNAP donor vector, for Hela Kyoto and U-2 OS cells, respectively.
vi. Electroporate cells using the 100µL-disposable tip using the following settings: Hela Kyoto: 1005 Volts, 35 msec, 2 pulses; U-2 OS: 1230 Volts, 10 msec, 4 pulses.
vii. Immediately after the pulse, transfer cells into 500µL of 37°C pre-warmed RPMI1640 supplemented with 10% vol/vol FBS; incubate for ∼15 minutes at 37°C/ 5%CO2 to allow cells to recover from electroporation.
viii. Transfer the entire volume of electroporated cells in 2.0 mL of pre-warmed and CO2-equilibrated, antibiotic-free growing medium supplemented with 40 µM final concentration of HDR enhancer from IDT in one well of a 6-well plate and put back to the incubator.
ix. Change medium after 24hours and incubate another 24 hours before proceeding to FACS.
B. Electroporation of HeLa Kyoto cells with RNPs and pMA-mEGFP donor plasmid (Plasmid-RNP hybrid approach, PRNP).

i. Prepare HeLaKyoto cells as described in Section (A).
ii. Meanwhile, assemble RNPs by mixing 12.3 pmol of recombinant S.p. HiFi Cas9 V3 with 86 pmol of sgRNA in 7.4µL of Buffer R and incubate for ∼20 minutes at RT. Afterwards add the Electroporation Enhancer reagent to achieve a final concentration of 4 µM and 2µg of the plasmid donor (4685bp) in a total volume of 10µL.
iii. Resuspend 1×10^5^ cells with the RNPs reaction.
iv. Electroporate HeLa Kyoto cells at 930 Volts, 30 msec, 2 pulses using the 10µL-disposable electroporation tip.
v. Immediately after the pulse, transfer cells into 500µL of 37°C pre-warmed RPMI1640 supplemented with 10% vol/vol FBS; incubate for ∼15 minutes at 37°C/ 5%CO2 to allow cells to recover from electroporation. Transfer the entire volume of electroporated cells in 2 mL of of pre-warmed and CO2-equilibrated, antibiotic-free growing medium supplemented with 40 µM final concentration of HDR enhancer from IDT in one well of a 6-well plate and put back to the incubator.
vi. Change medium after 24 hours and incubate another 24 hours before proceeding to FACS.
C. Electroporation of U-2 OS cells with RNPs and a 5’-phosphorylated PCR product (PCRNP- approach).

i. Prepare U-2 OS cells as described in Section (A).
ii. Meanwhile, assemble RNPs by mixing 100 pmol of recombinant S.p. HiFi Cas9 V3 with 100 pmol of sgRNA in Buffer R and incubate for ∼20 minutes at RT. Afterwards add the Electroporation Enhancer reagent to achieve a final concentration of 4 µM and the PCR-donor in a total volume of 10µL.
iii. Resuspend 1.5×10^5^ cells with the RNPs reaction.
iv. Electroporate U-2 OS cells at 1400 Volts, 15 msec, 2 pulses using the 10µL-disposable electroporation tip.
v. Immediately after the pulse, transfer cells into 500µL of 37°C pre-warmed RPMI1640 supplemented with 10% vol/vol FBS; incubate for ∼15 minutes at 37°C/ 5%CO2 to allow cells to recover from electroporation. Transfer the entire volume of electroporated cells in 2 mL of pre-warmed and CO2-equilibrated, antibiotic-free growing medium supplemented with 40µM final concentration of HDR enhancer from IDT in one well of a 6-well plate and put back to the incubator.
vi. Change medium after 24hours and incubate another 24 hours before proceeding to FACS.

### FACS ● TIMING 1d

#### 5|

A. *Pre-staining of cells.* Pre-staining with a SNAP-reactive fluorophores is necessary only when cells have been transfected with a donor encoding SNAP-tag.

i. To stain SNAP-expressing U-2 OS cells, dilute SNAP-Cell® 647-SiR dye to a final concentration of 1 µM growing medium and put on cells.
ii. Incubate for 30 minutes at 37°C/ 5%CO2.
iii. Wash labelled cells three times with pre-warmed DPBS and incubate growing medium for an additional 30 minutes at 37°C/ 5%CO2 to wash unbound dye.
iv. Repeat step C three additional times before harvesting cells by trypsinization.
B. *Preparation of U-2 OS and HeLa cells for FACS sorting.* Electroporated HeLa Kyoto or U-2 OS cells are harvested for pool-sorting 48h post-electroporation.

i. After trypsin inactivation with growing medium, centrifuge cells (200x g, 10 minutes, RT) and resuspend pellets in 1.0 mL FACS buffer (DPBS + 2% vol/vol FBS)
ii. Filter cells through a cell-strainer to remove cell clumps; keep cells on ice until sorted.
iii. Perform pool-sorting of cells with a BD FACS Aria™ Fusion cell sorter (BD Biosciences) using a 130 μm nozzle. To exclude dead cells, propidium iodide (PI) or DRQ7 viability dies were added 1:1000 vol/vol immediately prior to cell sorting. Doublets were carefully excluded by plotting FSC-height versus FCS-area and SSC-height versus SSC-area; cells with an increased area were discarded. *Preparation of U-2 OS and HeLa cells for FACS sorting.* Electroporated HeLa Kyoto or U-2 OS
(C) *Single-cell sorting*. Single-cell sorting was performed 5 days post-electroporation. HeLa Kyoto and U-2 OS cells were treated as described in Step 5| (B) for pool-sorting. Individual cells were collected into 96-well plates containing growing medium supplemented with 1% v/v Pen/Strep solution and grown for additional ∼10-15 days. Importantly, U-2 OS cells were plated in conditioned cell growth medium supplemented with 1% v/v Pen/Strep.

▴ **NOTE** For the PCRNP method : Single-cell sorting has been performed 48 hours post electroporation without the intermediate pool enrichment step.

### Automated BF/fluorescence imaging ● TIMING 1d

**6|** *Automated BF/fluorescence imaging.* Automated bright field (BF) and wide-field (WF) fluorescence imaging of mEGFP- or SNAP-expressing cells seeded in 96-well plates was performed using an ImageXpress Micro instrument (Molecular Device). BF imaging was performed ∼10 days after single-cell FACS sorting and represented an essential step to identify wells containing colonies developing out of single cells. WF fluorescence imaging was performed after one replica-plating of the parental 96-well plate and subsequent expansion and allowed us to check for correct subcellular localization of the tagged-protein in a fully automated fashion. Note that individual colony detection was performed using the plastic 96- well plates where cells were primarily sorted, whereas WF fluorescence imaging was performed on replicates seeded on glass-bottom plates compatible with fluorescence imaging.

A. BF imaging of plastic plates for single colony detection

i. Transfer the 96-well plate containing growing colonies directly into the ImageXpress Molecular device, pre-equilibrated at 37 °C.
ii. BF imaging using the settings listed in Supplementary Table 1
iii. Perform a visual inspection of generated images by running the Plate viewer plugin available through Fiji (https://doi.org/10.5281/zenodo.3522688). Select wells presenting single colonies for further propagation.
B. WF fluorescence imaging of glass-bottom plates for subcellular protein localization

i. Make a replicate of the wells selected above on a glass-bottom 96-well plate.
ii. Allow cells to adhere overnight at 37°C/ 5% CO2. The next day, replace the cell culture medium with pre-warmed 1X PBS and directly image using the ImageXpress Molecular device, pre-equilibrated at 37 °C.
iii. Perform combined BF- and WF-imaging using the settings described in Supplementary Table 1. Set imaging conditions according to either mEGFP (for HeLa Kyoto expressing Nup93-mEGFP) or TMR (for U-2 OS cells expressing TPR-SNAP labelled with TMR-SNAP dye).
C. Visual inspection of images using Fiji. We generated the images displayed in the Anticipated Results section using the following plugins:

i. run Brightness/Contrast
ii. run Enhance Contrast with a saturated value=0.35
iii. set LUT to green
iv. transform image type to RGB colour
v. save images as TIFF

### PCR ● TIMING 1d

#### 7| PCR to determine hetero- and homozygosity

A. Isolation of genomic DNA (gDNA).

i. To obtain a sufficient amount of gDNAs, the equivalent of two to three confluent wells of a 96-well plates per engineered clone should be pooled.
ii. Isolate gDNAs using the Wizard® SV 96 Genomic DNA Purification System (Promega) and the Vac-Man® 96 Vacuum Manifold (Promega) according to manufacturer’s instructions.
iii. Elute gDNAs with 100µL of nuclease-free dH2O containing 1:125 v/v of RNase (provided in the kit).

▲ **NOTE** To overcome any risks of cross-contamination or pipetting mistakes, the PLATEMASTER 220µL (Gibco) was used in all pipetting steps.

(B) Reaction assembly and thermocycling.
  i. Thaw 5x HotStar HiFidelity PCR Buffer, primer solutions and Q-Solution.
  ii. Mix the solutions completely before use.
  iii. Prepare the reactions according to Table 1.

▲ **NOTE** It is not necessary to keep PCR tubes on ice since HotStar HiFidelity DNA Polymerase is inactive at room temperature)

(iv) Mix the reaction solution thoroughly and dispense appropriate volumes into PCR tubes.
(v) Add template DNA into individual tubes containing the reaction mix.
(vi) Mix and spin down the reaction.
(vii) Perform the thermal cycling program depending on the cell line and primers (for HK- Nup93-mEGFP Table 2A and for U2OS-TPR-SNAP Table 2B).

(C) Gel electrophoresis

i. Prepare a 0.8% (w/v) agarose gel using 1× TBE and 1:10,000 SYBR-safe.
ii. Submerge the gel in electrophoresis tanks containing 1× TBE.
iii. Load 10μL of Gene Ruler as a marker into the gel.
iv. Add 10μL of 6× loading dye to the PCR product and load into the gel
v. Run the samples at 50 V overnight at 4 °C

**Table 2A:** PCR cycling conditions for HK-Nup93-mEGFP

| cycle | step | Time | Temperature [°C] |
| --- | --- | --- | --- |
| 1 | initial activation step | 5 min | 95 |
| 35 | denaturation | 15 sec | 94 |
|  | annealing | 1 min | 61 |
|  | extension | 3 min | 72 |
| 1 | final extension | 10 min | 72 |
| 1 | end | indefinite | 4 |

**Table 2B:** PCR cycling conditions for U2OS-TPR-SNAP

| cycle | step | Time | Temperature [°C] |
| --- | --- | --- | --- |
| 1 | initial activation step | 6:30 min | 95 |
| 20 | denaturation | 30 sec | 95 |
|  | annealing | 45 sec | 65-55 (-0.5/cycle) |
|  | extension | 2:30 min | 68 |
| 20 | denaturation | 30 sec | 95 |
|  | annealing | 45 sec | 55 |
|  | extension | 2:30 min | 68 |
| 1 | final extension | 6:30 min | 68 |
| 1 | end | indefinite | 4 |

#### Digital PCR ● TIMING 1d

**8|** *Digital PCR (dPCR) data acquisition and analysis*. (A) We first used the NAICA™ platform to determine the tag copy-number of edited HeLa Kyoto or U-2 OS clones pre-validated based on their Southern Blot profile. (B) We used the QX200 AutoDG Droplet Digital PCR System (BIORAD) to perform larger scale, dPCR-based screening of samples prepared in 96-well plate format. The screening was performed to compare homozygosity yields of two different CRISPR strategies used to tag the C-terminus of Nup93 with mEGFP in HeLa Kyoto cells. Specifically, the standard, PPP approach was compared with a PRNP approach. Furthermore, we included here an additional assay – namely, the «HDR assay» - to estimate on-target, specific integrations of mEGFP at the target locus. This allowed us to classify the screened samples in homozygous, heterozygous and clones with off-targets generated by each CRISPR strategy. The combined on-target and total copy-number assessment (PCRNP) is additionally described.

A. Isolation of genomic DNA (gDNA) from monoclonal colonies (Supplementary - Section 11|(B) (i))

(i) dPCR with NAICA™ system (STILLA). Assemble a 12.5x assay mix in a final volume of 100µL (Table 8B, 8D, 8F).

▲ **NOTE** Primer and probe sequences and amplicon sizes can be found in Supplementary Table 9.

ii. Assemble the dPCR reactions (Table 8A).
iii. Thoroughly vortex and spin down the dPCR reaction. Transfer into an oil-pre-filled well of a Sapphire Chip (STILLA).

▴ **NOTE** For both target and reference assays, an extra compensation assay was prepared. Each compensation assay combines target and reference probes but one set of primers has to be omitted.

iv. Perform partitioning using the Naica Geode (STILLA) system according to STILLA default program followed by the PCR program (Table 8G).
iv. Run up to three Sapphire Chips simultaneously.
vi. Image the processed Sapphire Chips by transferring them into Prism3 reader (STILLA). Operate with CrystalReader (STILLA) software.

▲ **NOTE** Each assay has its optimal exposure time (Table 8) which needs to be empirically optimized.

vii. Compute compensation after imaging Sapphire Chips using the dedicated tool implemented within CrystalReader software. Automated fluorescence thresholds were double-checked to account for reliable separation between positive – that is, fluorescent – and negative partitions.
viii. Export as *.csv* the file containing the number of total partitions, the number of positive partitions in target- and reference-channels using the CrystalMiner software.

(B) dPCR with QX200 AutoDG Droplet Digital PCR System (BIORAD).

▲ **NOTE** Primer and probe sequences and amplicon sizes can be found in Supplementary Table 9.

i. Prepare the dPCR reactions according to Table 9A or 9C
ii. Generate partitions according to manufacturer’s instructions.
iii. After partitioning, seal ddPCR™ 96-Well Plates (BioRad) using C1000 Touch™ Thermal Cycler (BioRad).
iv. Run the PCR program using a 96-Deep Well Reaction Module (BioRad). We ran both programs to measure the total number of mEGFP integrations in the whole genome (allGFP assay) or the total number of specific on-target integrations achieved at the locus of interest (HDR assay) (Table 9B or 9D).
v. After PCR cycling, read each processed 96-well plate according to manufacturer’s instructions using the QuantaSoft™ Analysis Pro (BioRad). The number of total partitions, and number of positive partitions in target- and reference-channel have been exported as a *.csv* file.

(C) (dPCR data analysis.

i. Integrated tag copy-number can be readily computed by using the Excel macro provided by STILLA. To perform such calculation:

a. Get the template from https://www.gene-pi.com/statistical-tools/poisson-law-2/.
b. Complete the file with the following dPCR information (indicated as «INPUT» in the column header):

i. Total number of partitions
ii. Number of positive partitions
iii. Stock dilution (e.g. dilution 10 = diluted 10 times from stock to well; dilution 1= output is the concentration in the well).
iv. Click Submit to automatically download results.
v. Use any suitable software to compute box-plots out of dPCR replicated measurements.
ii. dPCR data obtained from BioRads QX200 AutoDG Droplet Digital PCR System (Section 13|(B)) were analysed to classify CRISPR-generated genotypes as either homozygous, heterozygous or presenting off-target integrations of the tag. Such a classification relied on the combination of two statistical tests readouts.
iii. Compute and normalise the reference distributions of the copy numbers for the «allGFP» and «HDR» dPCR-assays and represent the theoretical distributions (Figure 5A).
iv. To determine the heterozygosity or homozygosity of each clone, we compared the distributions of the copy numbers of «allGFP» and «HDR» with the two-sided *t*-test (Figure 5B). For that purpose, we computed the probability – that is, the *p*-value – of having a clone with a copy-number equal or different than the mean copy-number of the normalized reference distribution by chance (clone #38, representing the homozygosity). FDR-adjusted p-values of «allGFP» and «HDR» dPCR assays were computed after performing the two-sided t-test, using a significance threshold, α = 10 %.
v. The two distributions could be considered similar only when the score is less or equal than 2 (**ζ** ≤ 2), and different when the score is greater than 3 (**ζ** > 3). In the latter case, i.e. for those cases in which the two distributions are dissimilar, and the copy number «allGFP» is greater than the copy number «HDR», the clones are classified as containing off-target. In those cases in which the distributions is similar and the mean of the total mEGFP and the on-target mEGFP coincide, then the clones are homozygous; otherwise, they are classified as heterozygous (Fig. 5C), with one or 2 alleles tagged out of 3. This process was repeated twice for two different CRISPR strategies: PPP and PRNP. The off-target rate was more prominent with the PPP technique (74 %) than with the PRNP (20 %) as expected. PRNP generated the highest percentage of heterozygous clones with 60 %, in contrast to the 13 % for PPP. Regarding the homozygous rate, PRNP showed a rate of 20 % while in PPP 13 % of clones are homozygous (Fig. 5D).
vi. As for what concerns the generation of box-plots and pie charts reported in this work, we opted to integrate the numerical and the statistical analysis in an R notebook. For reproducibility purposes, the notebook can be explored and re-run here: https://mybinder.org/v2/gh/beatrizserrano/test_AM/master?filepath=https%3A%2F%2Fgithub.com%2Fbeatrizserrano%2Ftest_AM%2Fblob%2Fab51b016b991982941eb244b23bcbdd07bee290c%2F1_analysisReplicates.Rmd

### Confocal imaging ● TIMING 2d

**9|** *Confocal imaging*. Fluorescence confocal imaging of representative clonal cells was performed using a DMI8-CS microscope (Leica).

A. Cell seeding

i. Seed ∼5.0×10^4^ cells of each representative clone into 8-well Lab-Tek chambers (#1, Thermo Fisher Scientific)
ii. Incubate overnight at 37°C/ 5% CO2 before proceeding either to staining (Section 12 (B)) or directly to imaging (Section 12 (C)).
(B) Staining of SNAP-expressing U-2 OS cells.

i. Incubate U-2 OS cells with 0.5 µM TMR-Star dye for 30 minutes at 37°C/ 5% CO2.
ii. Wash cells three times with pre-warmed, growing medium; add fresh medium and incubate for additional 30 minutes at 37°C/ 5% CO2.
iii. To minimize the background signal coming from unbound dye, repeat Step (ii) three times before continuing with imaging.
C. Imaging and image rendering.

i. Replace the growing medium ∼1 hour before imaging with pre-warmed imaging medium (for HeLa Kyoto cell lines: phenol red-free DMEM supplemented with 4.5 g/L glucose, 10% v/v heat-inactivated foetal bovine serum (FBS), 1 mM Natrium Pyruvate (NaPyr), 2 mM L-Glutamine (L-Gln) and 100 U/mL Penicillin/Streptomycin (P/S) solution; for U-2 OS cell lines: phenol red-free DMEM supplemented with 3 g/L glucose, 10% v/v heat-inactivated FBS, 1.5 mM L-Gln, 1% v/v NEAAs and 100 U/mL P/S solution).
ii. Select a field of view (FOV) of 184.52×184.52 µm to be imaged using the XYZ unidirectional scanning mode with a HC PL APO CS2 63x/1.20 NA water immersion objective.
iii. Image HeLa Kyoto cells at 0.180 µm XY-resolution, and U-2 OS cells at 0.361 µm XY- resolution.
iv. Select the 488 nm excitation line of an Argon laser to image mEGFP or HeNe 633 nm to image TMR Star fluorophore covalently bound to SNAP-tagged proteins.
v. Set the scanning speed to 100 Hz.
vi. Collect the fluorescence signals by setting the HyD SMD1 detector. For mEGFP, set the detection range between 494 and 546 nm; for TMR-Star, adjust the wavelength range from 571 to 601 nm.
vii. Adjust the master gain to ∼90-100%.
viii. Render images using the Fiji open-source software. Open the raw *.lif* files collected from DMI8-CS microscope and overlay the «*Fire*» lookup table (LUT) to each representative image.
ix. Adjust the brightness and contrast using the automatic enhancement; set the image type to Red-Green-Blue (RGB) format.

▲ **NOTE** (OPTIONAL) Flatten annotations before saving pictures as TIFF images.

### Protein expression validation ● TIMING 4d (10A) / 1d (10B) (see Supplementary Methods)

**10A|** Western Blot.

**10B|** Capillary Electrophoresis.

### Southern Blot ● TIMING 14d (see Supplementary Methods)

**11|** Southern Blot.

## ANTICIPATED RESULTS

After transitioning to electroporation as the delivery strategy, we tested two different reagent combinations to optimize the CRISPR editing efficiency. With the PPP and PRNP methods, we have observed transient leaky expression of the tag from the donor plasmid which led to selection of cells with free fluorescent tag by cell sorting. When using these methods, FACS displayed ∼39% (PPP) and ∼72% (PRNP) of mEGFP expressing cells (Fig. 2A-B, upper panels). Surprisingly, the corresponding controls electroporated with the donor plasmid alone showed comparable (∼28% and ∼89%) fractions of mEGFP- expressing cells («donor-ctrl » panels, Fig. 2A-B). These results indicated that a significant proportion of fluorescent cells collected at 48 hours post-electroporation expressed mEGFP in an unspecific and transient manner as a result of donor plasmid leakage rather than from the engineered locus. The detection of a transiently expressed tag from a donor plasmid and selection of the corresponding clones should be avoided in order to reduce the downstream handling of non-edited clones. This can be achieved by delaying the FACS analysis until free tag expression has diminished. In our case, we waited three additional days before performing a second round of FACS (Fig. 2A-B, lower panels). Cells expressing mEGFP dropped to ∼11% (PPP) and ∼24% (PRNP), compared to 0.11% and 0.77% in control experiments with donor plasmid alone. Taken together, our results show that a minimum of five days is required to reliably detect mEGFP integrations at the target locus in electroporated cells when using plasmid-based donors that have leaky expression. We also observed that HDR efficiency measured at five days post-electroporation was higher (∼24% vs. ∼11%) when using the PRNP hybrid approach.

The transient donor expression issue has prompted us to develop a third method which makes use of a linear PCR-product as a source of donor DNA instead of a complete plasmid (Fig. 2C, PCRNP). In our proof-of-principle experiment, the measured fraction of mEGFP expressing cells at two days post-electroporation (∼4%) corresponds to the true proportion resulting from HDR, since undetectable expression of mEGFP was measured from the donor PCR only controls. Remarkably, this strategy not only reduced the waiting time from five to two days but also simplified handling with a single FACS round. The single FACS round also reduced stress applied to cells during the protocol and the early single sorting ensures higher genetic diversity among single cell clones as it avoids clonal expansion prior to FACS sorting.

When performing the FACS step, we strongly suggest users to display the data in a dot-plot where the mEGFP fluorescence is plotted against an unused channel; in this case the 561-610/20 (or mCherry) as illustrated in Fig. 2C. This allows to clearly distinguish between mEGFP expressing cells and auto- fluorescence signal, known to display striking correlated patterns along diagonals as previously described Sorted single cells were grown in 96-well plates. The growth period needs to be adapted according to the specific cell line (Fig. 2D). In our hands, ∼10 days to grow represented the shortest time necessary to image single cell derived HeLa Kyoto and ∼14 days for U-2 OS colonies in plates. Wells containing single colonies were identified by performing automated bright-field imaging (BF, Fig. 2D, left panel) followed by running a dedicated Fiji plugin for image analysis ^19^. In our conditions, >50% of wells contained a single viable colony and only a residual proportion (<2%) had more than a single growing colony, indicating a high single-cell dispensing efficiency. The remaining wells were empty due to cell death which is linked to the difficulty of a given cell line to grow a colony from a single cell in the well of a plate. Users should assess growth from single cells for their specific cell type prior to start sorting CRISPR electroporated cells. Once colonies have reached a significant size, they were disrupted for replica-plating on a glass bottom plate compatible with automated fluorescence imaging, in a plastic plate for genomic DNA extraction for PCR and dPCR and another plastic plate for clone propagation (Fig. 2D, Cyan arrows). The identification of proper subcellular localization of the tagged fluorescent protein allowed us to identify undesirable clones and discontinue their downstream propagation at an early step of the protocol (Fig. 2D, right panel). An approximate estimation of clones showing improper fluorescent protein localization is ∼1% for the experiments performed in this study.

To test dPCR as a potential screening tool for our improved CRISPR pipeline, we generated and validated new cell lines using conventional biochemical assays, namely PCR (Fig. 3A) and Southern Blot (Fig. 3B). We selected a subset of clones to validate the prediction power of dPCR. We tagged the TPR locus in U-2 OS (Fig. 3B, left panel) and selected homozygous, heterozygous and clones showing extra-integrated tag copies. After designing a dedicated assay to measure the number of integrated SNAP-tag sequences (Fig. 3C, left panel), we tested the prediction power of dPCR. Analogously, the same scheme and dPCR assay has been used to measure the total number of mEGFP copies at the Nup93 locus in HeLa Kyoto (Fig. 3C, right panel).

The full assessment of the editing procedure relies not only on the quantification of the total number of editing events but also of their specificity. To address this, we designed an on-target dPCR assay to score the number of specific integrations which occurred at the Nup93 locus in HeLa Kyoto cells (Fig. 3, right panels). In this second example, the dPCR results were in agreement with the conventional biochemical analysis (Fig. 3A-B). For both loci we additionally performed confocal imaging (Fig. 3E) to confirm the correct subcellular localization of the tagged proteins and either Western Blot or capillary electrophoresis (Fig. 3D) which were used to assess the protein expression levels and molecular weights.

Western blotting (WB) is a common technique used to check whether successful tagging was achieved at the POI (Fig. 3D, right panel) but is limited when diagnosing large molecular weight (MW) proteins. For example, with a theoretical mass of 267 kDa, wild-type TPR cannot easily be distinguished from its SNAP-tagged version by conventional protein gel-electrophoresis (SDS-PAGE). For this reason, we have used capillary electrophoresis for the TPR-SNAP edited clones (Fig. 3D, left panel). This method is performed within 4 hours including ∼1 hour hands-on time. We recommend the user to perform capillary electrophoresis especially when limited amounts of biological sample are available and whenever multiplexing over several POI is required. Notably, as in Western Blot, optimization of detection conditions for each antibody might be needed. Here, we successfully performed capillary electrophoresis with anti-TPR and anti-SNAP antibodies (Fig. 3D, left panel).

Standard WB was carried out for lower MW POI, in this case Nup93 (∼93kDa). All edited clones show one band at the predicted MW of Nup93-mEGFP (∼125 kDa; Fig. 3D, right panel, anti-Nup93). Noticeably, the predicted heterozygous clone 214 does not express the wildtype allele anymore, speculatively due to INDEL-disruption of one allele. Analogously, extra copies were predicted for clone 92 but they are undetectable (Fig. 3D, right panel, anti-mEGFP). These results clearly show that complementary techniques interrogating genome modifications (dPCR) and protein expression (capillary electrophoresis or WB) are needed to validate CRISPR-generated clones. We therefore encourage the user to perform both techniques.

Given the dPCR accuracy we obtained in our validation experiments, we reasoned that a dPCR-based screening tool enabling both total and on-target tag metrics in a single assay would be ideal. This prompted us to develop a novel, triple-colour assay that was tested with U-2 OS clones, where mEGFP was integrated at Nup93 locus (Fig. 4). Homozygous clones were readily identified through their similar estimates of total and on-target mEGFP integrated copies (Fig. 4D; clones 35, 52 and 247, boxplot) and confirmed by Southern Blot (Fig. 4E) and confocal live cell imaging (Fig. 4F).

We used the allGFP and the HDR-assay in combination with their specific reference assays to assess the tagging efficiency as well as the off-target rate of the PPP and PRNP methods with HeLa Kyoto Nup93- mEGFP clones (Fig. 5). The overall performance is captured in Fig. 5D. The pie-charts show that the CRISPR protocol using RNPs combined with HDR-donor plasmid generated only ∼22% of clones bearing extra mEGFP integrations, showing a significant improvement as compared to the PPP method with ∼73% off-target insertions. It is worth mentioning here that the actual number of clones populating the illustrated pie-charts is lower than the number of clones initially scored. This is the result of a strict quality-control step the user should perform before challenging selected clones through hypothesis testing. First, clones must develop from individual colonies only, as identified from automated BF imaging (Fig. 1). This ensures the highest genetic homogeneity of the CRISPR-engineered cells of a line. Second, selected clones should display a clear PCR pattern to predict the undergone genetic modifications. This implies to handle clones that provided enough of genomic DNA to perform PCR and classify them as potential homozygous or heterozygous clones (Supplementary Fig. 3). Third, we retained clones that displayed the expected subcellular localization of the tagged protein, as determined by wide-field fluorescence microscopy (Fig. 2 for representative results).

Taken together, our results show that compared to our previously published protocol ^1^, the generation of homozygous knock-in cell lines is significantly improved by performing electroporation of an optimized combination of CRISPR reagents consisting of linear PCR product donor and reconstituted RNPs. This allows the tagging of all target alleles in a single CRISPR engineering round and reduces off-target modifications. It also allows proceeding with an early and single FACS step as there is no leaky background donor expression. The dPCR step adds a fast and accurate diagnostic tool which reduces the extent of in-depth downstream analysis and removes the time consuming Southern Blot. The capillary electrophoresis is the method of choice to rapidly obtain quantitative information on expression of tagged proteins. The R notebooks used to compute the results allow fast and unbiased diagnosis of clones and are publicly available at: https://github.com/beatrizserrano/CRISPR_updated_protocol. Overall, following the guidelines provided in this protocol, one can expect to obtain more than 10% of homozygously edited clones devoid of extra integrations out of a single 96-well plate taking into account a ∼50% single cell survivor rate. Besides cell line generation, this method can be used to score the performance of alternative CRISPR design strategies or to test the efficiency of new engineering reagents.

## TROUBLESHOOTING TABLE

| Step | Problem | Possible reason | Solution |
| --- | --- | --- | --- |
| 4 | Low transfection efficiency or low cell survival rate | Sub-optimal electrical parameters | Perform an optimization step of the electroporation parameters for the specific cell line in use |
| 4 | Low transfection efficiency or low cell survival rate | Lipopolysaccharide (LPS)-contaminated plasmid preparation or plasmid concentration too low | Purify plasmids from bacterial endotoxins (endo-free plasmid kit) |
| 4 | Low transfection efficiency or low cell survival rate | Stressed, damaged or *Mycoplasma* contaminated cells | Check cells before performing the electroporation for *Mycoplasma* contamination. Use early-passage cells |
| 4 | Low transfection efficiency or low cell survival rate | Cell density too low or too high | Titrate cell density in a preliminary optimization step |
| 4 | Low transfection efficiency or low cell survival rate | Cells with high passage number | Use early-passage, exponentially growing cells |
| 4 | Arching issues | High salt in the DNA preparation | Use desalting mini-columns to prepare DNA for electroporation |
| 4 | Arching issues | Air bubbles in the electrode tip | Pipette the sample in a slow, smooth, and continuous motion to avoid air uptake |
| 4 | Arching issues | Too high cell density | Titrate cell density in a preliminary optimization step |
| 5 | Weak or no signal | Signal not correctly compensated | Run a sample with a single fluorophore to account for appropriate signal compensation |
| 5 | Weak or no signal | Target protein expression is low | Run appropriate controls (i.e. positive control, untreated control etc.) or consider to introduce an amplification step to maximize the signal (e.g. with labelled antibodies) |
| 5 | Weak or no signal | Too dim fluorophore(s) | Pair a bright fluorophore with a low expressed protein and avoid unnecessary light exposure |
| 5 | Weak or no signal | Incorrect laser/filter set | Ensure that the flow cytometer is equipped with the correct lasers and filter sets for the chosen fluorophores |
| 5 | High signal intensity | Too high labelling fluorophore concentration | Titrate down the concentration of labelling fluorophore |
| 5 | High signal intensity | Too high fluorescence background | Increase the number of washes when preparing cells for FACS |
| 5 | Low event rate | Cell clumping | Use cell strainer before running the FACS analysis |
| 8 | Dirty surface chip | Dusty working environment or contaminated working reagents | Pipette reagents in the chip in a sterile hood. Use suitably purified gDNA and filter tips |
| 8 | No or few positive droplets | The selected restriction enzyme may have cut within the selected amplicon | Change restriction enzyme |
| 8 | No or few positive droplets | The target locus resides in a region rich in secondary structures | Select a restriction enzyme that possibly cuts around the region to be amplified |
| 8 | No or few positive droplets | Sub-optimal annealing temperature | Perform an annealing temperature optimization by running a temperature gradient |
| 8 | No or few total droplets | Sub-optimal concentration of genomic DNA, primers or master-mix | Use the manufacturer's suggested concentrations of gDNA, primers and master mix |
| 8 | No or few total droplets | gDNA not pure | Use only purified gDNA (e.g. with silica-based mini-columns) |
| 8 | No or few total droplets | Presence of big air bubbles | Run the «unstacking.js» script or replace chip |
| 8 | No or few total droplets | Presence of big air bubbles | Always use 25uL of input volume for your Crystal Digital PCR experiment. When loading the chips, make sure not to carry any air bubbles in the pipet tip. When you prepare the PCR, wait 2-3 minutes after the final vortex/centrifugation and before filling the chips' inlet ports |
| 8 | Positive droplets in the no-template control well | Template contamination in the dPCR reagents | Wipe-down pipettes and boxes, use filter tips, prepare master-mixes in a template-free environment and wear appropriate personal protective equipment |
| 8 | Inconsistent concentration results | Insufficient mixing | Thoroughly mix master mixes and all reagents before and after combining them |
| 8 | Inconsistent concentration results | Poor cycler performance | Check the entire plate for a discrepancy in concentration that exceeds the 95% confidence bounds for the wells. If a consistent drop is detected in the same wells, consider replacing the cycler's Peltier plate |
| 8 | Droplet coalescences in some chips | Static electricity | Ensure that chips are not exposed to static electricity: avoid contact with e.g. hair, hands, clothes |

## LIST OF MATERIALS

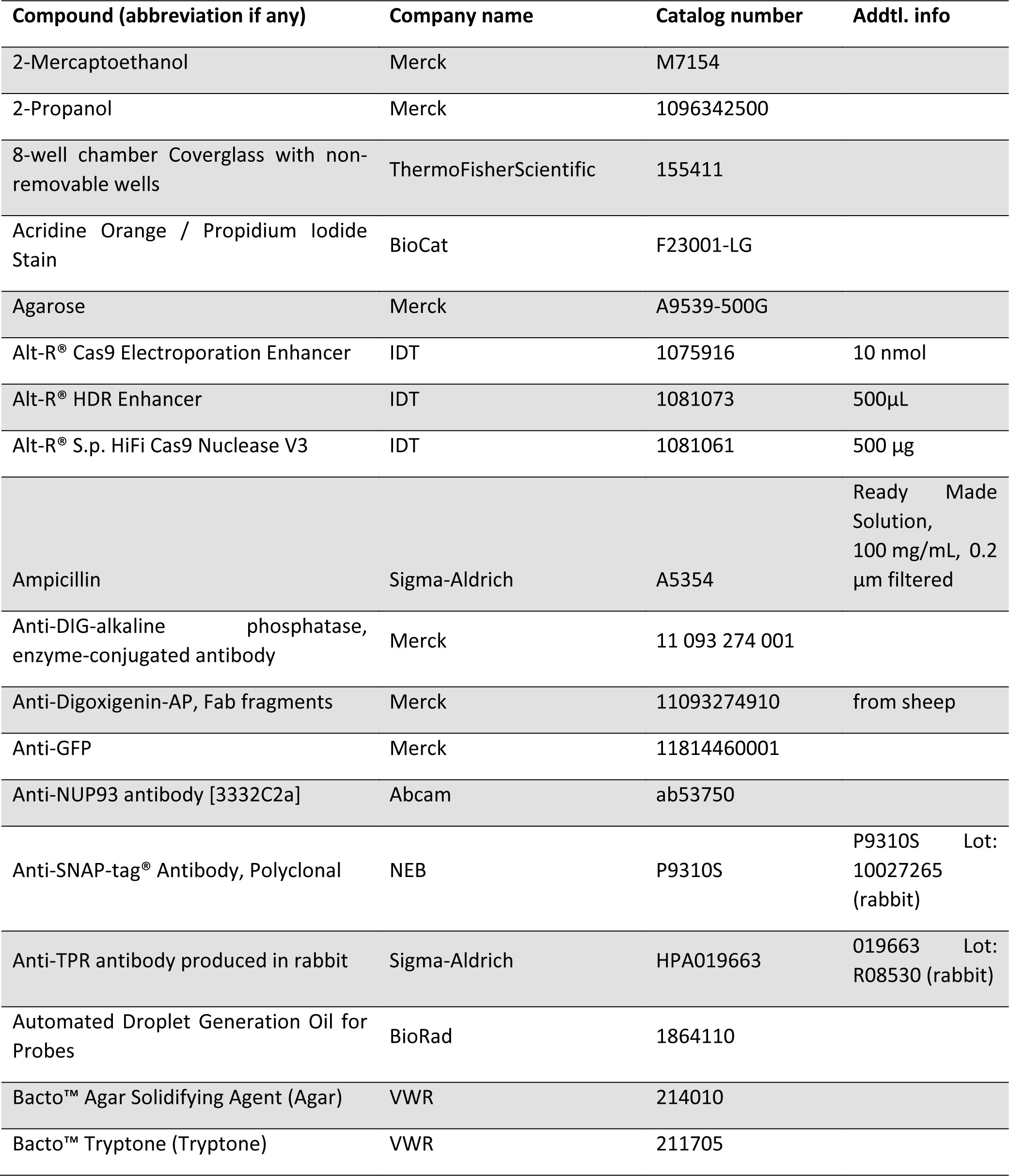
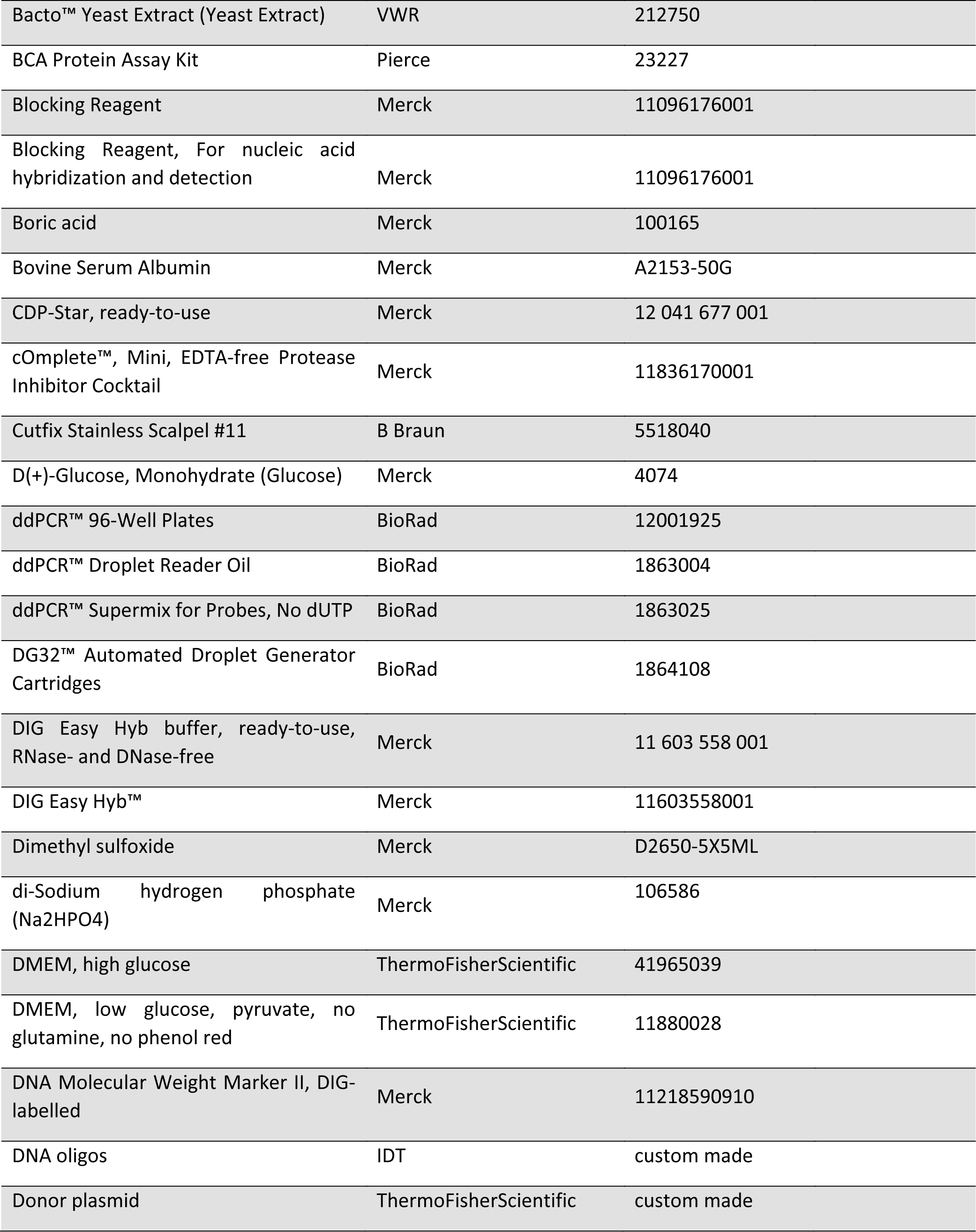
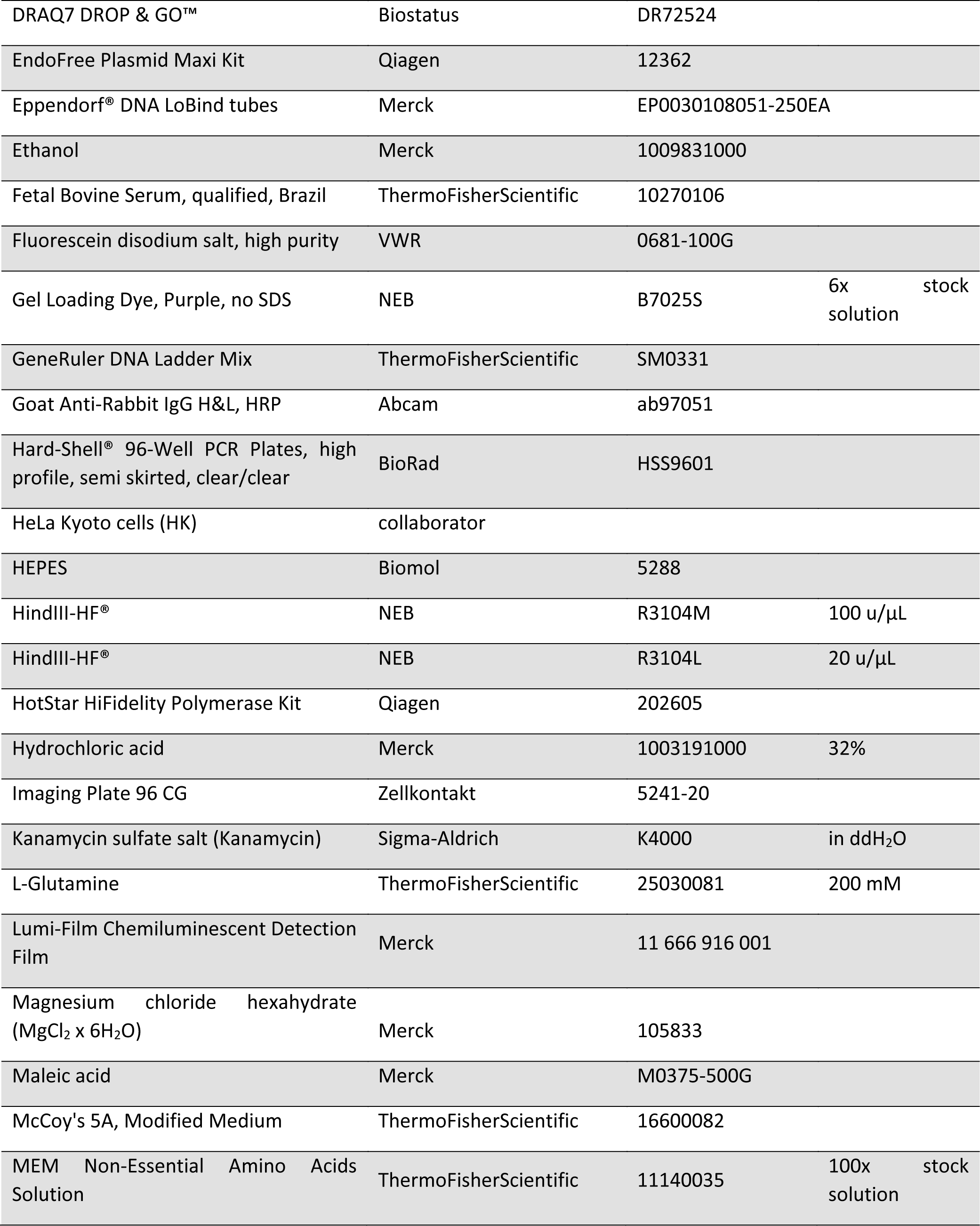
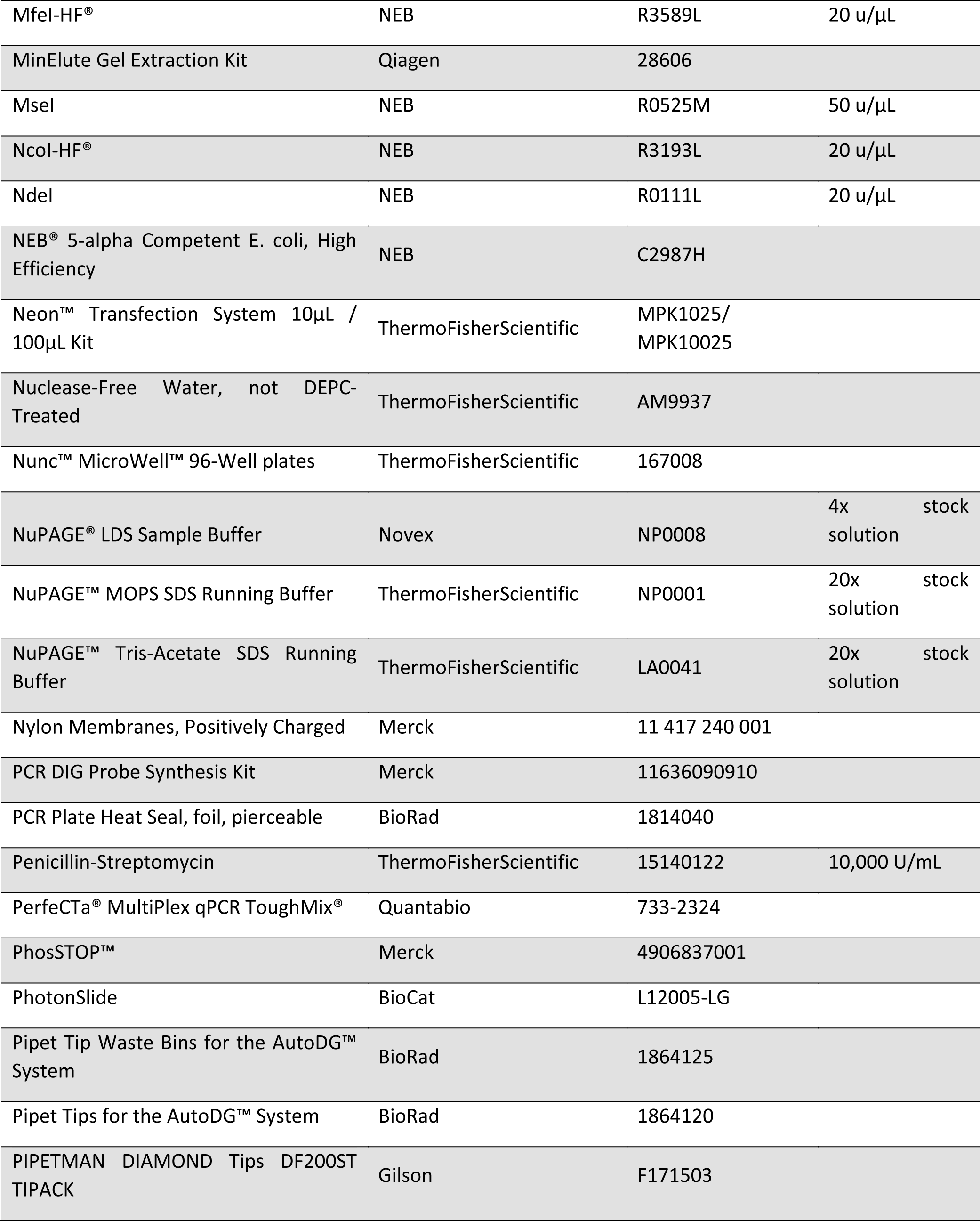
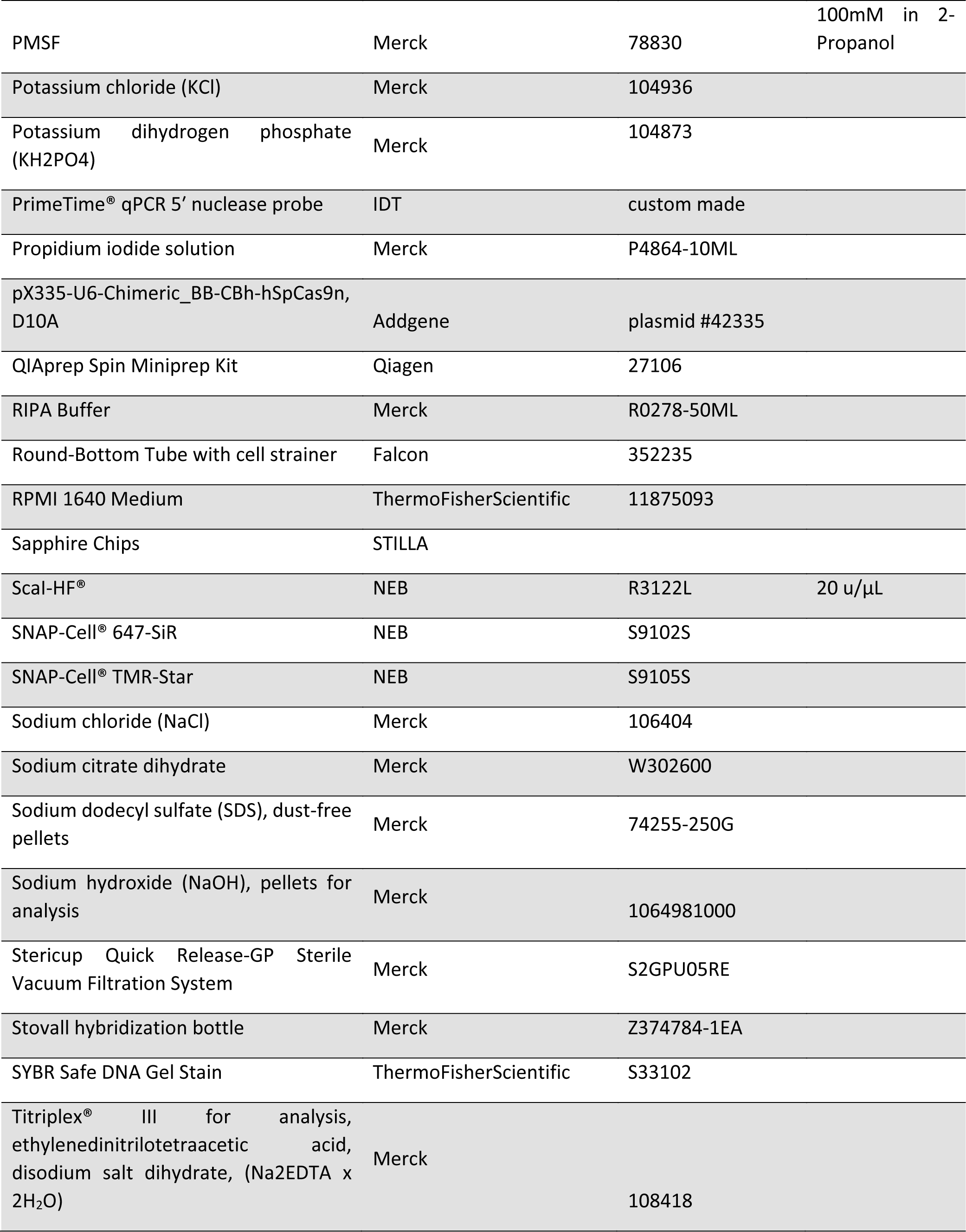
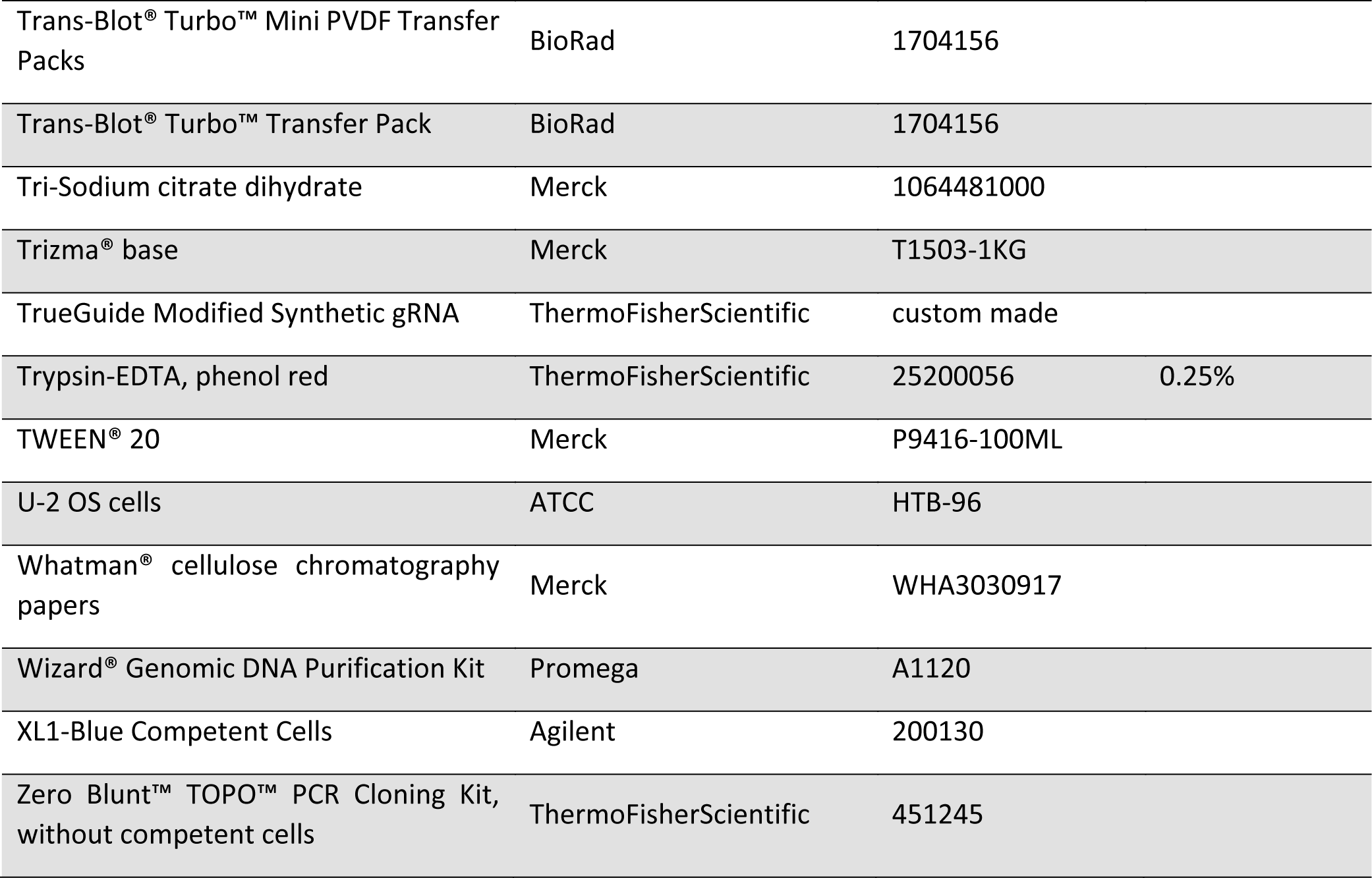

## LIST OF SOLUTIONS

| Solutions (abbreviation if any) | Addtl. info |
| --- | --- |
| 10x TBE | 891.5mM Tris base, 889.5mM Boric acid |
| 20% SDS solution | dissolve in ddH<sub>2</sub>O, filter through a 0.22µm filter |
| 20x SSC | 3 M NaCl, 0.3 M sodium citrate in ddH<sub>2</sub>O, pH 7.0 |
| Anti-Digoxigenin-AP solution | Centrifuge Anti-Digoxigenin-AP for 5 min at ~9000x g in the original vial prior to each use, and pipet the necessary amount carefully from the surface. Dilute Anti-Digoxigenin-AP 1:10,000 (75 mU/ml) in Blocking solution. This solution needs to be prepared fresh. |
| Blocking solution | Dissolve 10% (wt/vol) Roche blocking agent in Maleic Acid Buffer by heating the solution to ~65 °C. Autoclave and aliquot sterile. |
| Denaturation solution | 0.5 M NaOH, 1.5 M NaCl |
| Detection Buffer | 0.1 M Tris-HCl, 0.1 M NaCl, pH 9.5 |
| D-PBS | 2.7 mM KCl, 1.5 mM KH<sub>2</sub>PO<sub>4</sub>, 8.1 mM Na<sub>2</sub>HPO<sub>4</sub>, 136.9 mM NaCl, in ddH<sub>2</sub>O (autoclave) |
| EDTA solution | Na<sup>2</sup>EDTA x 2H<sub>2</sub>O in ddH<sub>2</sub>O, pH 8.0 (autoclave) |
| Glucose solution | 20% in ddH<sub>2</sub>O (autoclave) |
| HEPES, pH 7.9 | 1M in ddH<sub>2</sub>O, adjust pH with NaOH (autoclave) |
| High Stringency Buffer | 0.5x SSC, 0.1% SDS |
| Hybridization solution | DIG Easy Hyb |
| LB-Agar plates | for 1L: 10 g Bactotryptone, 5 g Yeast Extract, 10 g NaCl, 15 g Agar, pH7.0, autoclave (add 100mg/mL Ampicillin or 30mg/mL Kanamycin when cooled down but still liquid) |
| LB-medium | for 1L: 10g Bactotryptone, 5g Yeast Extract, 5g NaCl, autoclave (add 100mg/mL Ampicillin or 30mg/mL Kanamycin when cooled down) |
| Low Stringency Buffer | 2x SSC, 0.1% SDS |
| Maleic Acid Buffer | 0.1 M Maleic acid, 0.15 M NaCl; adjust with NaOH (solid) to pH 7.5 |
| MgCl<sub>2</sub> solution | dissolve in ddH<sub>2</sub>O, 1 M stock solution (autoclave) |
| Neutralization solution | 0.5 M Tris-HCl (pH 7.5), 1.5M NaCl |
| PMSF solution | dissolve in 2-Propanol, stock 100mM, aliquot and freeze at -20°C |
| Stripping Buffer | 0.2 M NaOH containing 0.1% SDS |
| Terrific Broth | 1.2% tryptone, 2.4% yeast extract, 0.5% glycerol + 1x TB salts<br>(10x TB salts = 23.1g KH<sub>2</sub>PO<sub>4</sub>/L, 125.4g K<sub>2</sub>HPO<sub>4</sub>/L) |
| Washing Buffer | 0.1 M Maleic acid, 0.15 M NaCl; pH 7.5; 0.3% (v/v) Tween 20 |

## LIST OF EQUIPMENT

| Instrument | Company | Catalog no. |
| --- | --- | --- |
| BD FACSAria Fusion Cell Sorter | BD | 656700 |
| C1000 Touch™ Thermal Cycler, with 96-Deep Well Reaction Module | BioRad |  |
| Confocal microscope SP8 change in alphabetical order | Leica |  |
| High-Throughput Microscope Molecular Devices IXM | Molecular Devices |  |
| LUNA-FL Dual Fluorescence Cell Counter | BioCat | L20001-LG |
| NAICA™ system | STILLA |  |
| Neon Transfection System | ThermoFisherScientific | MPK5000 |
| PIPETMAN L Multichannel P8x200L, 20-200µL | Gilson | FA10011 |
| PIPETMAN L Multichannel P8x20L, 2-20µL | Gilson | FA10009 |
| PLATEMASTER 220µL | Gilson | F110762 |
| PX1™ PCR Plate Sealer | BioRad | 1814000 |
| Quantum ST5 1100 Series | Vilber Lourmat |  |
| QXDx Automated Droplet Generator | BioRad | 12001630 |
| QXDx Droplet Reader | BioRad | 12001045 |
| Safe floor centrifuge (Avanti J20XP) | Beckman Coulter |  |
| Shaking Hybridization Oven | AMEREX | HS-111 |
| T100™ Thermal Cycler | BioRad | 1861096 |
| Thermomixer Compact | Eppendorf | T1317-1EA |
| Trans-Blot® Turbo™ Transfer System | BioRad | 1704150 |
| Vac-Man® 96 Vacuum Manifold | Promega | A2291 |
| Wizard® SV 96 Genomic DNA Purification System | Promega | A2371 |
| XCell SureLock Mini-Cell Electrophoresis System | ThermoFisherScientific | EI0001 |

## ACKNOWLEDGEMENTS

We want thank Nathalie Daigle for valuable discussions during the entire project and for her great support during the writing of this manuscript. Additionally, we thank our EMBL facilities; the Flow Cytometry Facility, the Advanced Light Microscopy Facility, the Centre for Bioimage Analysis Facility and the Genomics Facility. Furthermore, we highly appreciate the demonstration systems and application specialists from STILLA technologies and Bio-Rad Laboratories.

## SUPPLEMENTARY METHODS

### Protein expression validation ● TIMING 4d (10A) / 1d (10B)

**10A|** Western Blot.

(A) Preparation of HeLa Kyoto nuclear cell lysates.

i. Add 0.5 mL of ice-cold Buffer A (10 mM HEPES, 1.5 mM MgCl2, 10 mM KCl, 0.5 mM DTT, 0.05% v/v NP-40, pH 7.9) supplemented with protease inhibitors into each 10- cm Petri dish containing HeLa Kyoto confluent monolayers.
ii. Scrape off cells from plates and incubate on ice for 10 minutes before centrifuging (780 × g, 10 minutes, 4°C) to get rid of the majority of plasma membranes, DNA and nucleoli.
iii. Resuspend pellets in 376µL of RIPA buffer supplemented with protease inhibitors and 24µL of 5M NaCl.
iv. Transfer lysates into pre-chilled 1 mL Dounce homogenizers and perform ∼20-30 strokes using the tight pestle.
v. Incubate for 30 minutes on ice and centrifuge lysates at 16,000 × g for 20 minutes at 4°C.
vi. Collect supernatant containing the enriched nuclear fraction and store at -80°C until use. Save ∼20µL for quantification (see Step (vii)).
vii. Quantify lysates for total protein content using BCA assay according to the manufacturer’s instructions.
B. Electrophoresis, semi-dry blotting and detection.

i. Denature 10 µg of HeLa Kyoto nuclear cell lysates or 15 µg of U-2 OS whole extracts in 1x NuPAGE® LDS sample buffer supplemented with 357.5 mM β-mercaptoethanol at 70°C, for 10 minutes.
ii. Perform electrophoresis of HeLa nuclear cell lysates in 4-12% Bis-Tris and of U-2 OS whole cell lysates in 3-8% Tris-Acetate pre-casted gels using either MOPS or Tris-Acetate running buffers, respectively.
iii. After the electrophoresis run is complete (∼1 hour), transfer gels onto 0.2 µm PVDF membranes using the mixed- (1.3 mA, 25 V, 7 minutes) or high-molecular weight (1.3 mA, 25 V, 10 minutes) transfer programs using a Trans-Blot® Turbo™ semi-dry transfer system (Biorad, 690BR023187).
iv. Quickly wash PVDF filters in dH2O and block overnight at 4°C in PBST (D-PBS + 0.1% v/v Tween 20) supplemented with 5% w/v BSA (for HeLa Kyoto nuclear extracts) or 1 hour at RT in PBST with 10% w/v non-fat dry milk (for U-2 OS whole cell lysates).
v. Incubate HeLa Kyoto PVDF filters with mouse anti-Nup93 (1:1000 in PBST/5% w/v BSA) and U-2 OS membranes with rabbit anti-SNAP primary antibodies (1:1000 in PBST/10% w/v dry milk); incubate overnight at 4°C.
vi. Wash three times with PBST and incubate membranes for ∼1 hour at RT with 1:5000 HRP-conjugated anti-mouse or anti-rabbit secondary antibodies; detect using the ECL system according to the manufacturer’s instructions.

**10B|** *Capillary electrophoresis.* For capillary electrophoresis, we have used the Jess device (ProteinSimple) using 66-440 kDa Jess Separation Module, 25 capillary cartridges.

A. Preparation of U-2 OS whole cell lysates.

i. U-2 OS whole-cell lysates were prepared, starting from confluent 10 cm Petri dishes and all steps were performed rigorously on ice.
ii. Wash confluent 10-cm Petri dishes three times with ice-cold D-PBS before adding 0.5 mL/plate of ice-cold RIPA lysis buffer supplemented with protease inhibitors and 1mM PMSF just before use.
iii. Scrape off cell monolayers from plates and collect in pre-chilled 1.5-mL Eppendorf tubes.
iv. Disrupt cell membranes by performing two freeze-thaw cycles using liquid nitrogen.
v. Centrifuge lysates at 16000 × g for 10 minutes at 4°C to pellet cell debris; collect supernatants in new Eppendorf tubes and store at -80°C until use.
B. Preparation of plates for capillary electrophoresis.

i. Prepare standard pack 3 reagents

a. Pierce foil of DTT, add 40µL of nuclease-free H2O to make a 400 mM solution.
b. Pierce foil of 5x Master Mix, add 20µL of 10x Sample Buffer and 20µL of prepared 400 mM DTT solution.
c. Pierce foil of Ladder, add 20µL of nuclease-free H2O.
d. Dilute Sample Buffer provided at 10x concentration in nuclease-free H2O to make the 0.1x Sample Buffer.
e. The optimal protein concentration depends on the expression level of the POI.
f. Dilute lysates 1:40 in 0.1x Sample Buffer.
g. Add 5x Fluorescent Master Mix to the diluted lysate (final dilution of lysate 1:50).
h. Gently mix by pipetting and close the tube.
i. Denature lysates:
j. Vortex and incubate at 95°C for 5 minutes.
k. Vortex and briefly spin down the denatured lysates.
l. Store on ice until use.
m. Prepare Luminol and Peroxide 1:1 v/v solution and gently mix; store on ice.
n. Dilute primary antibodies 1:50 in AB diluent, store on ice
o. . Dispense reagents into the assay plate using the volumes shown in the plate diagram, except washing buffer into the simple western well plate.
p. Centrifuge the plate for 5 minutes at ∼1000 x g at RT. Ensure liquid is fully down in all wells.
q. Add washing buffer to the desired wells.
r. Select the desired assay parameters (Supplementary Table 8) in Compass software.
s. Open Jess’s door and insert a 25 capillary cartridge into the holder. The interior light will change from orange to blue.
t. Remove the assay plate lid. Hold the plate firmly on the bench and carefully peel-off the evaporation seal. Pop any bubbles observed in the Separation Matrix wells with a pipette tip.
u. Place the assay plate on the plate holder, close Jess’s or Wes’s door and click the Start button in Compass software.
v. When the run is complete, discard the plate and cartridge.

Determination of genomic DNA concentration (Supplementary Methods 1)

(A) See manufacturer’s instructions

a. Calculate the genomic DNA concentrations using the standard curve (Supplementary Table 6).

## Southern Blot ● TIMING 14d

**11|** Southern Blot.

A. Design of Southern Blot probes

i. Ensure that the Southern blotting probe for the GOI is 100–500 bp. GC content should not exceed 40%.
ii. Ensure that this probe binds outside of the homology arms.
iii. In addition, design a probe against the CRISPR edited tag.
B. Cloning of probe-template DNA into backbone plasmid

i. Isolation of genomic DNA.

a. Pellet ∼4x10^6 cells and wash the pellet once with D-PBS. Remove excess D-PBS and leave ∼50µL residual D-PBS.

▪ **PAUSE POINT** Freeze pellets at -80°C.

b. Thaw cells at RT and tap the tubes gently to lose cellular pellets.
c. Mix 600μl of Nuclei Lysis Solution with 1:200 v/v RNase Solution.
d. Add 600μl Lysis Solution to thawed pellets (Step #3) by manual shaking until no visible clumps remain (∼10sec).
e. Incubate the mixture for 30 minutes at 37°C with shaking in a Thermomixer (Eppendorf).

▲ **NOTE** Allow samples to cool to room temperature for 5 minutes before proceeding.

f. Add 200μl of Protein Precipitation Solution and mix vigorously for 20 seconds; chill samples on ice for 5 minutes.
g. Centrifuge for 5 minutes at 16,000×g, RT. Precipitated protein will form a tight white pellet.
h. Carefully transfer the supernatant containing the DNA without disturbing the protein pellet into a clean 1.5ml microcentrifuge tube.

▲ **NOTE** Some supernatant may remain in the original tube containing the protein pellet. Leave this residual liquid in the tube to avoid contaminating the DNA solution with the precipitated protein.

i. Add 600μl of RT isopropanol. Mix the solution by inversion until the white thread-like strands of DNA form a visible mass.
j. Centrifuge for 10 minutes at 16,000 × g at RT. The DNA will be visible as a small white pellet. Carefully decant the supernatant.
k. Wash the DNA by adding 600μl of RT 70% ethanol followed by centrifuging for 5 minutes at 16,000×g, RT.
l. Thoroughly aspirate the ethanol by pipetting. Note that gDNA pellets are very loose at this stage.
m. Allow pellets to air-dry for 5-10 minutes.
n. Add 70μl of Rehydration Solution to rehydrate the DNA by incubating the solution overnight at RT.

▪ **PAUSE POINT** Store gDNAs at 2–8°C until use.

(ii) PCR to generate insert for cloning

a. Thaw 5x HotStar HiFidelity PCR Buffer, primers and Q-Solution.

▲ **NOTE** Mix solutions thoroughly before use

a. b. Prepare PCR reactions according to table 1

▲ **NOTE** It is not necessary to keep PCR tubes on ice since HotStar HiFidelity DNA Polymerase is inactive at RT.

c. c. Aliquot the indicated volumes into sterile PCR tubes and add template DNA. Mix thoroughly and spin down the tubes.
d. d. Place the PCR tubes in the thermal cycler and start the cycling program (Table 3). After amplification, samples can be stored overnight at 2–8°C or at –20°C for longer storage.

iii. Agarose gel electrophoresis and purification of PCR amplicon

a. Prepare a 1.5% (w/v) agarose gel using 1× TBE and 1:10,000 v/v SYBR-safe and equilibrate it in 1x TBE electrophoresis buffer.
b. Add 10μL of 6× loading dye to each PCR tube and load samples into the agarose gel; include one well with 10μL of Gene Ruler as marker.
c. Run electrophoresis at 120 V, 60 min.
d. Check the size of the PCR products and excise the band of interest from the gel using a scalpel. Transfer the gel-slice containing the PCR product into a 1.5 mL Eppendorf tube.
e. Purify DNA using MinElute Gel extraction kit (QIagen) according to manufacturer’s spin protocol instructions.
f. Elute DNA with 15µL nuclease-free d H2O

iv. Blunt end ligation of PCR amplicon in pCR-II-blunt-TOPO and transformation

a. Set up reactions according to Table 4. Mix well and incubate for 5 minutes at RT.
b. Store the reaction on ice. Meanwhile, prepare transformation reactions.

**Table 3:** PCR cycling conditions to generate insert for southern blot probe template

| cycle | step | Time | Temperature [°C] |
| --- | --- | --- | --- |
| 1 | initial activation step | 5 min | 95 |
| 40 | denaturation | 15 sec | 94 |
|  | annealing | 1 min | See Supplementary Table 4 |
|  | extension | 1 min | 72 |
| 1 | final extension | 10 min | 72 |
| 1 | end | indefinite | 4 |

**Table 4:** Blunt end ligation of PCR amplicon in pCR-II-blunt-TOPO

| Reagent | Volume [μL] |
| --- | --- |
| purified PCR product | 0.5–4 |
| Salt Solution | 1 |
| nuclease-free H<sub>2</sub>O | up to 5 |
| pCR™II-Blunt-TOPO® | 1 |

▴ **NOTE** You may store the TOPO® Cloning reaction at −20°C overnight.

c. Thaw a tube of NEB 5-alpha Competent E. coli cells on ice for 10 minutes.
d. Add 2 μL of the TOPO® Cloning reaction into a vial of One Shot® chemically competent E. coli.
e. Incubate on ice for 20–30 minutes.
f. Heat-shock the cells for 30 seconds at 42°C without shaking.
g. Immediately transfer the tubes into ice.
h. Add 950 μL of RT SOC medium. Cap the tube tightly and shake the tube horizontally (200 rpm) at 37°C for 1 hour.
i. Spread 100-200 μL from each transformation on a pre-warmed kanamycin-containing selection Agar plate; incubate overnight at 37°C.

v Plasmid isolation

a. Inoculate 2-4 tubes containing 5 mL LB-medium supplemented with 1 µg/mL kanamycin with single colonies under sterile conditions.
b. Incubate inoculated tubes overnight at 37°C with vigorous shaking.
c. The day next, pellet bacteria at 3000 x g, 5 min, 4°C.
d. Decant supernatant and isolate plasmid DNA using QIAprep® Miniprep kit (QIagen) according to the manufacturer’s instructions.

vi. Sanger sequencing

a. Prepare 20µL of 30-100 ng/µL purified plasmid and 10µL of 10 µM sequencing Primer (M13-RP 5’-CAGGAAACAGCTATGAC-3’) for sequencing
b. Send as Predefined Service to GENEWIZ.
c. Check sequences for correctness.

(C) DIG-labelling of Southern Blot probes.

(ii) Blunt end ligation of PCR amplicon in pCR-II-blunt-TOPO and transformation

a. PCR reaction setup and thermocycling

i. Thaw reagents and keep on ice. Briefly vortex and centrifuge all reagents before setting up reactions (Supplementary Table 3)
  i. Mix and spin down reactions. Transfer PCR tubes in the thermal cycler and start the cycling program (Table 5).
  (iii) Quality control of DIG labelling
    a. Prepare a 2% (w/v) agarose gel using 1× TBE and 1:10,000 SYBR-safe.
    b. Submerge the gel in electrophoresis tanks containing 1× TBE.
    c. Load 2µL of labelled and unlabelled PCR amplicon; include 10 μL of Gene Ruler as a marker.
    d. Run electrophoresis at 120 V, ∼60 min.
    e. Check at the transillumifnator the migration pattern. The unlabelled probe will run at the predicted size, whereas the DIG-labelled probe migrates slower.
    f. Store DIG-labelled PCR products at –20°C (up to one year).
  (D) Sample preparation, digestion and gel electrophoresis

i. Isolation of genomic DNA (see Section 10 (B) (i))
ii. Determination of genomic DNA concentration (Supplementary Methods 1)
iii. Digestion of genomic DNAs (gDNAs)
  a. Choose restriction enzymes that cut gDNA in fragments including the integration cassette and the annealing region of the probe. The electrophoretic shift of tagged vs. untagged sequences will be then maximized.
  ▲ **NOTE** Use restriction enzymes that are not sensitive to CpG methylation.
  b. Prepare digestions according to Table 6 and incubate overnight at 37°C. Digest between 2-5µg of genomic DNA per sample/lane.
  (iv) Gel electrophoresis of digested gDNAs
  (v) Prepare a 0.7% (w/v) agarose gel in 1× TBE. The thickness of the gel should not exceed 6 mm to allow the complete transfer of the digested gDNA fragments. Gels should be prepared just before use. Do not include EtBr in the gel, because it can cause uneven background if the gel is not running long enough.
  (vi) Submerge the gel in electrophoresis tanks containing 1× TBE.
  (vii) Mix 0.5µL of HindIII-digested, DIG-labelled λ-DNA with 1µL 6x loading dye and 4.5µL of nuclease-free H2O (marker).
  (vii) Add 8 μL of 6x loading dye to the digested gDNAs.
  (ix) Run the samples at ∼25-30 V overnight at 4°C
  (E) Blotting and crosslinking

(i) Trim off excess gel.
  ▲ **NOTE** Depurination is only necessary if the target DNA fragment is >5 kb.

(ii) Denature gel twice with Denaturation Solution, 15 minutes each. ▲ **NOTE** Use the minimal volume of solution required to fully cover the gel.

(iii) Rinse the gel briefly with nuclease-free dH2O. Neutralize the gel twice for 15 min with Neutralization Solution.
(iv) Equilibrate the gel for at least 10 min in 20x SSC.
(v) Place a piece of Whatman 3MM paper that has been soaked with 20× SSC above a «bridge» (e.g. a glass plate) that rests in a shallow reservoir of 20× SSC.
(vi) Place the gel over the soaked sheet of Whatman 3MM paper. Roll a sterile pipette over the sandwich to remove air bubbles formed between the gel and the paper.
(vii) Cut a piece of Positively Charged Nylon Membrane to the size of the gel.
(viii) Place the dry membrane on the DNA-containing surface of the gel. Use a pipette to eliminate air bubbles as before.
(ix) Complete the blot assembly by adding a dry sheet of Whatman 3MM paper, a stack of paper towels, a glass plate and a 200 – 500 g weight on top.
(x) Allow the blot to transfer overnight in 20× SSC, RT.
(xi) Remove the paper stacks and place the membrane-gel sandwich (DNA side facing up) on Whatman 3MM paper that has been soaked in 2x SSC.
(xii) Mark the wells with a pencil and remove the gel.
(xiii) Expose the wet membrane to UV light, using the UV Stratalinker auto-crosslinking protocol (120 mJ) to immobilize DNA.
(xiv) Rinse the membrane briefly in nuclease-free H2O.

**Table 5:** PCR cycling for DIG labelling of Southern Blot Probes

| cycle | step | Time | Temperature [°C] |
| --- | --- | --- | --- |
| 1 | initial activation step | 5 min | 95 |
| 30 | denaturation | 30 sec | 94 |
|  | annealing | 30 sec | See Supplementary Table 4 |
|  | extension | 40 sec | 72 |
| 1 | final extension | 7 min | 72 |
| 1 | end | indefinite | 4 |

**Table 6:**
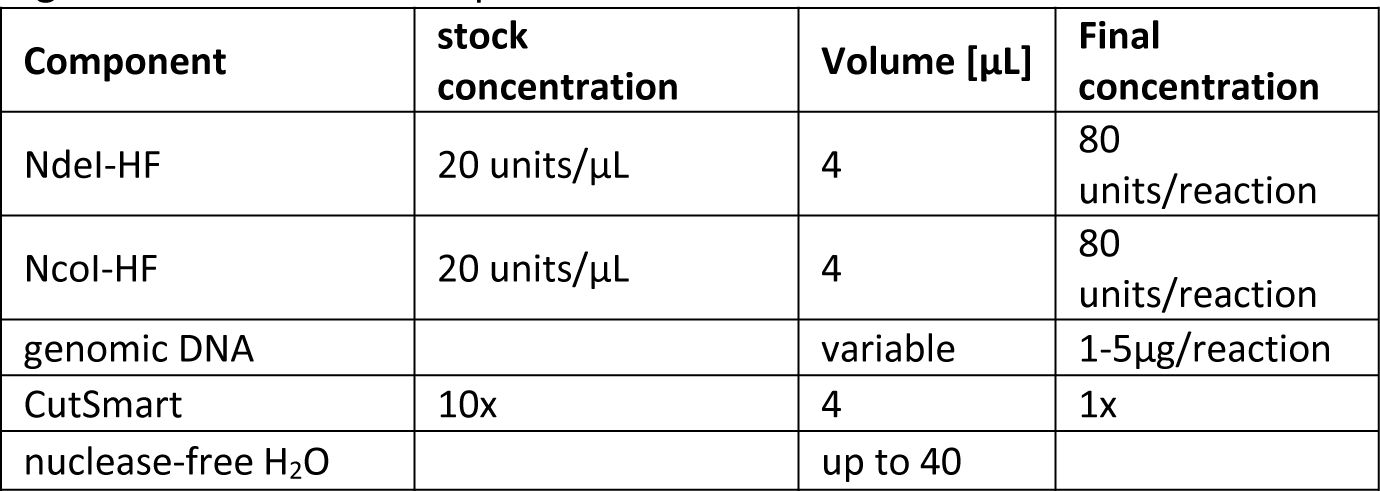
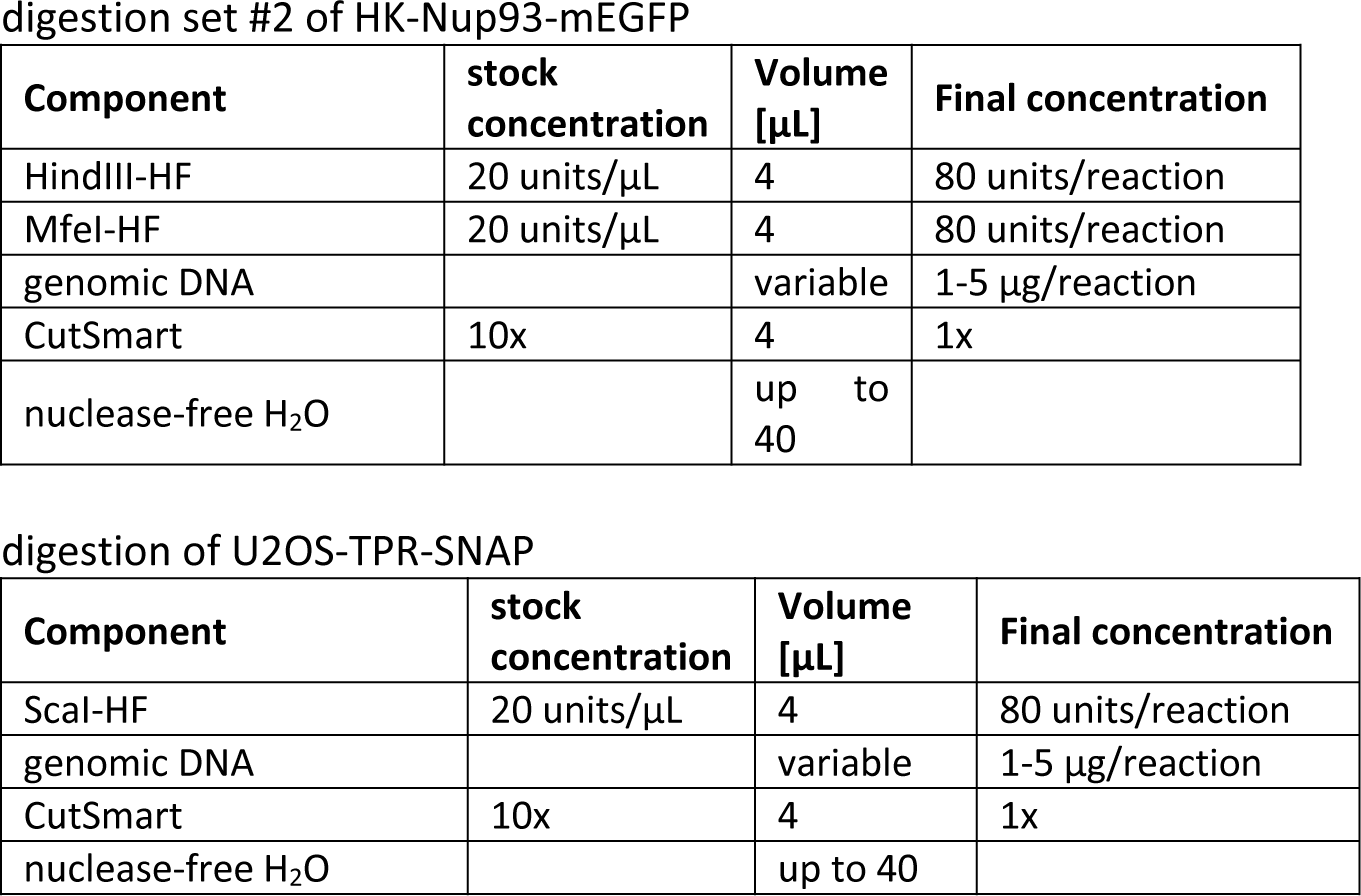
Restriction digestion setup for Southern Blotting

## ▪ **PAUSE POINT** store the membrane in 2x SSC at 4°C

(F) Pre-hybridization, hybridization and signal detection

▲ **NOTE** All hybridization steps were performed in hybridization bottles using a hybridization oven (Amerex) pre-set at the calculated hybridization temperatures of the specific probe used (Table 7). The volumes of solution required depend on the size of the membrane and of the hybridization bottles. Membranes need to be covered. In our case, we used 35 mL bottles. In case the membrane is too large to fit into the bottle and overlaps itself, we recommend overlying the membrane with a nylon mesh exceeding 0.5cm the membrane size. The mesh was pre-equilibrated in 2× SSC buffer and rolled up with the wet membrane before inserting them together into the hybridization bottle.

i. Incubate the membrane for at least 30 minutes at the correct hybridization temperature in Hybridization Solution. Agitate the membrane gently during this pre-hybridization step. Meanwhile, thaw the frozen Southern Blot probe.
ii. Use ∼2µL of DIG-labelled probe/mL and dilute it in nuclease-free H2O up to 100µL.
iii. Denature the diluted Southern Blot probe by boiling it in H2O for 5 min.
iv. Chill the probe quickly on an ice bath and spin down the tube briefly.
v. Immediately transfer the denatured probe to pre-warmed Hybridization Solution (Step (i).
vi. Discard the pre-hybridization solution from the bottle and replace it with the hybridization solution containing the denatured DIG-labelled probe (Step (v)).
vii. Incubate overnight at the desired temperature.

**Table 7:** Hybridization temperatures used for Southern Blotting

| probe | GC [%] | Length [bp] | T<sub>m</sub> [°C] | T hyb max [°C] | T hyb min [°C] | chosen hyb temp for SB [°C] |
| --- | --- | --- | --- | --- | --- | --- |
| TPR-Cterm-sbp1 | 39 | 233 | 63 | 43 | 38 | 42 |
| Nup93-Cterm-sbp2 | 41 | 206 | 64 | 44 | 39 | 43 |
| mEGFP-sbp | 57 | 331 | 71 | 51 | 46 | 51 |
| SNAP-sbp | 56 | 111 | 67 | 47 | 42 | 45 |

**Table 8:** dPCR setup and cycling conditions (STILLA)

| component | stock conc. | final conc. | volume [μL] |
| --- | --- | --- | --- |
| Perfecta Multiplex qScript ToughMix _ 5x | 5x | 1X | 5 |
| Fluorescein | 1 μM | 100 nM | 2.5 |
| Probe and primer Mix 12.5X | 12.5x | 1X | 2 |
| HindIII-HF | (100u/μL) | 0.4u/μL | 0.1 |
| DNA | variable |  | 12 |
| Nuclease-free H<sub>2</sub>O |  |  | Up to 25 |

12.5x probe and primer mix – U2OS-TPR-SNAP (allSNAP assay)
| component | ID | stock conc. [μM] | final conc. [μM] |
| --- | --- | --- | --- |
| target primer f | mito_821 | 100 | 12.5 |
| target primer r | mito_822 | 100 | 12.5 |
| target probe | ddPCR6-allSNAP_HEX | 100 | 3.125 |
| ref primer f | mito_823 | 100 | 12.5 |
| ref primer r | mito_824 | 100 | 12.5 |
| ref probe | ddPCR7-Ref_Cy5 | 100 | 3.125 |

Thermo cycling conditions – U2OS-TPR-SNAP (allSNAP assay)
| Temperature [$^{\circ}$C] | Time | No. of cycles |
| --- | --- | --- |
| 40 | default | 1 |
| 95 | 10 min | 1 |
| 94 | 30sec | 45 |
| 61 | 1min |  |

12.5x probe and primer mix – HK-Nup93-mEGFP (allGFP assay)
| component | ID | stock conc. [μM] | final conc. [μM] |
| --- | --- | --- | --- |
| target primer f | #808 | 100 | 12.5 |
| target primer r | #809 | 100 | 12.5 |
| target probe | dPCR1-HEX | 100 | 3.125 |
| ref primer f | #801 | 100 | 12.5 |
| ref primer r | #802 | 100 | 12.5 |
| ref probe | dPCR3-Cy5 | 100 | 3.125 |

Thermo cycling conditions – HK-Nup93-mEGFP (allGFP assay)
| Temperature [$^{\circ}$C] | Time | No. of cycles | Step |
| --- | --- | --- | --- |
| 40 | default | 1 | Droplet generation |
| 95 | 10 min | 1 | Denaturation |
| 94 | 30 sec | 45 | Denaturation |
| 61 | 1 min |  | Annealing |

12.5x probe and primer mix – HK-Nup93-mEGFP (HDR assay)
| component | ID | stock conc. [μM] | final conc. [μM] |
| --- | --- | --- | --- |
| target primer f | #814 | 100 | 12.5 |
| target primer r | #809 | 100 | 12.5 |
| target probe | dPCR1-HEX | 100 | 3.125 |
| ref primer f | #801 | 100 | 12.5 |
| ref primer r | #815 | 100 | 12.5 |
| ref probe | dPCR3-Cy5 | 100 | 3.125 |

Thermo cycling conditions - HK-Nup93-mEGFP (HDR assay)
| Temperature [°C] | Time | No. of cycles | Step |
| --- | --- | --- | --- |
| 40 | default | 1 | Droplet generation |
| 95 | 10 min | 1 | Denaturation |
| 94 | 30 sec | 45 | Denaturation |
| 61 | 1 min |  | Annealing |
| 72 | 2 min |  | Elongation |

**Table 9:**
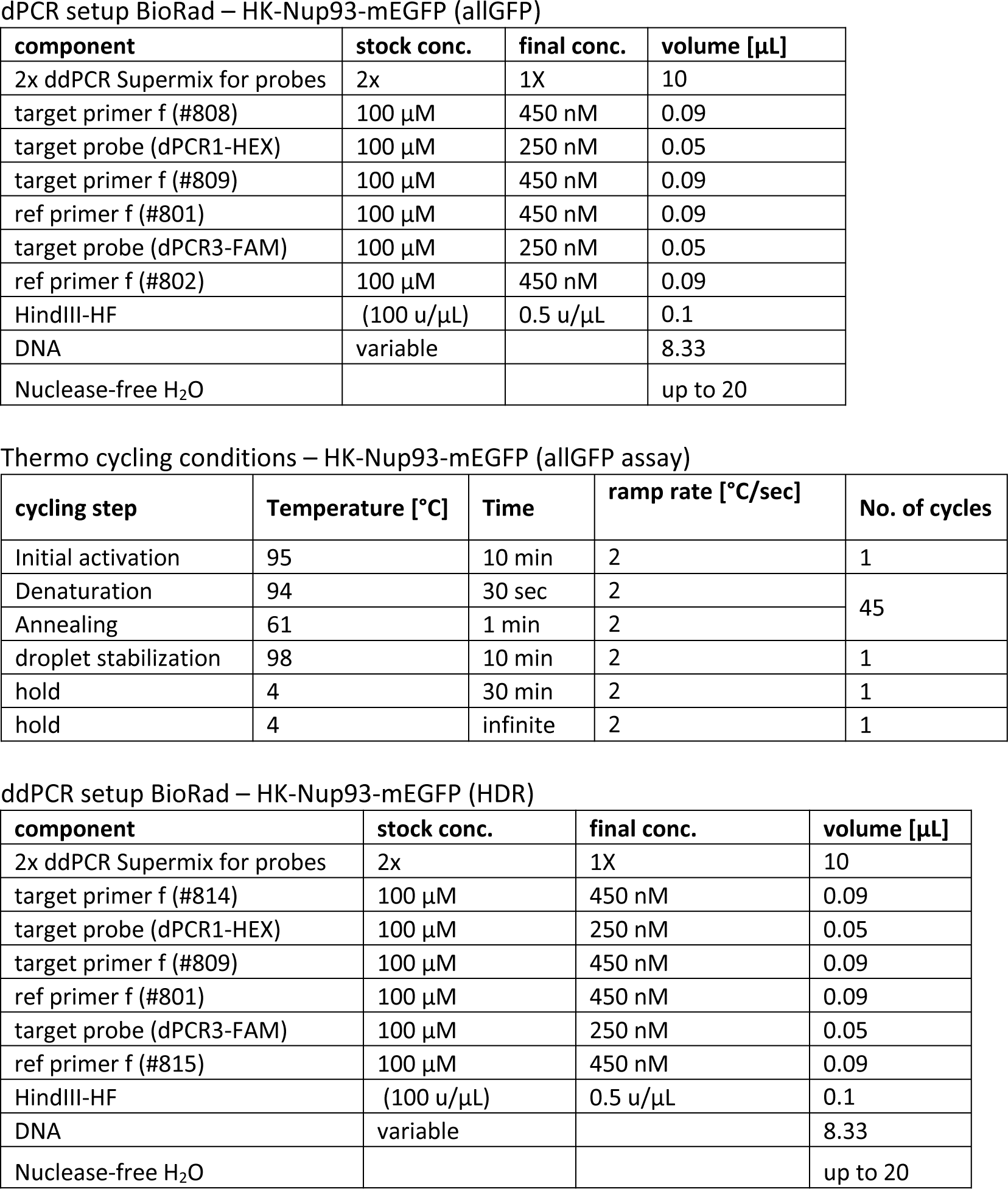
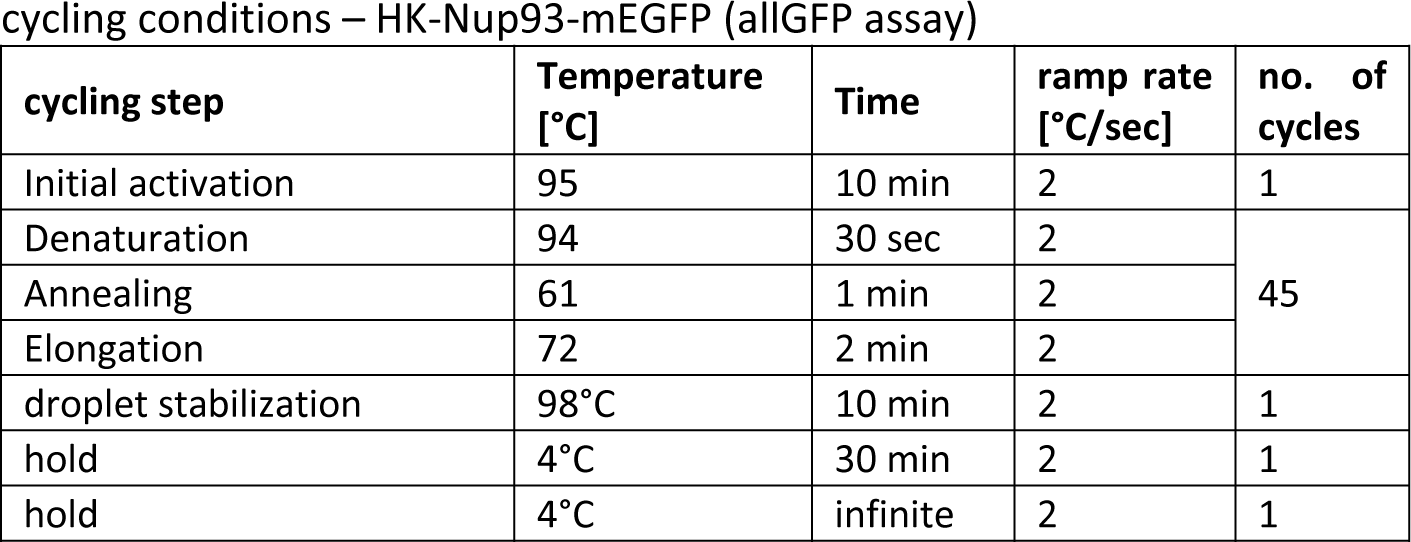
dPCR setup and cycling conditions (BioRad)

▴ **NOTE** In this and the following steps, the amount of buffer depends on the size of the tray used. For each step, be sure the membrane is completely covered with solution.

(viii) Open the hybridization bottle and immediately place the membrane in a plastic tray with the Low Stringency Buffer. Gently shake the blot during the washes.
(ix) Wash the blot twice for 15 min with pre-heated Low Stringency Buffer at RT.
(x) Meanwhile, pre-heat a suitable volume of High Stringency Buffer at 65°C. Add it immediately to the tray containing the blot and wash twice for 15 min at 65°C.
(xi) Transfer the membrane to a plastic container containing Washing Buffer. Incubate for 2 min at RT with shaking. Discard the Washing Buffer afterwards.
(xii) Add the Blocking Solution to the blot and incubate for 30 min, with shaking. Discard the Blocking Solution afterwards.

▲ **NOTE** This blocking step can last up to 3 hours without affecting results.

(xiii) Add the Antibody Solution to the blot and incubate the membrane for 30 min, with shaking.
(xiv) Discard the Antibody Solution. Wash membrane twice 15 min with Washing Buffer.
(xv) Equilibrate the membrane for 3 min in Detection Buffer.
(xvi) Place the membrane (DNA side facing up) inside a tightly sealed plastic envelope-like container.
(xvii) Add Ready-to-use-CDP-Star, dropwise, over the surface of the blot until the entire surface is evenly soaked.
(xviii) Immediately cover the dampened part of the membrane with the second side of the container so the substrate spreads evenly over the membrane area. Do not let air bubbles form between the membrane and the upper surface of the container.
(xix) Incubate membrane for 5 min, RT.
(xx) Drain the excess liquid out and seal the sides of the container close to the membrane. Expose the sealed envelope (containing the membrane) at RT to Lumi-Film X-ray film (15 – 25 min).

▲ **NOTE** Continuous exposure can be extended up to two days after the addition of substrate.

(G) Stripping and re-hybridization

(i) Rinse membrane thoroughly in nuclease-free H2O for 1 min
(ii) Wash membrane twice at 37°C in Stripping Buffer 2x 15 min
(iii) Rinse membrane for 5 min in 2x SSC
(iv) Store stripped membrane for later use in 2x SSC at 4°C or repeat the hybridization and detection procedure with a different DIG-labelled probe.

## SUPPLEMENTARY TABLES

**Supplementary Table 1:**
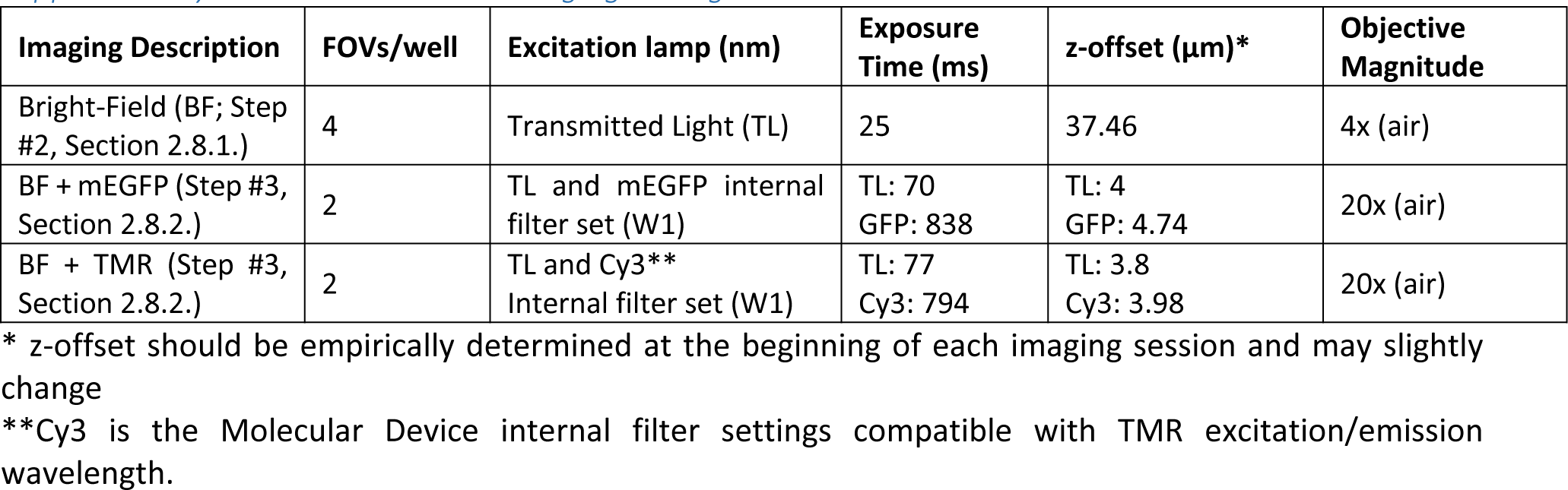
BF- and WF-imaging settings

**Supplementary Table 2:**
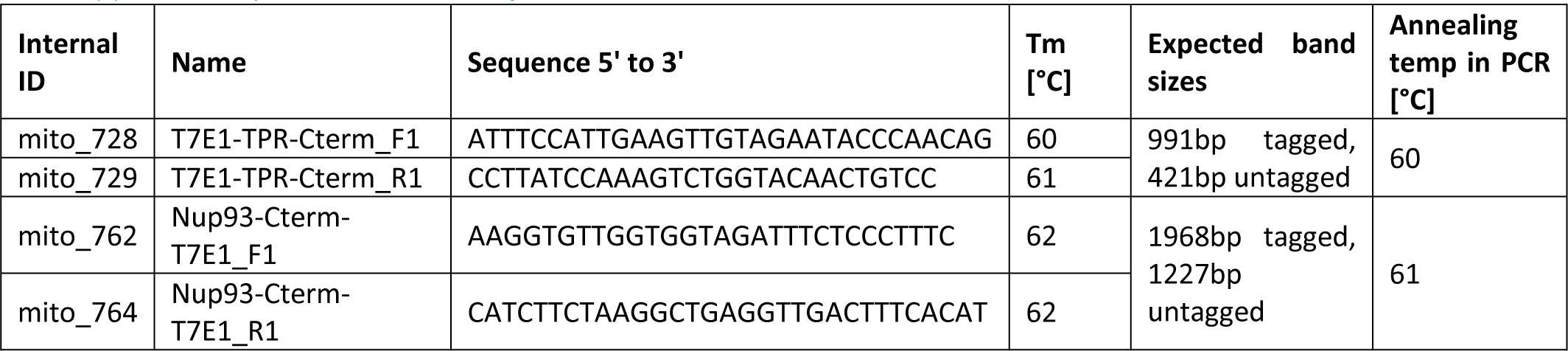
Primer info

**Supplementary Table 3:** DIG-labelling PCR setup of Southern Blot Probes

| Component | stock concentration | Volume [μL] | Final concentration |
| --- | --- | --- | --- |
| PCR buffer (vial 3) | 10x | 5 | 1x |
| PCR DIG Probe Synthesis Mix (vial 2) | 2 mM dATP, dCTP, 1.3 mM dTTP, 700 μM DIG-dUTP | 5 | 200 μM dATP, dCTP, 130 μM dTTP, 70 μM DIG-dUTP |
| dNTP stock solution | 2 mM each dNTP | 0 |  |
| Primer fwd | 10 μM | 5 | 1 μM |
| Primer rev | 10 μM | 5 | 1 μM |
| Enzyme mix (vial 1) |  | 0.75 |  |
| template plasmid DNA |  | variable | 10-100 pg/reaction |
| nuclease-free H<sub>2</sub>O |  | up to 50 |  |

Half labelling
| Component | stock concentration | Volume [μL] | Final concentration |
| --- | --- | --- | --- |
| PCR buffer (vial 3) | 10x | 5 | 1x |
| PCR DIG Probe Synthesis Mix (vial 2) | 2 mM dATP, dCTP, 1.3 mM dTTP, 700 μM DIG-dUTP | 2.5 | 200 μM dATP, dCTP, 165 μM dTTP, 35 μM DIG-dUTP |
| dNTP stock solution | 2 mM each dNTP | 2.5 |  |
| Primer fwd | 10 μM | 5 | 1 μM |
| Primer rev | 10 μM | 5 | 1 μM |
| Enzyme mix (vial 1) |  | 0.75 |  |
| template plasmid DNA |  | variable | 10-100 pg/reaction |
| nuclease-free H<sub>2</sub>O |  | up to 50 |  |

Unlabelled control
| Component | stock concentration | Volume [μL] | Final concentration |
| --- | --- | --- | --- |
| PCR buffer (vial 3) | 10x | 5 | 1x |
| PCR DIG Probe Synthesis Mix (vial 2) | 2 mM dATP, dCTP, 1.3 mM dTTP, 700 μM DIG-dUTP | 0 | 200 μM each dNTP |
| dNTP stock solution | 2 mM each dNTP | 5 |  |
| Primer fwd | 10 μM | 5 | 1 μM |
| Primer rev | 10 μM | 5 | 1 μM |
| Enzyme mix (vial 1) |  | 0.75 |  |
| template plasmid DNA |  | variable | 10-100 pg/reaction |
| nuclease-free H<sub>2</sub>O |  | up to 50 |  |

**Supplementary Table 4:**
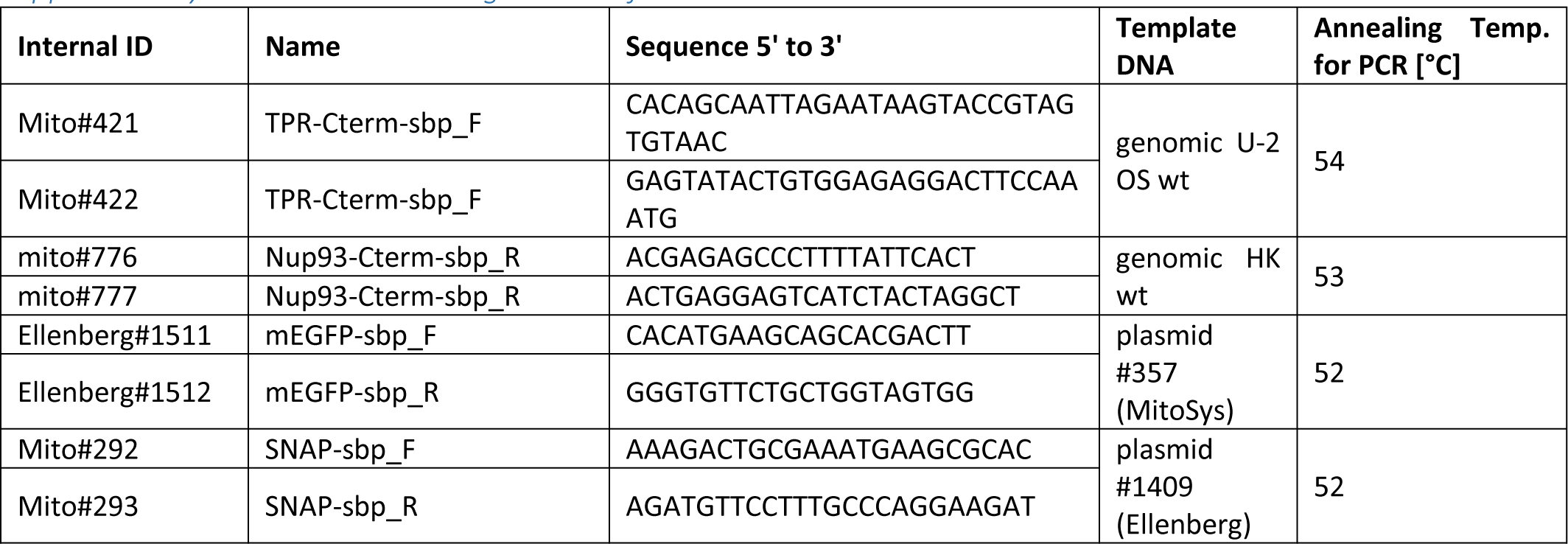
DIG-labelling Primer info

**Supplementary Table 5:** Pipetting scheme to prepare high-range standard curve for genomic DNA concentration determination

| Volume (μL) of TE | Volume (μL) of λ-DNA | Conc. in λ-DNA dilution | Total Volume (μL) |
| --- | --- | --- | --- |
| 0 | 500 of 2 μg/mL | 2000 μg/mL | 500 |
| 450 | 50 of 2 μg/mL | 200 ng/mL | 500 |
| 450 | 50 of 200 μg/mL | 20 ng/mL | 500 |
| 450 | 50 of 20 μg/mL | 2 ng/mL | 500 |
| 500 | 0 | blank | 500 |

**Supplementary Table 6:**
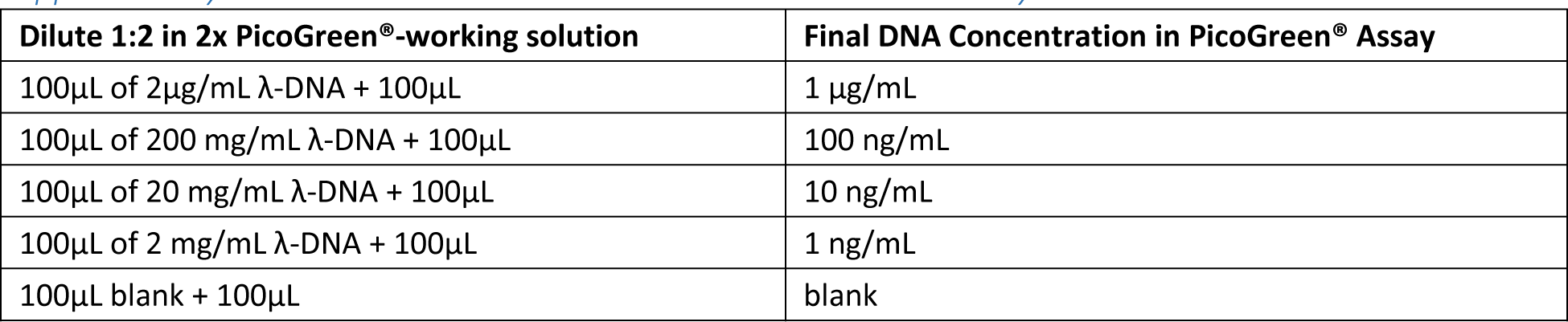
Final DNA concentrations in PicoGreen® Assay

**Supplementary Table 7:**
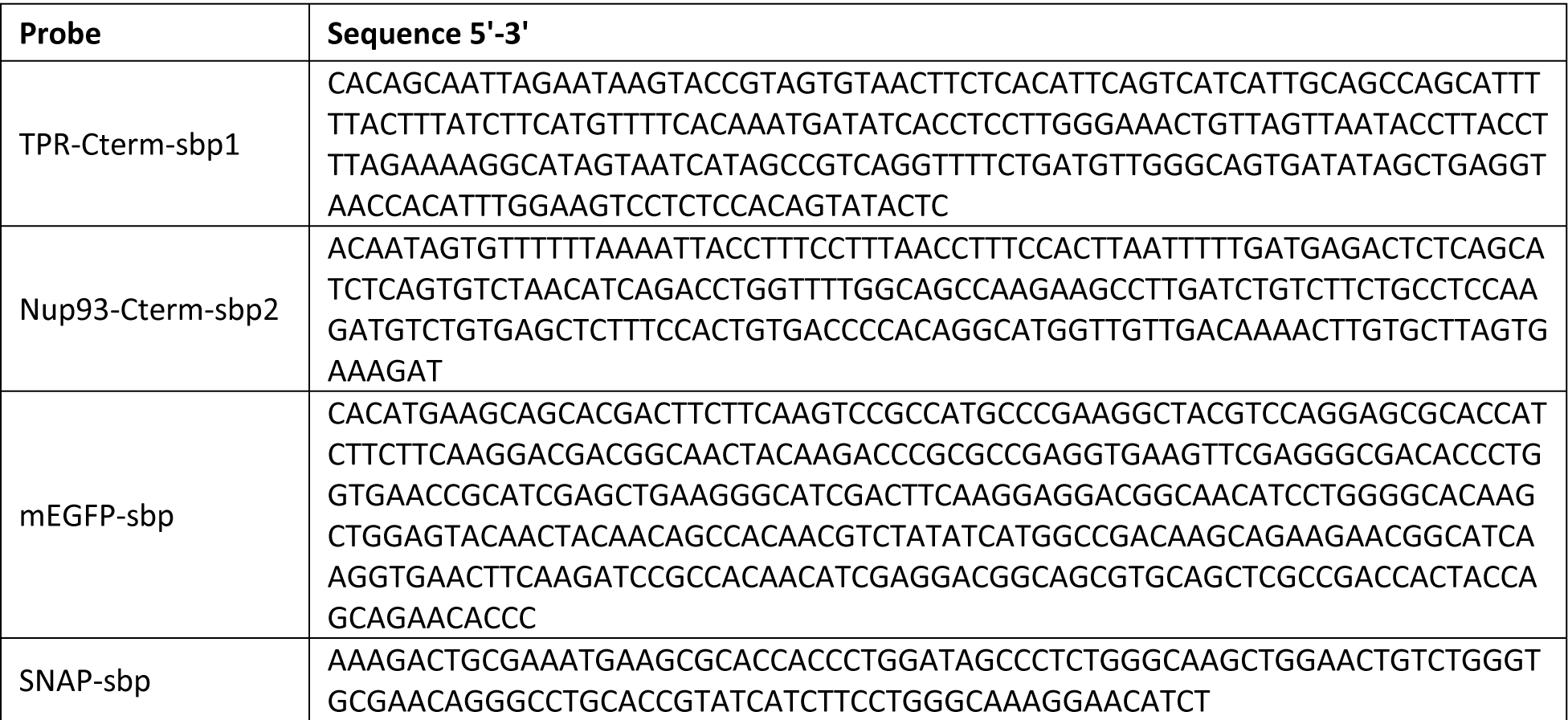
Sequences of Southern Blot Probes

**Supplementary Table 8:**
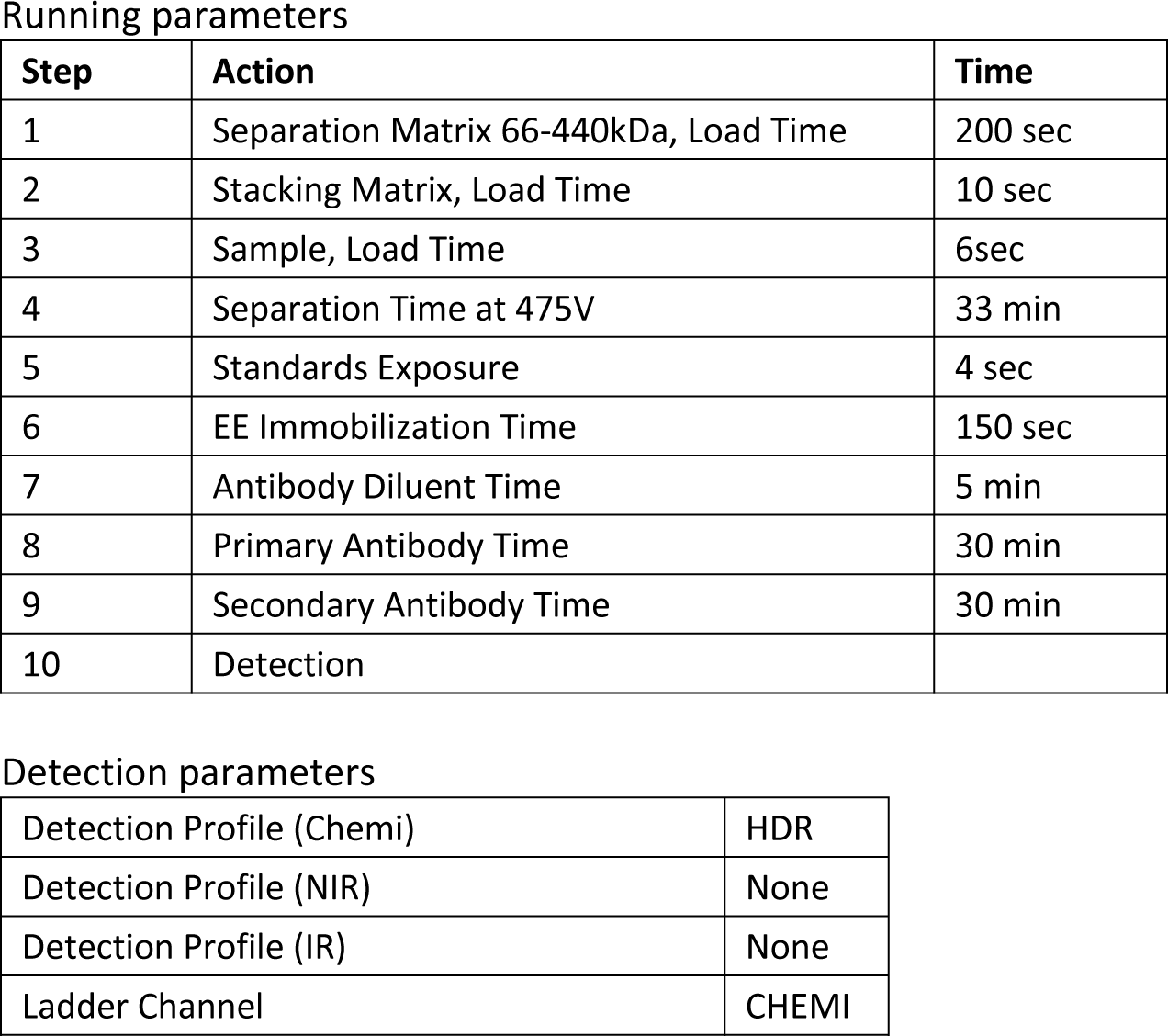
capillary electrophoresis parameters

**Running parameters**
| **Step** | **Action** | **Time** |
| --- | --- | --- |
| 1 | Separation Matrix 66-440kDa, Load Time | 200 sec |
| 2 | Stacking Matrix, Load Time | 10 sec |
| 3 | Sample, Load Time | 6sec |
| 4 | Separation Time at 475V | 33 min |
| 5 | Standards Exposure | 4 sec |
| 6 | EE Immobilization Time | 150 sec |
| 7 | Antibody Diluent Time | 5 min |
| 8 | Primary Antibody Time | 30 min |
| 9 | Secondary Antibody Time | 30 min |
| 10 | Detection |  |

**Detection parameters**
|  |  |
| --- | --- |
| Detection Profile (Chemi) | HDR |
| Detection Profile (NIR) | None |
| Detection Profile (IR) | None |
| Ladder Channel | CHEMI |

**Supplementary Table 9:**
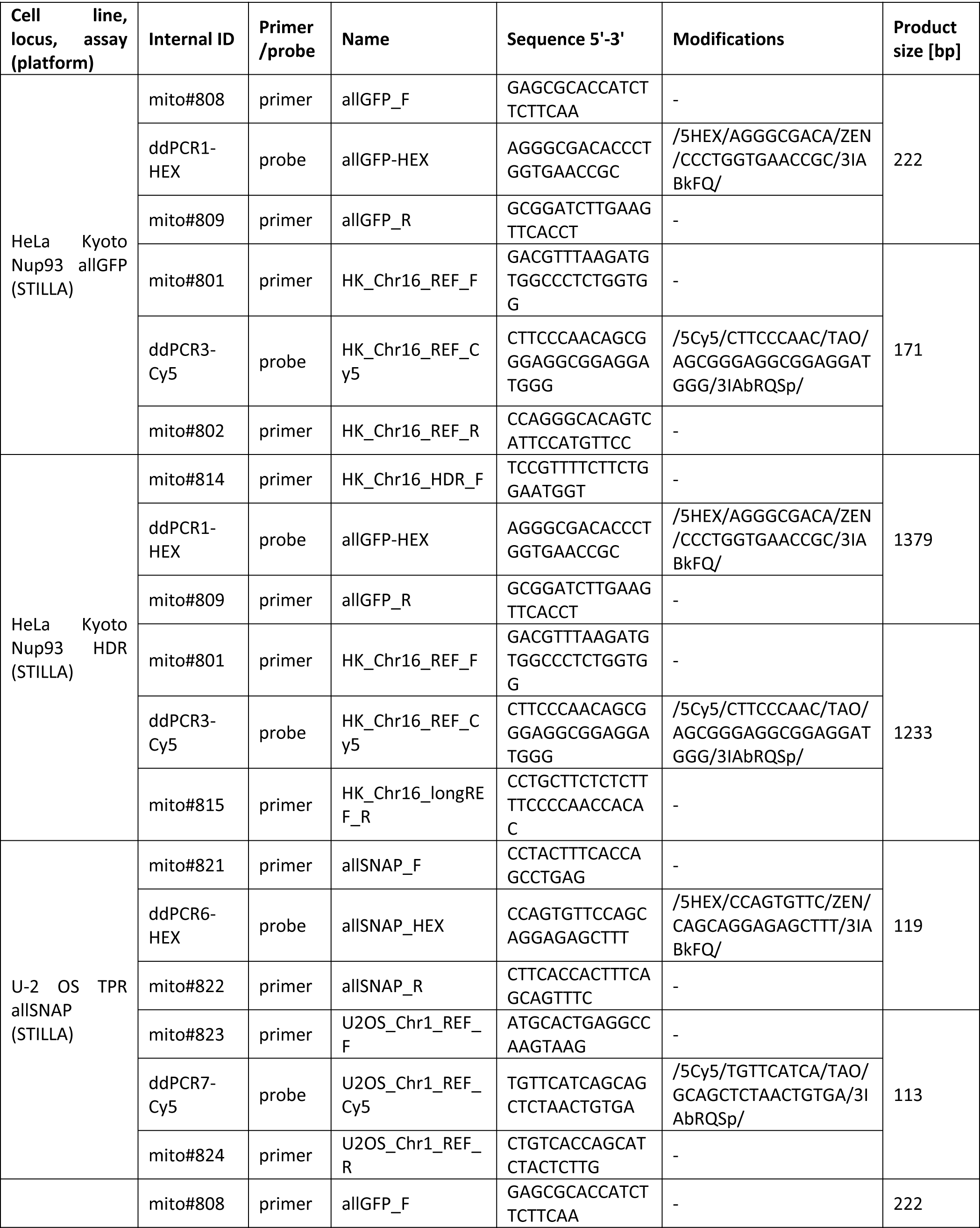
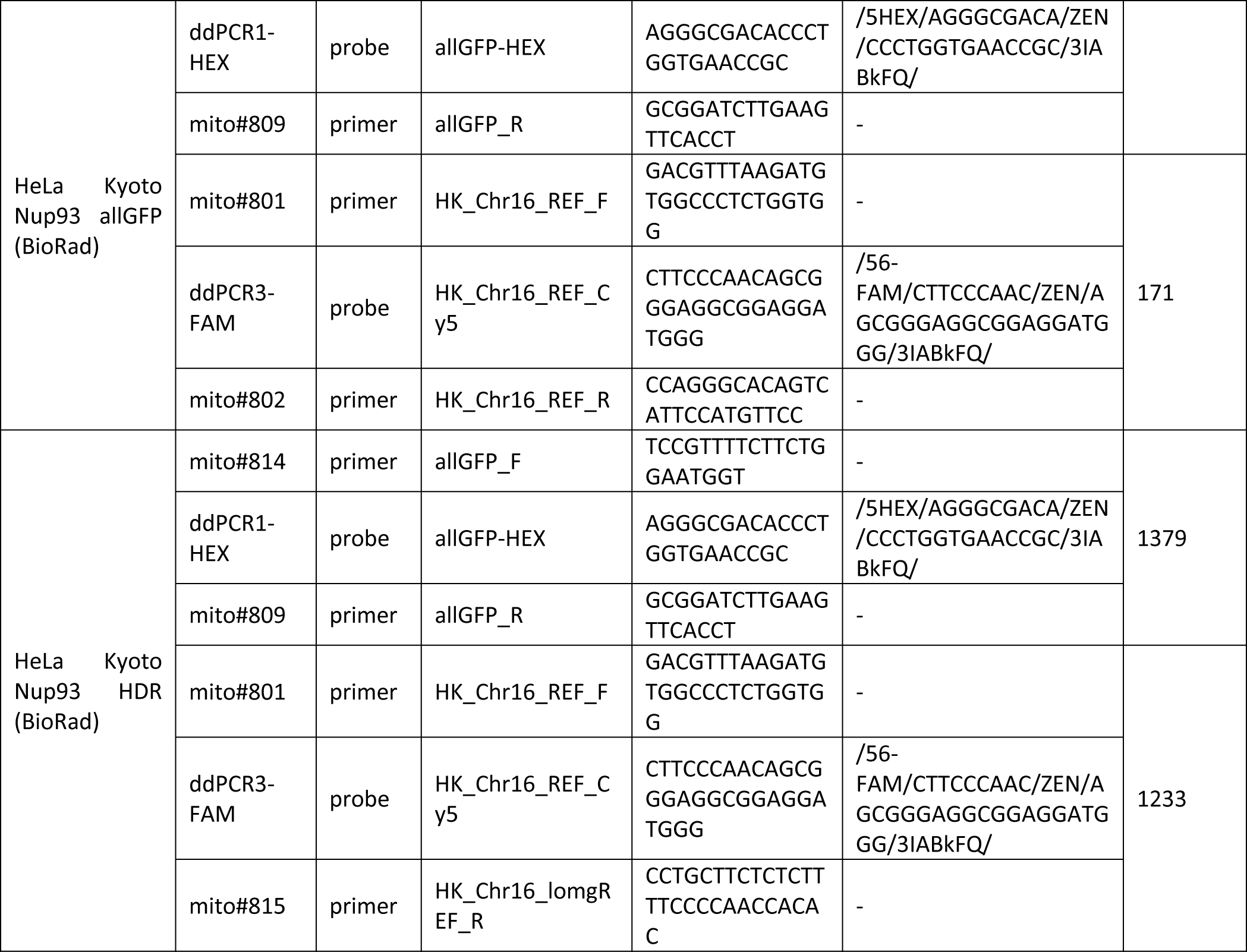
dPCR primer/probe sequences and product sizes

**Supplementary Table 10:** dPCR exposure times for the different detection channels (STILLA)

| Cell line and locus | Channel | Target name | Exposure time [ms] |
| --- | --- | --- | --- |
| HeLa Kyoto Nup93 allGFP | FAM - blue | Fluorescein | 150 |
|  | HEX - green | allGFP | 150 |
|  | Cy5 - red | REF_Ch16 | 50 |
| HeLa Kyoto Nup93 HDR | FAM - blue | Fluorescein | 150 |
|  | HEX - green | HDR | 200 |
|  | Cy5 - red | REF_Ch16 | 40 |
| U-2 OS TPR allSNAP | FAM - blue | Fluorescein | 100 |
|  | HEX - green | allSNAP | 250 |
|  | Cy5 - red | REF_Ch1 | 50 |

## SUPPLEMENTARY FIGURE CAPTIONS

**Supplementary Figure 1.**
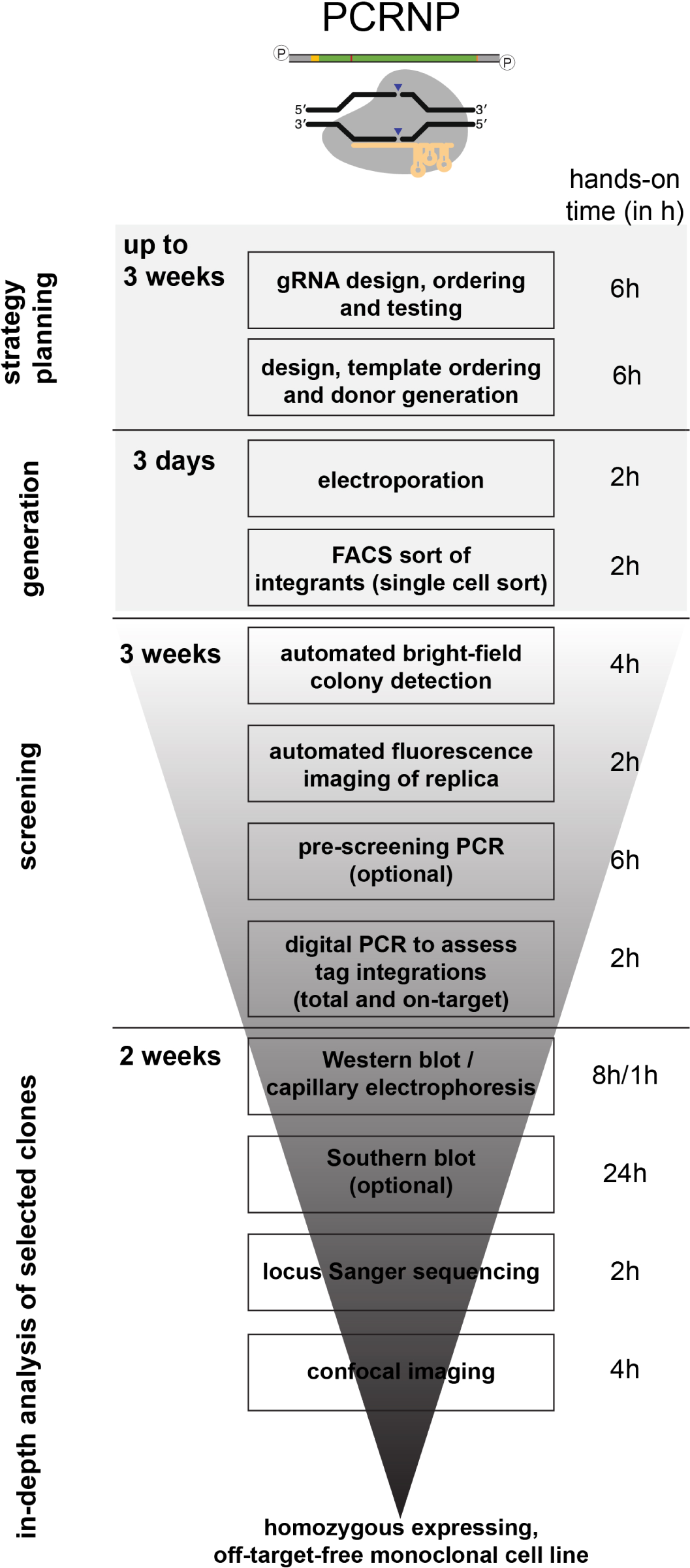
Generation and validation pipeline of CRISPR genome edited cell lines using the PCRNP method. After delivery of CRISPR components by electroporation, fluorescent edited cells were sorted via FACS into 96-well plates. Afterwards, monoclonal colonies were screened and selected for in-depth analysis. Notably, dPCR was implemented as a screening step to assess the number of tag integrations and select genotypes of interest to be propagated. The CRISPR pipeline lasts approximately 10 weeks. The hands-on times are indicated on the right side.

**Supplementary Figure 2.**
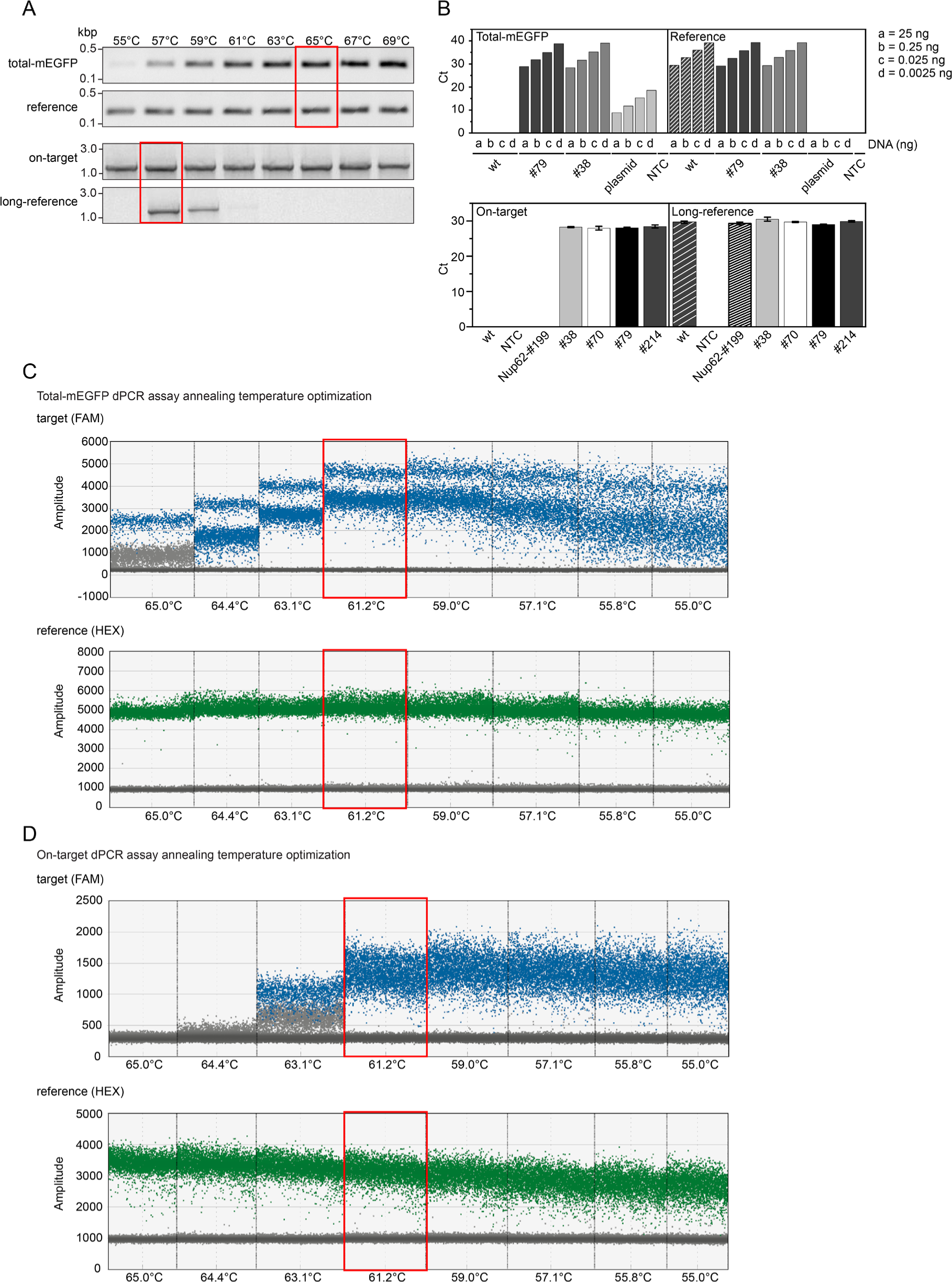
dPCR optimization. (A) Standard, temperature-gradient PCR of total-mEGFP and on-target assays including their respective reference assays. (B) qPCR threshold-Cycle (Ct) measurement for different genomic DNAs. Top panel: titration of genomic DNA using four different amounts (see legend) for the total-mEGFP assay and its reference; bottom panel: Ct-assessment for the on-target assay and its reference assay with different genomic DNAs. (C-D) 2-D dot-plots generated with QX200 (BioRad) to optimize the annealing temperatures of both assays.

**Supplementary Figure 3.**
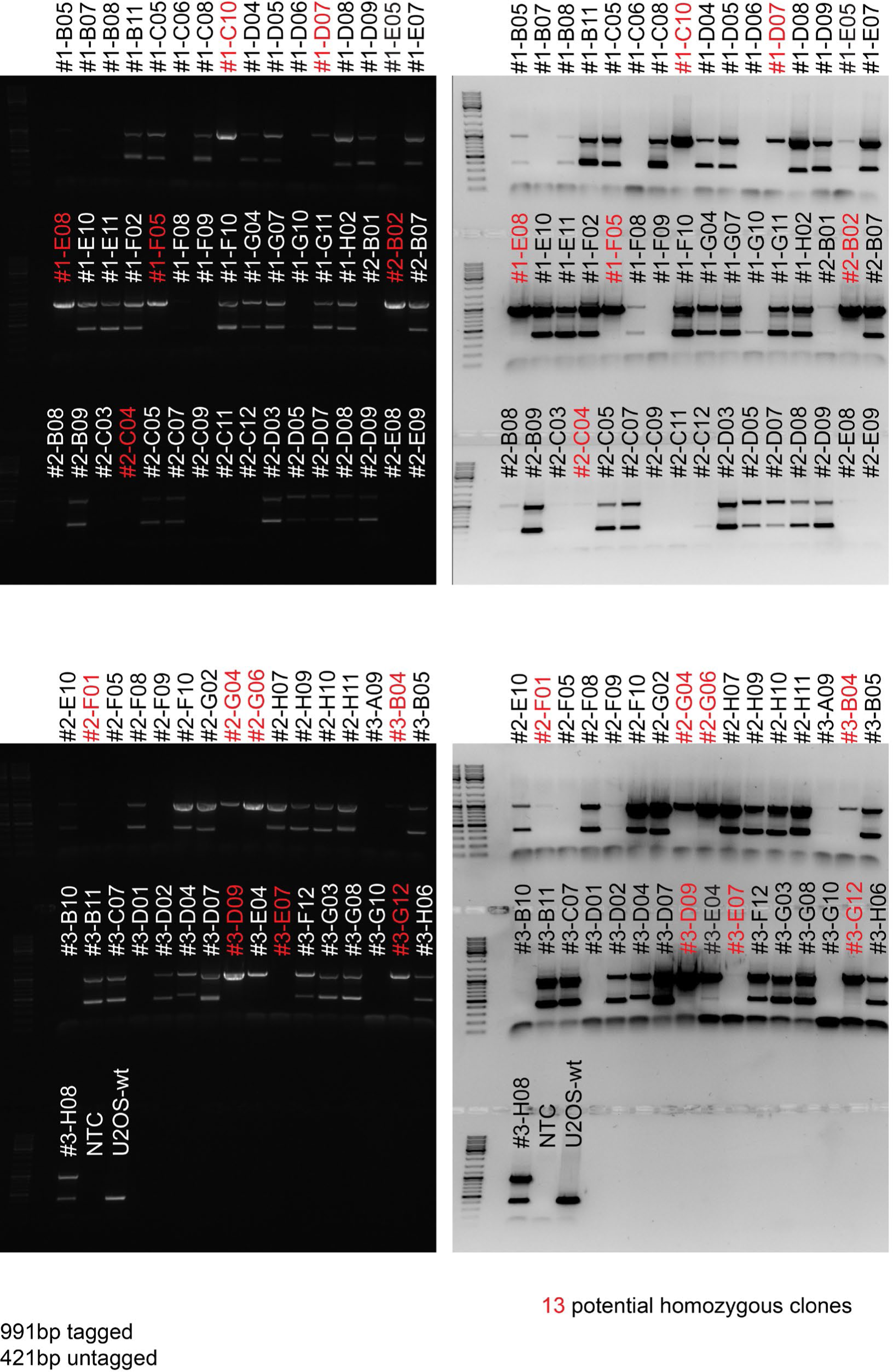
Standard PCR screening. SYBR SAFE (left panels) and inverted, black & white pictures (right panels) of electrophoretic gels containing amplified, genomic DNA extracted from expanded single colonies. Each clone is genetically homogeneous and displays either a single or double band corresponding to homozygous (indicated in red) or heterozygous genomes (indicated in black), respectively. Remarkably, the PCR illustrated in this figure has been carried out using primers annealing within the left- and right-homology regions (i.e. T7-primers). Expected size of a untagged allele 421bp, expected size of a tagged allele 991bp.

**Supplementary Figure 4.**
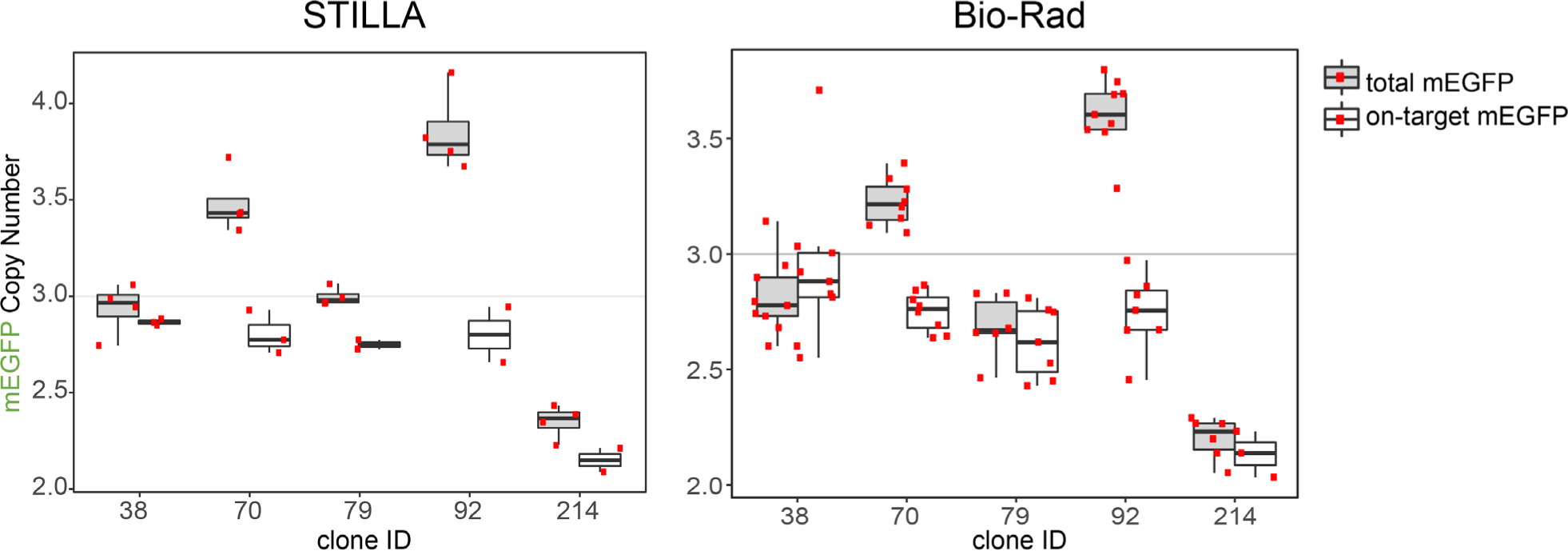
STILLA vs. Biorad dPCR performances. Boxplots illustrating copy-number analysis obtained from STILLA (left) and Biorad (right) dPCR devices. Data trend is comparable for both instruments across all tested clones. Note that despite the higher throughput offered by Biorad, (each individual red spot corresponds to an independent measurement), the dispersion of data is broader as compared to STILLA assessments.

**Supplementary Figure 5.**
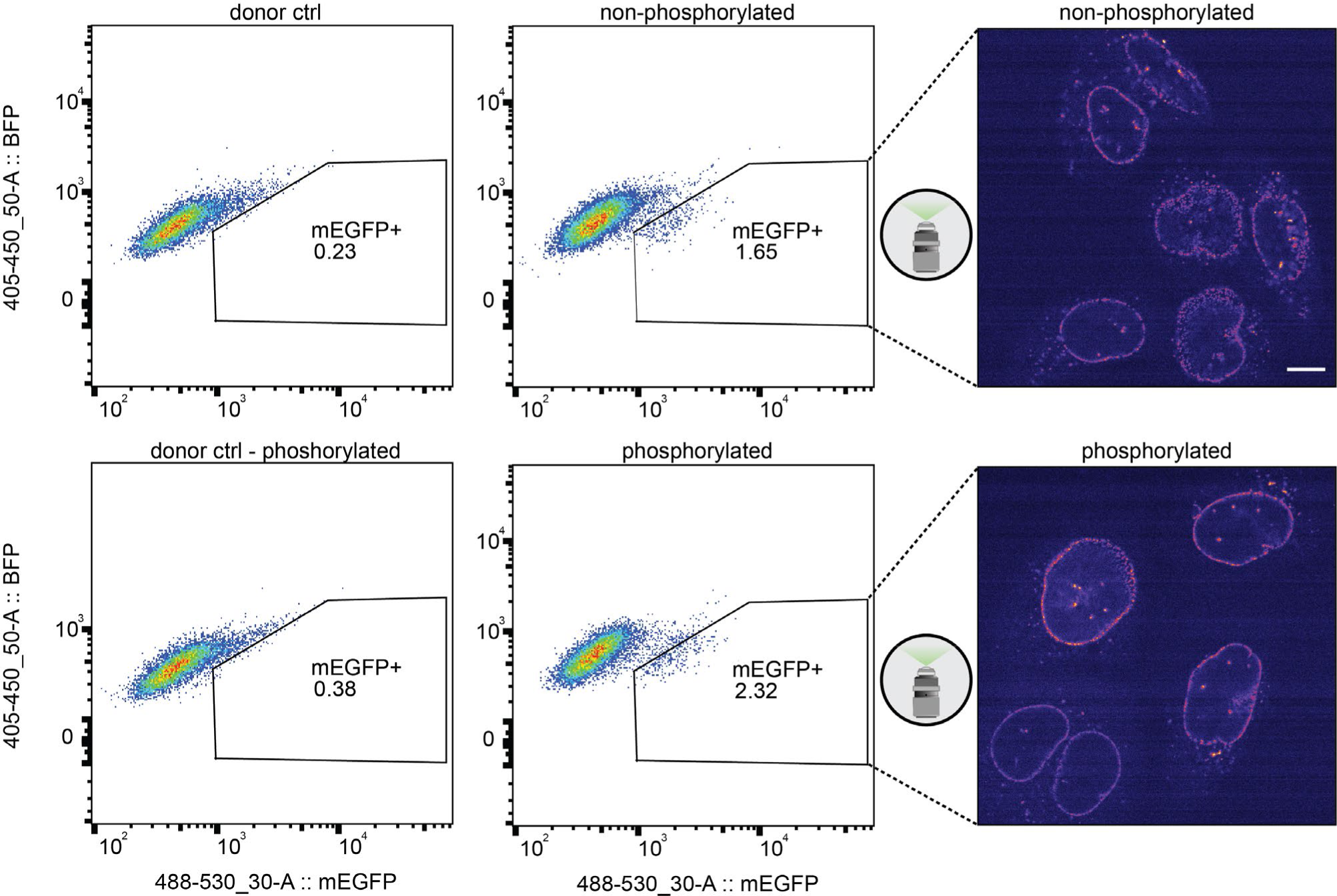
CRISPR efficiency in the presence of phosphorylated or non-phosphorylated DNA donor molecules. DNA fragments used as CRISPR donor molecules were PCR-amplified using either phosphorylated or non-phosphorylated primers and subsequently transfected into recipient cells. FACS sorting (left panel) reveals that higher tagging efficiencies with mEGFP were obtained using a phosphorylated PCR donor (bottom, mEGFP^+^ cells ∼2.3%) compared to a non-phosphorylated donor (upper panels, mEGFP^+^ cells ∼1.65%). In both cases, the tagged protein (Nup93) localizes in the nucleus outer rim, as determined by confocal imaging (right panel). PCR product with symmetric (40bp) homology arms [electroporation conditions - 1400V / 15ms / 2 pulses - 10µM sgRNA - Nup93-Cterm-as (Thermo), 10µM Cas9-HIFI-V3 (IDT), 4µM Electroporation enhancer (IDT), 1µM dsDNA-donor, 30nM HDR enhancer (IDT)]

